# Therapeutic suppression of *Tubb4a* rescues H-ABC leukodystrophy

**DOI:** 10.1101/2024.08.27.609903

**Authors:** Sunetra Sase, Julia L. Hacker, Prabhat R. Napit, Anjali Bhagavatula, Sarah Woidill, Annemarie D’Alessandro, Marisa A. Jeffries, Akshata Almad, Asako Takanohashi, Quasar Padiath, Judith Grinspan, Eric D. Marsh, Adeline Vanderver

## Abstract

Hypomyelination and atrophy of basal ganglia and cerebellum (H-ABC) is a rare leukodystrophy associated with causal variants in β-tubulin 4A (*TUBB4A*). The recurring variant p.Asp249Asn (D249N) presents in infancy with dystonia, communication deficits, and loss of ambulation during the first decade of life. In this study, we characterized a genetic murine series (*Tubb4a^KO/KO^*, *Tubb4a^D249N/+^*, *Tubb4a^D249N/KO,^* and *Tubb4a^D249N/D249N^*) to demonstrate that disease severity correlates with the expression of mutant Tubb4a and relative preservation of WT tubulin. To further evaluate the translational potential of *Tubb4a* suppression as a therapy in H-ABC, we identified a well-tolerated *Tubb4a*-targeted antisense oligonucleotide (ASO) candidate that selectively reduces Tubb4a. Notably, single intracerebroventricular (i.c.v.) administration of ASO in postnatal *Tubb4a^D249N/KO^* mice drastically extends its lifespan, improves motor phenotypes, and reduces seizures. Neuropathologically, treating ASO *Tubb4a^D249N/KO^* mice prevents myelin and oligodendrocyte loss and recovers visual evoked potential latencies *in vivo*. Furthermore, the microtubule function of *Mbp* mRNA transport from oligodendrocyte (OL) soma to myelin sheath is retained. A major limitation we noted is that ASOs fail to target cerebellar granule neurons even with multiple routes of administration in the brain. This is the first preclinical proof-of-concept for *Tubb4a* suppression via ASO as a disease-modifying therapy for H-ABC.

## Introduction

*TUBB4A*-related leukodystrophy is a rare neurodevelopmental disorder associated with monoallelic pathogenic variants in the *TUBB4A* gene (1). These variants cause a heterogeneous disease spectrum ranging from a severe early infantile encephalopathy, a late infantile presentation typified by progressive motor impairment, and a mild juvenile or adult-onset phenotype with prominent dystonia and spastic diplegia. The late infantile form is often characterized by distinctive neuroimaging features of hypomyelination and atrophy of the basal ganglia and cerebellum (H-ABC) (2, 3). H-ABC is also closely associated with a recurrent variant, p.Asp249Asn (D249N), present in over 20% of published individuals (4). H-ABC typically manifests in toddlers after normal development with progressive gait impairment and clinical features of spasticity, rigidity, dystonia, and cerebellar ataxia. Affected individuals experience feeding difficulties, seizures, loss of communication, and ambulation loss before the end of the first decade of life. No curative treatments are available for H-ABC; disease-modifying therapies are urgently needed (1–3, 5).

*TUBB4A* encodes the β-tubulin 4A protein, which is highly expressed in oligodendrocytes (OLs; myelin-producing cells) and specific central nervous system (CNS) neurons (6, 7). β-tubulins such as TUBB4A heterodimerize with the α-tubulin to form microtubules (MT), which are highly dynamic polymers essential for the extension of OL processes, myelin sheath elongation, and neuronal growth, morphology, and migration (8, 9). *TUBB4A* mutations disrupt MT dynamics and polymerization, which may burden critical myelin, neuronal protein, and organelle transport necessary for early neurodevelopment (10–12). H-ABC pathology demonstrates cell-autonomous deficits in OL lineage cells, striatal neurons (medium spiny neurons; MSNs), and cerebellar granule neurons (CGNs) (7, 13) corresponding to the characteristic clinical and radiologic manifestations of the disease.

To model H-ABC, different *Tubb4a-*associated rodent models have been reported (10, 14, 15). Our own published H-ABC mouse model with homozygous D249N variants (*Tubb4a^D249N/D249N^*) shows a severe phenotype with progressive ataxia, gait abnormalities, tremor, severe hypomyelination, and significant loss of OLs, CGNs, and striatal neurons (humane end-stage at postnatal day [P32-37]) (10). A study using a mouse model with a different *Tubb4a* variant, *Tubb4a* p.Asn414Lys mutation ([N414K]; *Tubb4a^N414K/+^*), also defined CGN and OL involvement as cell-autonomous (14). A recent study reported a *Tubb4a* knockout (*Tubb4a^KO/KO^*) mouse model with no behavioral or myelination deficits was attributed to compensation by other tubulin monomers in MT assembly (14). Studies of MT dynamics across published rodent and induced pluripotent stem cell (iPSC) models show that mutant *Tubb4a* causes toxic gain-of-function, disrupting MT polymerization and stability (10, 11, 14). These findings support the idea that reducing toxic Tubb4a may be a valuable therapeutic strategy in *TUBB4A*-related leukodystrophy.

Targeted antisense oligonucleotides (ASOs) that reduce messenger RNA (mRNA) levels are in clinical use for pediatric neurodegenerative diseases and provide promising clinical benefits (16, 17). Pre-clinical ASO treatments demonstrate outstanding rescue for other hypomyelinating leukodystrophies, such as Pelizaeus-Merzbacher disease (18). ASOs are short, single-stranded DNA molecules designed to degrade target mRNA and block translation of the encoded protein. ASO gapmer design chemistry is well-established and contains a phosphorothioate (PS) backbone with 2′-O-methoxyethyl (2′-MOE) and locked nucleic acid modifications (19). These ASO gapmers bind to target mRNA, resulting in RNase H1-mediated mRNA degradation.

The current study investigated the germline and therapeutic potential of reducing Tubb4a in our H-ABC mouse model. We characterized *Tubb4a^KO/KO^* and *Tubb4a^D249N/KO^*mice relative to wildtype (WT), *Tubb4a^D249N/+^*, and *Tubb4a^D249N/D249N^*mice. The compound heterozygote model, *Tubb4a^D249N/KO^*, displays severe deficits, albeit with slower disease progression relative to *Tubb4a^D249N/D249N^* mice. We also confirmed that germline loss of *Tubb4a* is well-tolerated (14). To test the therapeutic potential of *Tubb4a* downregulation, we developed, screened, and selected anti-*Tubb4a* ASO. We show that administration of a *Tubb4a*-targeted ASO via intracerebroventricular injection (i.c.v.) bolus into postnatal *Tubb4a^D249N/KO^* mice remarkably lengthened survival, improved behavioral performance, decreased seizures, and preserved myelin. A *Tubb4a* ASO-based therapy is the first molecular therapy for H-ABC and supports *Tubb4a* suppression approaches in clinical trials with potential applicability across the spectrum of *TUBB4A*-associated leukodystrophies.

## Results

### Germline suppression of *Tubb4a* and relative preservation of WT rescue D249N-associated motor phenotypes

To confirm whether *Tubb4a* suppression provides a therapeutic benefit in H-ABC, we bred H-ABC models *Tubb4a^D249N/+^* (10) and the *Tubb4a^KO/+^* mouse model (obtained from KOMP consortium) to create a phenotypic spectrum with variable dosage of Tubb4a mutant and WT. The *Tubb4a^D249N/+^* mouse model exhibits myelin deficits without behavioral abnormalities and can reproduce. The *Tubb4a^KO/+^* mice are phenotypically normal and breed well. Their progeny includes *Tubb4a^KO/KO^*, *Tubb4a^D249N/KO^* and *Tubb4a^D249N/D249N^* mice. As per published reports (14), *Tubb4a^KO/KO^* mice are viable and fertile, have no overt cage behavior abnormalities or differences in weaning compared with WT littermates, and have normal survival (**Figure 1B**). The homozygous *Tubb4a^D249N/D249N^*mice mirror the cellular, molecular, and neurologic phenotypes (myelin loss and neuronal atrophy) of classical H-ABC (10). They show severe tremors, ataxia, and seizures starting at P7-P10, reaching a humane end-stage at P32-P37. The compound heterozygous mouse model *Tubb4a^D249N/KO^*exhibits an extended lifespan, with a later onset of tremors and ataxia (P19-21) and a later onset of severe dystonia and seizures, approaching the humane end-stage only after several months (Mean survival of P108-P110). To determine the disease course, we conducted various motor neurobehavioral tests across all genotypes at different ages, extending to one year for less symptomatic genotypes. Additionally, we quantified tremors, a common disease phenotype, by measuring tremor amplitudes at multiple ages (**Figure 1C**). As expected, *Tubb4a^KO/KO^*and *Tubb4a^D249N/+^* mice show no tremors or motor impairment up to a year of life (**Figure 1C-F, Figure S1A-D**). Although *Tubb4a^D249N/D249N^* mice exhibit intense tremors at P21, the tremor amplitude significantly decreases at the humane end-stage (P32-37) due to limited locomotion (**Figure 1C**). In contrast, *Tubb4a^D249N/KO^* mice show only mild ataxia at P19-P21 and tremors at P30. This tremor progressively increases at P60 and P90 (**Figure 1C**). We tested mice on an accelerating rotarod and assessed grip strength across all genotypes, limited by survival (**Figure 1B**). While *Tubb4a^D249N/D249N^*mice exhibit poor rotarod and grip strength performance, *Tubb4a^D249N/KO^*mice initially perform like WT mice but progressively decline by P60 to P90 (**Figure 1D-F**). As *Tubb4a^D249N/D249N^* and *Tubb4a^D249N/KO^* mice approach their humane end-stage (P32-P37 and P108-110, respectively), they exhibit multiple spontaneous tonic-clonic seizures, characterized by hindlimb extension and tonic stiffening, consistent with clinical presentations of H-ABC-affected individuals exhibiting epilepsy. We counted seizures over a three-hour period and observed increased seizures in both models, with the most significant frequency in *Tubb4a^D249N/D249N^* mice. Notably, *Tubb4a^KO/KO^*and *Tubb4a^D249N/+^* do not exhibit any seizures (**Figure 1G**). Overall, motor impairment, survival, and seizures were proportional to the dosage of mutant *Tubb4a* relative to WT *Tubb4a* (*Tubb4a^D249N/D249N^ > Tubb4a^D249N/KO^ > Tubb4a^D249N/+^ > Tubb4a^KO/KO^*or WT mice).

**Figure 1:**
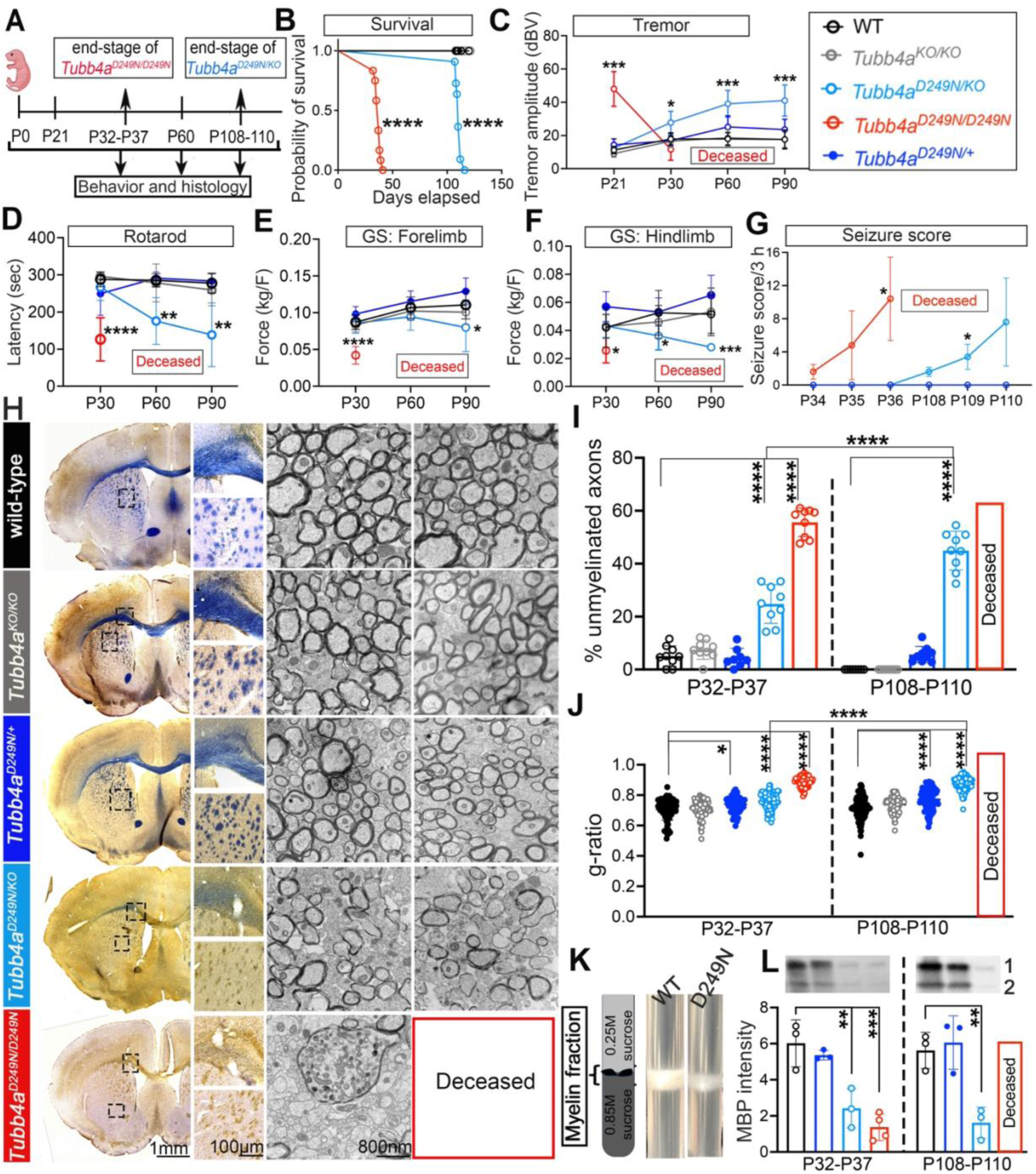
Tubb4a germline deletion is well-tolerated, and *Tubb4a^D249N/KO^* mice show severe H-ABC deficits. **(A)** Timeline schematics depicting the end-stages of H-ABC mouse models. Genotypes include WT, *Tubb4a^KO/KO^*, *Tubb4a^D249N/+^, Tubb4a^D249N/KO^*, and *Tubb4a^D249N/D249N^*. **(B)** Kaplan-Meier survival plot representing lifespans of all genotypes (n=11-15 per genotype). P-values were calculated using the log-rank test. **(C)** Graphical presentation of tremor amplitudes. n=12-15 animals per genotype. Performance of **(D)** Accelerating rotarod, **(E)** Forelimb grip strength (GS), **(F)** Hindlimb GS. n=6-15 mice per genotype and age. **(G)** Seizure counts of 3 hours. n=5 animals per genotype. P-values calculated using repeated measures post hoc two-way ANOVA represent the effect of genotypes compared to WT. See Tables S6 and 7 for detailed p-values of genotype and age interactions. **(H)** Immunohistochemical tiled images of myelin (Eriochrome cyanine [Eri-C]) staining and electron micrograph (EM) images of the corpus callosum at P32-P37 (end-stage of *Tubb4a^D249N/D249N^*) and P108-P110 (end-stage of *Tubb4a^D249N/KO^*). **(I)** Graphical presentation of % unmyelinated axons at P32-P37 and P108-P110. Each point represents the % unmyelinated axon per image (n=2-3 images per mouse). n=3-4 animals per genotype and age. For EM, a two-way ANOVA with Tukey corrections is used to represent the effects of genotypes and ages. See Table S8 for detailed p-values of genotype and age interactions. **(J)** Graphical presentation of g-ratio at P32-P37 in the corpus callosum. n=3-4 animals per genotype and age. For EM, two-way ANOVA with Tukey corrections was used to assess the effects of genotypes and ages. **(K)** Gradient preparation and representative image of the ultracentrifuge tube containing myelin fraction from WT and *Tubb4a^D249N/D249N^*mice. **(L)** Representative MBP western blot (see Figure S4B for the remainder of the cropped blots) and graphical plots of MBP band intensity from purified myelin fraction across WT and *Tubb4a* genotype at P32-P37 and P108-P110. 1=25kDa; 2= 18.5kDa. *p<0.05, **p<0.01 ***p<0.001, ****p<0.0001. For MBP quantification, a two-way ANOVA with Tukey’s correction is used to represent the effects of genotypes. See Table S10 for the detailed p-values. Graphs, which are represented as Mean (SD). See long-term behavior in Figure S1, Figures S2-3 for Eri-C quantification, Figure S4 for Western blots of MBP, PLP, and CNP of isolated myelin fractions, and Figures S5-6 for optic nerve and corpus callosum EM images and quantification.

### Germline suppression of the Tubb4a mutant copy preserves myelin

H-ABC-affected individuals show CNS hypomyelination (1). To examine myelin deficits in mice with varying *Tubb4a^D249N^* dosage, we measured Eriochrome Cyanine (Eri-C) staining and the signal of myelin basic protein (MBP) fluorescent staining, a marker for mature myelin sheaths, at different disease stages. *Tubb4a^KO/KO^*and *Tubb4a^D249N/+^* mice exhibit myelin (Eri-C and MBP signal) similar to WT in the corpus callosum and cerebellum at all ages (**Figure 1H; Figures S2A-E and S3A-C).** Unlike the sparse staining and reduced MBP in *Tubb4a^D249N/D249N^*mice at P21 and P32-P37 (end-stage of *Tubb4a^D249N/D249N^* mice), *Tubb4a^D249N/KO^* mice display moderate myelin (Eri-C) loss with intact MBP sheaths at P21. Nevertheless, these mice experience progressive myelin degeneration (Eri-C and MBP staining) at P32-P37, P60, and at end-stage (P108-P110) in both corpus callosum and cerebellum, resembling end-stage *Tubb4a^D249N/D249N^*mice over time (**Figure 1H; Figure S2-3**). Importantly, *Mbp* mRNA transport is a MT-dependent process (20), and *taeip* rats (p.Ala302Thr Tubb4a homozygous mutation) exhibit altered trafficking of *Mbp* mRNA in OLs (21, 22). Since MBP is essential for myelin compaction (21, 23–25), which is disrupted in our model, we examined whether MBP protein levels are affected in myelin membranes by extracting the whole brain myelin fraction using sucrose density gradient ultracentrifugation. Of note, after myelin purification, we noted no or sparse myelin fraction in *Tubb4a^D249N/KO^* and *Tubb4a^D249N/D249N^*mice (**Figure 1K**). We also assessed the expression of other abundant myelin proteins, proteolipid protein (PLP) and 2’,3’-cyclic nucleotide 3’-phosphodiesterase (CNP). *Tubb4a* mutants (*Tubb4a^D249N/KO^* and *Tubb4a^D249N/D249N^*) show decreased levels of MBP, PLP, and CNP in the myelin fraction compared to WT controls (**Figure 1K-L and Figure S4A-I**). *Tubb4a^D249N/+^* mice have myelin protein levels comparable to WT mice.

To obtain more detailed measures of the ultrastructural complexity of myelin sheaths, we performed electron microscopy (EM) on the optic nerve and corpus callosum at P32-P37 (end-stage of *Tubb4a^D249N/D249N^* mice) and P108-P110 (end-stage of *Tubb4a^D249N/KO^* mice). We calculated the % of unmyelinated axons and the g-ratio, a measure of myelin thickness (26). *Tubb4a^KO/KO^* mice show well-defined and compact myelinated axons in both regions, similar to WT **(Figure 1H-J, Figures S5A-C and S6**). *Tubb4a^D249N/+^* mice display increased g-ratios, indicating thinner myelin in both areas (10) (**Figure 1I-J**, **Figure S5A and S5B [Corpus callosum and Optic nerve]**). At P32-P37, *Tubb4a^D249N/KO^*mice exhibit a higher % of unmyelinated axons, with most axons showing poorly compacted myelin (**Figure 1I and Figure S6**). By end-stage (P108-P110), most myelin has degenerated, as shown by a significant increase in unmyelinated axons and g-ratio (**Figure 1H-J, Figures S5A-C and S6**); however, this myelin degeneration is never as severe as that observed in end-stage *Tubb4a^D249N/D249N^* mice. *Tubb4a^D249N/D249N^*mutants experience early and severe myelin loss compared to all other genotypes, with the sparse myelin sheaths inappropriately thin for their axonal caliber (**Figure 1H-I, Figure S5A-B**). Alongside myelin degeneration, both *Tubb4a^D249N/D249N^* and *Tubb4a^D249N/KO^*mice show increased microglia and macrophages containing myelin debris and vacuoles at the end-stage (**Figure 1H-I, Figures S5A-C and S6**).

Together, these results suggest that lowering the dosage of mutant *Tubb4a* and relatively preserving the WT allele delays and reduces the disease phenotype, supporting a toxic gain-of-function mechanism for the D249N mutation in *TUBB4A*.

### Germline suppression of the Tubb4a mutant copy mitigates cellular pathology

We previously reported that *Tubb4a^D249N/D249N^* mice exhibit a severe reduction in Ols (10). We quantified the total number of double-positive ASPA+ Olig2+ cells (a mature OL marker and an OL lineage marker, respectively) per mm^2^ across all genotypes in the corpus callosum, cerebellar white matter, and granule layer at different ages. OLs and Olig2+ cells remain unchanged in *Tubb4a^KO/KO^* and *Tubb4a^D249N/+^*mice compared to WT mice in these regions (**Figure 2A-D, Figure S7A-E)**. In the corpus callosum, *Tubb4a^D249N/D249N^* and *Tubb4a^D249N/KO^* mice show a significant reduction in ASPA+ Olig2+ cells at P21 and at end-stage (P32-P37 and P108-P110, respectively) relative to WT mice (**Figure 2A-B, Figure S7A**). Similar OL reductions are observed in the cerebellar white matter and granule layers of these mutant models at their respective end stages (**Figure 2A, Figure 2C-D**, and **Figure S7B)**. Olig2+ cell numbers are comparable across all these genotypes (**Figure S7C**), suggesting that overall OL lineage cells are preserved, despite maturation deficits, as reported in our previous results (10).

**Figure 2:**
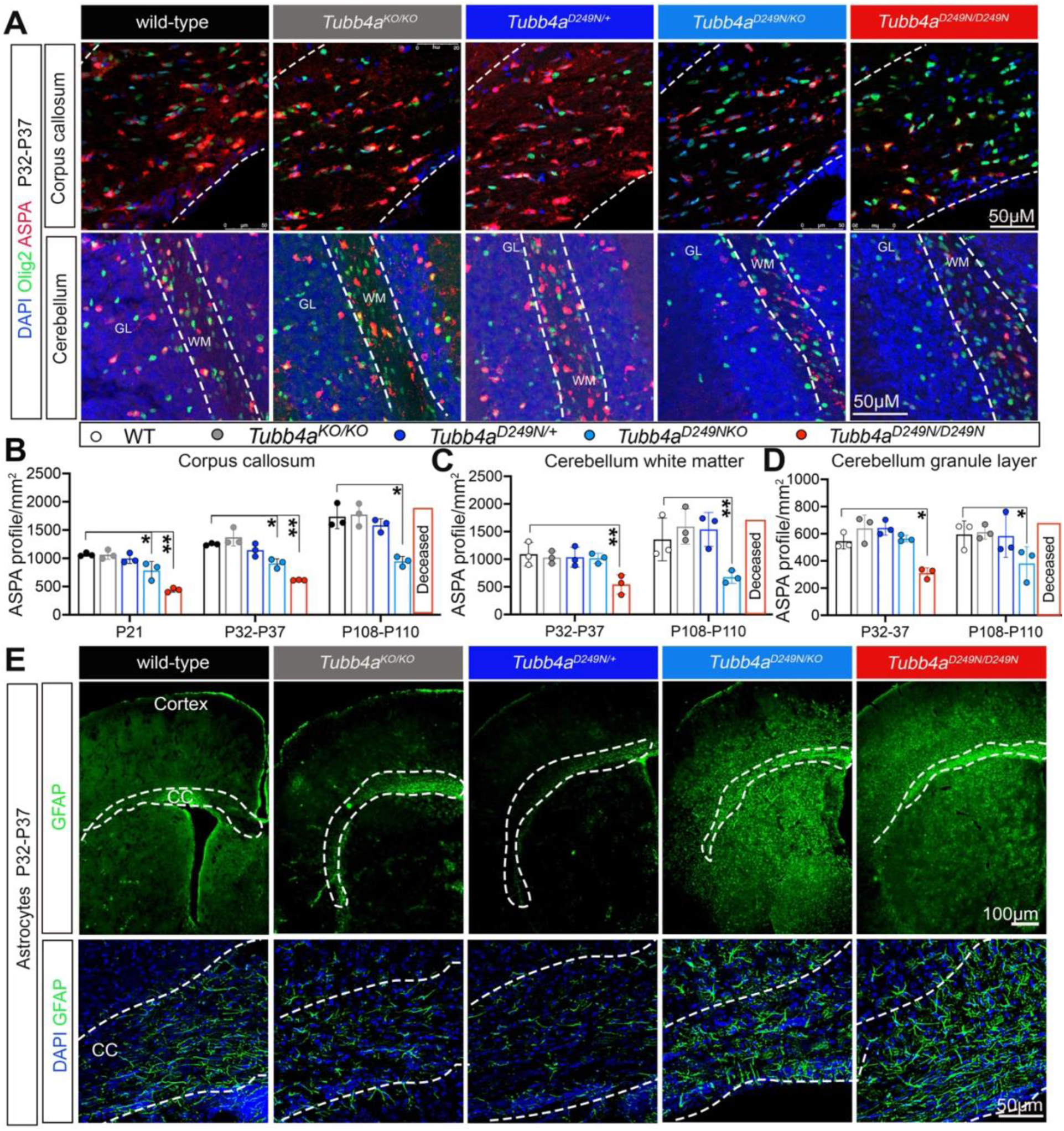
Germline suppression of mutant Tubb4a and relative preservation of WT Tubb4a rescues glial cell pathology. Groups include WT, Tubb4a^KO/KO^, Tubb4a^D249N/+^, Tubb4a^D249N/KO^, and Tubb4a^D249N/D249N^. **(A)** Immunohistochemical images of the corpus callosum and cerebellum at the P32-P37 showing Olig2 (OL lineage marker; green), and ASPA (mature OL marker; red) with DAPI (nuclear marker; blue). GL=granule layer; WM = white matter. **(B-D)** Quantification of ASPA counts per mm^2^ in the corpus callosum (B), cerebellar white matter (C), and cerebellar granule layer (D). n=3-4 mice per genotype and age. **(E)** Immunohistochemical images of the corpus callosum at low (top panel) and high (bottom panel) magnification at the P32-P37 showing GFAP+ cells (green; astrocyte marker) and DAPI+ (blue). For these image panels, all sections are co-stained with Iba1 from Figure S8A-B, and Keyence software was used to overlay the respective channels. P-values for ASPA counts are calculated using two-way ANOVA with Tukey corrections to represent the effects of genotypes compared to WTs. Graphs are represented as Mean (SD). See Table S13 for detailed statistical analysis. *p<0.05, **p<0.01 ***p<0.001, ****p<0.0001. Figure S7 represents immunohistochemical images of ASPA+ Olig2+ cells at P108-P110 and Olig2 quantification at all ages. Other connected Figures include Figure S8-9 (GFAP and Iba1 immunostaining).

In various leukodystrophies, injury promptly activates astrocytes (astrogliosis) and microglia (microgliosis). We immunostained for GFAP (an astrocyte marker) and Iba1 (a microglial marker) at P32-P37 and P108-P110. Astrocytes and microglia are comparable in WT littermates, *Tubb4a^KO/KO^*, and *Tubb4a^D249N/+^* mice, whereas *Tubb4a^D249N/D249N^* and *Tubb4a^D249N/KO^* mice show elevated GFAP and Iba1 density in the corpus callosum and cerebellum (**Figure 2E, Figures S8A-D and S9A-D**).

### Germline suppression of the Tubb4a mutant copy ameliorates neuronal pathology

One of the most notable neurologic phenotypes in H-ABC-affected individuals is cerebellar and basal ganglia atrophy (1). CGNs (cerebellar granule neurons) make up the majority of cerebellar mass (27). As CGNs exhibit apoptotic death in *Tubb4a^D249N/D249N^*mutant mice (10), we quantified CGNs and their cell death by co-immunostaining for NeuN (a neuronal cell marker) and Caspase-3 (an apoptotic marker) in all genotypes across different ages (**Figure 3A**). As expected, *Tubb4a^KO/KO^*and *Tubb4a^D249N/+^* mice do not experience neuronal loss or cell death (**Figure 3B-E, Figures S10B and S11A-B**). In contrast, there is drastic neuronal apoptosis in *Tubb4a^D249N/D249N^*mice at P32-P37 (10). *Tubb4a^D249N/KO^*mice exhibit delayed development, mild neuronal loss, and apoptosis, evident from postnatal day 60 to humane end-stage (P108-P110; **Figure 3B-E, Figures S10B and S11A-B**).

**Figure 3:**
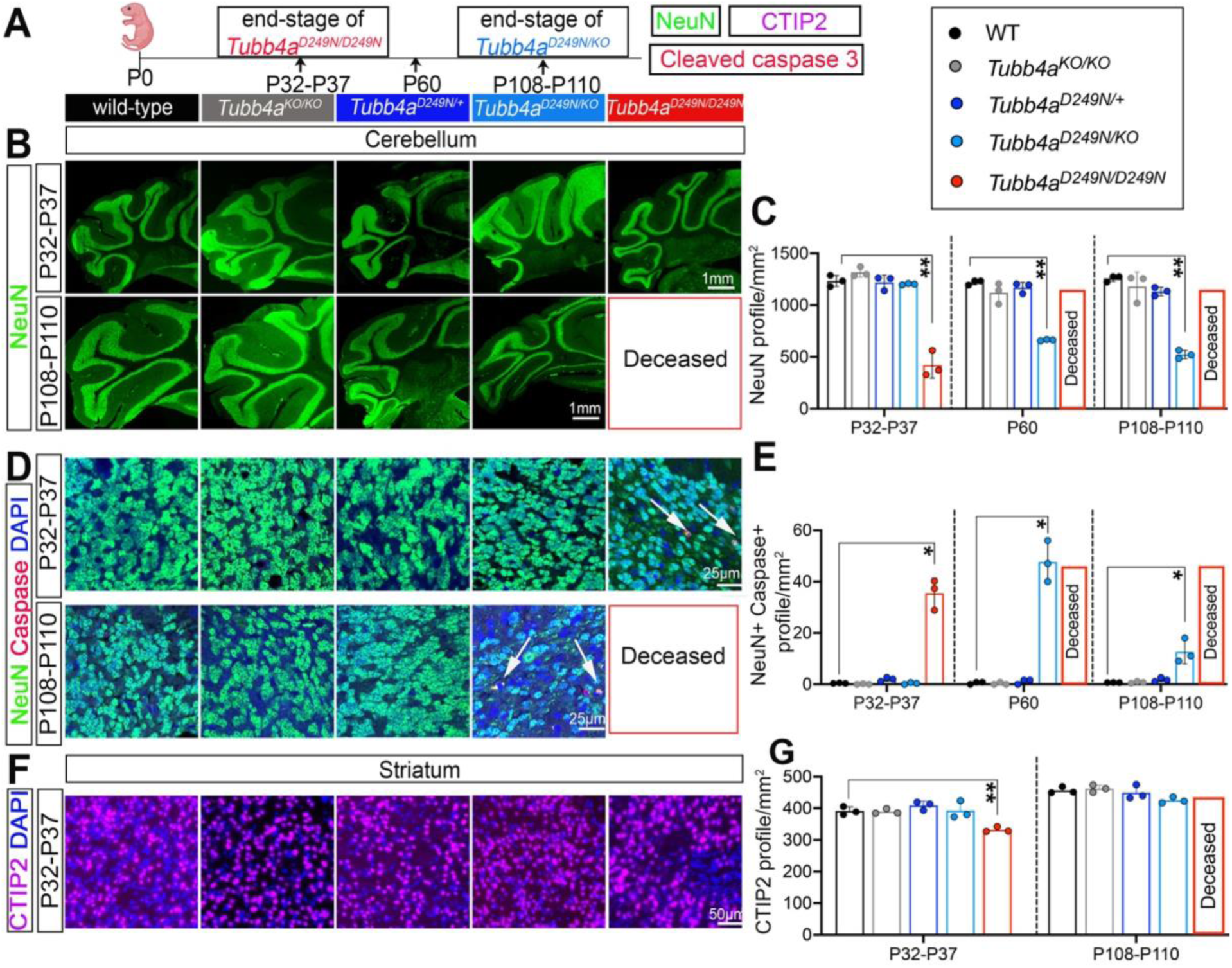
Germline suppression of mutant *Tubb4a* and relative preservation of WT rescues neuronal pathology. **(A)** Timeline schematic of humane endpoints for *Tubb4a* mouse models. Groups include WT, *Tubb4a^KO/KO^*, *Tubb4a^D249N/+^, Tubb4a^D249N/KO^*, and *Tubb4a^D249N/D249N^*. **(B)** Representative tiled images of NeuN immunostaining of cerebellar sections at the P32-P37 and P108-P110. **(C)** Graphical presentation of NeuN profiles per mm^2^ at P32-P37 (end-stage of *Tubb4a^D249N/D249N^*), P60, and P108-P110 (end-stage of *Tubb4a^D249N/KO^*). **(D)** Representative immunofluorescent images of cerebellum sections at high magnification at P32-P37 and P108-P110 that are stained with NeuN (green; neuronal marker), cleaved Caspase 3 (red; apoptotic marker), and DAPI (blue). Arrows represent co-localized NeuN+ cleaved Caspase 3+ cells. Unmerged individual channel images are provided in Figure S11. **(E)** Graphical presentation of double-positive NeuN+ Caspase+ profiles per mm^2^. **(F)** Representative images of striatum sections at the P32-P37 stained with CTIP2 (magenta; MSN marker) and DAPI (blue). **(G)** Graphical presentation of CTIP2 counts per field. n=3-4 animals are used per genotype and age. P-values are calculated using two-way ANOVA with Tukey corrections to represent the effects of genotypes versus WTs. Graphs are represented as Mean (SD). See Table S15 for detailed p-values. *p<0.05, **p<0.01 ***p<0.001, ****p<0.0001. Associated Figure S10 includes representative images of the P60 cerebellum and NeuN+ and CTIP2 profiles in the striatum for other ages.

To assess the basal ganglia phenotype, we examined the number of striatal neurons at P32-P37 and P108-P110 across all genotypes by immunofluorescent staining of NeuN and CTIP2 antibodies. CTIP2 (a MSN marker) denotes striatal neurons, the neuronal population with the highest deficit in H-ABC-affected individuals (28) (5). Notably, no striatal neuron (CTIP2+) loss is evident in *Tubb4a^KO/KO^* and *Tubb4a^D249N/+^* mice (**Figure 3F-G, Figure S10C-E**). *Tubb4a^D249N/D249N^* mice at end-stage (P32-P37) show a significant reduction in CTIP2+ and all striatal neurons (NeuN; **Figure 3F-G**, and **Figure S10C-E**). *Tubb4a^D249N/KO^* mice exhibit a trend of decreased striatal neurons at the end-stage (P108-P110), though the reduction is not statistically significant (**Figure 3F-G, Figure S10D-E**). We also measured striatal volume across all genotypes, which were comparable to WT (**Figure S10F**).

### Spatial *Tubb4a* transcripts demonstrate robust expression in H-ABC-relevant brain areas

To understand the spatial-temporal expression of *Tubb4a* mRNA at different ages, we conducted RNAscope *in situ* hybridization at P0, P7, P14, P21, P32-P37 and P108-P110 in WT and *Tubb4a^KO/KO^* sagittal brain sections. The mouse *Tubb4a* probe was specific, as no signal was detected in the *Tubb4a^KO/KO^* brain sections (**Figure S12A-B**).

Within the forebrain at P0, a few weakly stained *Tubb4a* cytoplasmic puncta are present in cells within the cortical layers, striatum, and corpus callosum (**Figures 4A-E**). At P7 in the cortical layers, we observed *Tubb4a* transcripts localized to layer I; however, by P14, most cortical layers show an even distribution of transcripts (**Figure 4I, 4K, and Figure S13B**). At all ages after P7, cells containing saturated *Tubb4a* transcripts resemble OL lineage cells (shown by arrow in **Figure 4, See Corpus callosum [CC] and Figure S13**). *Tubb4a* transcripts increase from P7 to P21 in different brain areas with stable *Tubb4a* expression patterns at P35 and P110 (**Figure 4, Figures S12 and S13**), suggesting a crucial role of Tubb4a in the early neurodevelopmental phase.

**Figure 4:**
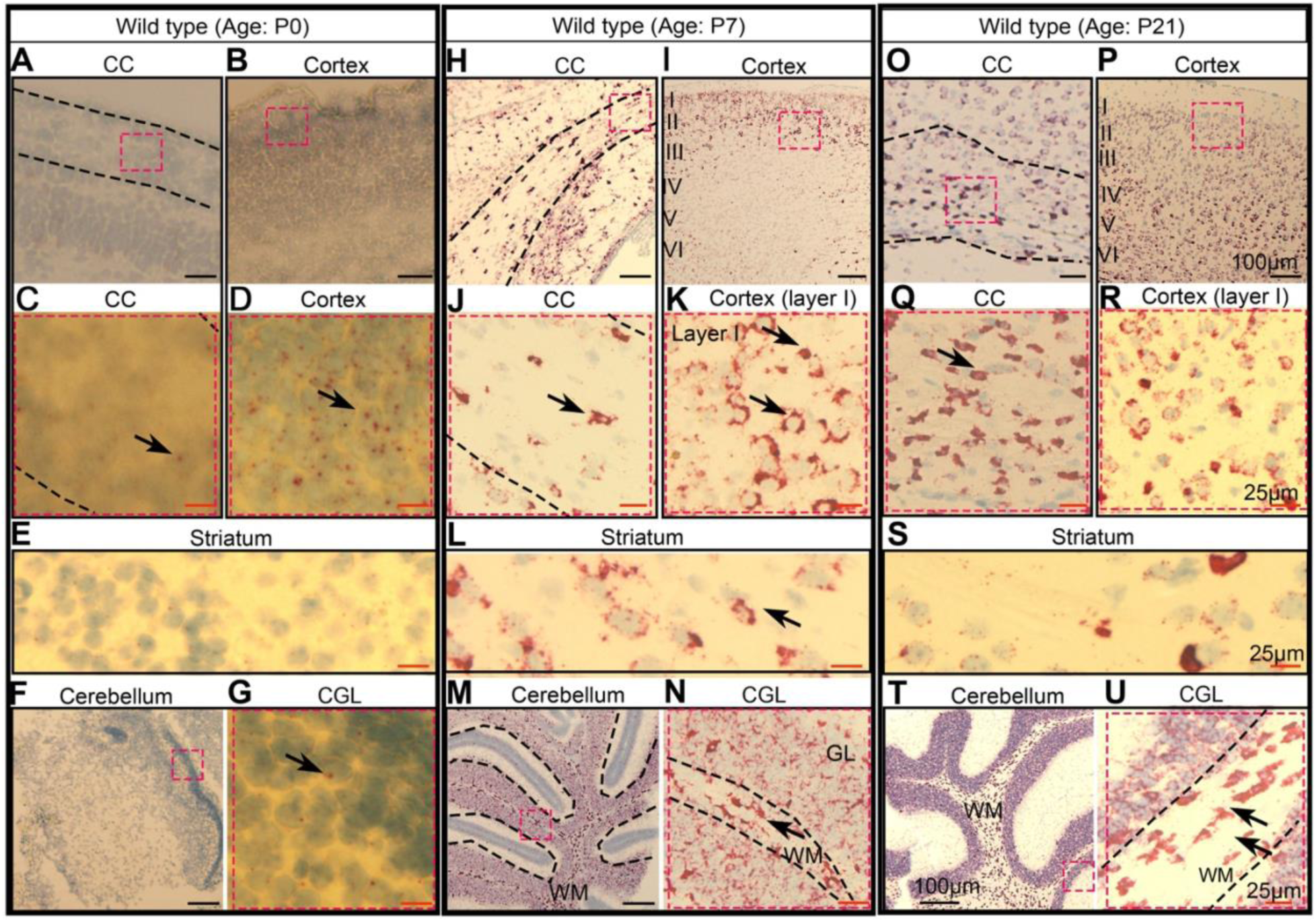
RNAscope *in situ* hybridization detecting Tubb4a transcripts at P0, P7, and P21 in WT sagittal brain sections. (**A-G**) Representative images of low [A-B, and F] and high [C-D, E, and G] magnification Tubb4a RNAscope at P0 of the cortex, corpus callosum (CC), striatum, and cerebellum. (**H-N**) Representative images of low [H-I, M] and high [J-K, L, and N] magnification of RNAscope detecting *Tubb4a* transcripts at P7 of the cortex, CC, cerebellum, and striatum. (**O-U**) Representative images of low [O-P, T] and high [Q-R, S, and U] magnification of RNAscope detecting *Tubb4a* transcripts at P21 of the cortex, CC, striatum, and cerebellum. Arrows are used to denote a few *Tubb4a* transcripts. Tiled images of all corresponding ages are shown in Figure S12. Figure S13 provides Tubb4a RNAscope images at P14, P32-P35, and P108-P110.

Similarly, we examined cerebellar white matter and neuronal layers for *Tubb4a* transcripts across all ages. In the cerebellar white matter, from age P7, the pattern of *Tubb4a* expression follows a similar time course as that observed in the corpus callosum (**Figure 4F-G, 4M-N, and 4T-U, Figure S13F-G, S13M-N, and S13T-U**). The cerebellar granule layer (CGL) includes multiple cell types, such as a large volume of CGNs, Golgi cells, Bergmann glia, and OL lineage cells. At P0, the cerebellar granule progenitors are located in the external granule layer and have not migrated yet (29). In this non-migrated external granule layer, we noted a few early cytoplasmic *Tubb4a* puncta at P0 (**Figure 4F-G**). Over time in the internal granule layer, these CGNs show increased Tubb4a transcripts from P7 to P21 and reach stable expression at later time points (**Figure 4M-N and 4T-U, Figure S13F-G, S13M-N, and S13T-U**).

### Anti-sense administration reduces Tubb4a *in vitro* with minimal toxicity

Given that germline *Tubb4a* reduction is well-tolerated, we explored the translational potential of lowering Tubb4a levels using ASOs. Considering Tubb4a’s crucial role in early neurodevelopment, we proposed early intervention. We designed a series of ASO candidates targeting mouse *Tubb4a* (coding regions). All ASOs included in the initial screen contained a 2’MOE modification with a phosphorothioate (PS) backbone, characteristics that are known for stability and low cytotoxicity (30, 31). ASO design included gapmers with complete PS bonds or reduced PS bonds to reduce ASO-associated toxicity (30) (ASO designs are provided in **Table S1**).

ASOs were screened in WT mouse cortical neurons *in vitro* by free uptake (gymnosis), followed by analysis of *Tubb4a* transcripts using qRT-PCR. To assess *in vitro* ASO toxicity, we performed the ApoTox-Glo Triplex assay, which measures viability, cytotoxicity, and caspase activation within a single well. We identified and selected ASOs (8 ASO candidates) that effectively reduce Tubb4a (>80%) at 5µM. *Tubb4a* downregulation of selected potential ASOs is listed in **Table S2.** The ASOs in bold in **Table S2** are the 8 chosen candidates.

### Tubb4a ASO 18-3 administration *in vivo* via i.c.v route exhibits maximum potency with minimal toxicity

To investigate ASO toxicity and efficacy *in vivo*, we injected selected ASOs (8 ASO candidates) at a dose of 15µg/g via an i.c.v. route in WT mice at P1 alongside vehicle-treated animals that received 2 µL sterile PBS. Pups injected with ASO candidates 4, 6, and 7 died within 3-4 hours after injection, which was attributed to acute neurotoxicity (**Table S3**) (32). To examine the chronic neurotoxicity and off-target effects of the remaining ASO candidates (ASO 7-2, ASO 7-3, ASO 8-3, ASO 18-1, and ASO 18-3), we monitored the mice weekly using a functional observational battery (FOB) at P30 (30 days post-injection). FOB is a well-established method for quantifying behavioral neurotoxicity (33). WT mice injected with ASO 7-2 and ASO 18-1 displayed mild to severe neurologic deficits, including poor posture, ataxia, weak hindlimb grasp, and hyperactivity (**Table S4**). WT mice injected with ASO 7-3 and ASO 8-3 exhibit mild gait deficits, weight loss, and impaired motor coordination as assessed by rotarod (**Table S4, Figure S14A**). Out of our 8 ASO candidates, we observed that WT mice injected with ASO 18-3 performed similarly to PBS-injected controls on the neurobehavioral battery and rotarod tests (**Table S4, Figure S14A and C**). Despite mild motor deficits observed with ASO 7-3 and ASO 8-3, we also measured their ability to downregulate *Tubb4a* to compare efficacy with ASO 18-3. After completing the neurobehavioral assays, we dissected and isolated different brain regions (cortex, cerebellum, and striatum) 30 days post-injection and used qRT-PCR to measure *Tubb4a* expression levels. All three ASO candidates showed similar *Tubb4a* suppression potential, but because ASO 18-3 demonstrated a more consistent safety profile at 15µg/g, it was selected for future experiments **(Figure S14B**).

### Administration of Tubb4a ASO 18-3 results in dose-dependent, long-lasting, and specific decreases of Tubb4a in WT and *Tubb4a^D249N/KO^* mice

To determine the dose-dependent *in vivo* safety profile and the potential for *Tubb4a* downregulation of ASO 18-3, we performed i.c.v. injections of ASO 18-3 at P1 in WT mice with different doses (5,15, 20, and 30 µg/g, n=3-4 mice per dose) along with vehicle-treated (PBS) controls. We monitored these mice weekly and observed dose-related lethality at a 30µg/g dose (75% of mice died within a few days post-injection). The surviving 25% of mice injected with 30 µg/g exhibited atypical FOB scores (**Figure S14C**). Analysis of FOB and motor behavioral assays at 15 and 20 µg/g doses at P30 showed typical neurological scores compared to PBS-injected controls (**Figure S14C**).

To assess dose-dependent *Tubb4a* downregulation, we dissected and isolated different brain regions at P30 and measured *Tubb4a* mRNA levels relative to PBS-injected controls using qRT-PCR. The administration of *Tubb4a* ASO 18-3 resulted in a dose-dependent knockdown of *Tubb4a* mRNA in the cortex and striatum with no significant decrease in the cerebellum (**Figure 5B and Figure S15A**). Tubb4a protein levels analyzed by immunoblotting at the 20 µg/g dose matched with mRNA expression levels at P30 (**Figure 5B-C**; Tubb4a antibody specificity confirmed with *Tubb4a^KO/KO^* brain tissue; **Figure S15C**).

**Figure 5:**
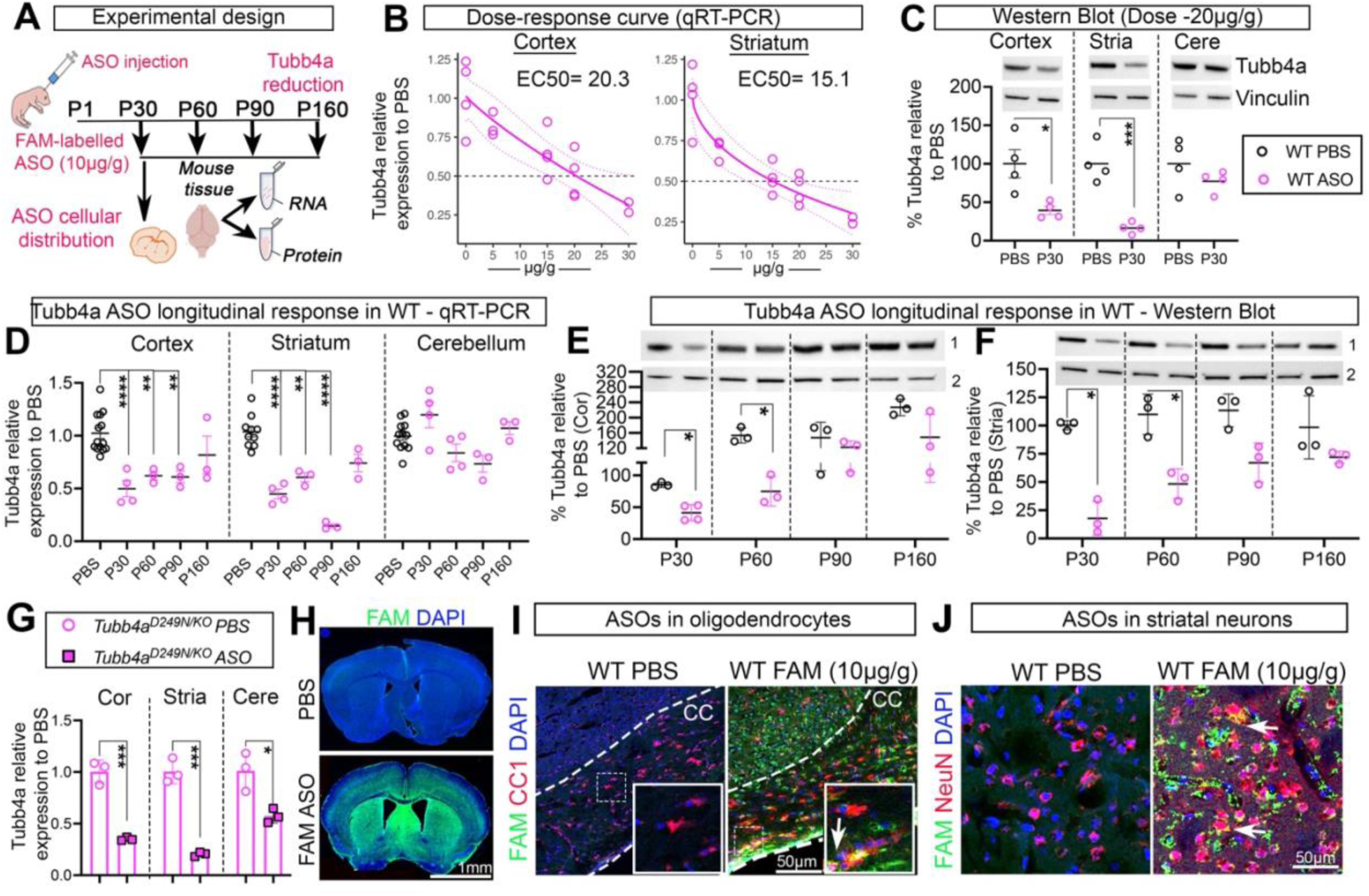
ASO 18-3 *in vivo* distribution in WT mice. **(A)** Experimental design. **(B)** Dose-dependent *Tubb4a* expression relative to PBS-injected controls at P30 in the cortex and striatum via qRT-PCR. **(C)** Western blot of Tubb4a protein at 20µg/g of ASO at P30 in the cortex, striatum, and cerebellum. n=3-4 WT mice are used for each dosage. **(D)** Longitudinal *Tubb4a* expression in WT mice was analyzed using qRT-PCR at 20µg/g of ASO 18-3 in different brain areas at P30, P60, P90, and P160. **(E-F)** Longitudinal Tubb4a protein levels in WT mice at P30, P60, P90, and P160 in the cortex (E) and striatum (F), 1=Tubb4a; 2=vinculin (housekeeping protein). n=3-4 WT mice are used for corresponding ages for Tubb4a expression. **(G)** Tubb4a expression via qRT-PCR in ASO-treated *Tubb4a^D249N/KO^* mice relative to PBS-treated *Tubb4a^D249N/KO^* mice at 20µg/g. For qRT-PCR, the *sfrs9* was used as a housekeeping gene to normalize the *Tubb4a* expression. For the Western blot, Tubb4a protein levels are normalized to vinculin. P-values are calculated using one-way or two-way ANOVA with Tukey corrections, and detailed p-values are provided in Table S17. Graphs are represented as Mean (SD). *p<0.05, **p<0.01 ***p<0.001, ****p<0.0001. **(H)** Tiled images of coronal forebrain sections of DAPI (blue) and FAM (green). **(I)** Representative immunofluorescent images of the corpus callosum showing co-staining of CC1 (red; mature OL marker) with FAM (green) and DAPI (blue). Unmerged and merged channel images are provided in Figure S16A. **(J)** Representative immunofluorescent images of the striatum region showing NeuN (red; neuronal marker) co-staining with FAM (green). Unmerged and merged channel images are provided in Figure S19B. Arrows represent co-localized signals. Associated Figures include S14-19.

To assess the long-term persistence of *Tubb4a* downregulation, we performed i.c.v. ASO injections at P1 at a dose of 20 µg/g in WT mice and collected tissues at P30, P60, P90, and P160. We measured both *Tubb4a* mRNA and protein levels in the cortex, striatum, and cerebellum at these ages using qRT-PCR and Western blot. We observed significant Tubb4a downregulation until P60; however, Tubb4a levels returned to normal at P90 and P160, indicating that ASO 18-3 is effective for approximately 8-9 weeks (**Figure 5D-F**). Similarly, qRT-PCR showed significant *Tubb4a* downregulation in the cortex, striatum, and cerebellum in *Tubb4a^D249N/KO^* mice (**Figure 5G**). Of note, we did not analyze the Tubb4a protein downregulation in *Tubb4a^D249N/KO^* mice as the Tubb4a protein is downregulated in mutant mice relative to WT (**Figure S15C**).

### OLs, cortical, and striatal neurons show robust Tubb4a 18-3 ASO uptake

To investigate the spatial uptake of ASO in different brain cells and regions, we generated fluorescently labeled Tubb4a ASO 18-3 using fluorescein amidites (FAM). Administering FAM-tagged ASO 18-3 at 20µg/g via i.c.v. injection in postnatal WT mice resulted in 100% mortality, likely due to a synergetic toxicity effect between ASO and FAM. Therefore, we reduced the FAM-ASO dose to 10 µg/g and administered this ASO via i.c.v. injection in pups. This lower dose was well-tolerated in all pups and was used to assess ASO distribution in various brain regions, including corpus callosum, cortex, striatum, and cerebellum.

We observed a strong ASO distribution in the forebrain (**Figure 5H**), particularly near the lateral ventricles, cortex, corpus callosum, and striatum. The cerebellum shows minimal to moderate ASO uptake with no fluorescent signal in the granule layers (**Figure S15D**). To evaluate the ASO distribution in OL lineage cells, we performed co-staining of CC1 (a mature OL marker) with FAM and NG2 (an OL precursor cell (OPC) marker) with FAM. We found robust ASO uptake in OPCs and OLs in the corpus callosum, cortex, and striatum (**Figure 5I**, **Figures S16-18**). Cerebellar white matter exhibits ASO uptake in OLs, though it is less extensive than corpus callosum (**Figure S17B**). Additionally, we co-stained NeuN with FAM to assess ASO uptake in neurons. We observed ASO distribution in cortical and striatal neurons with no significant ASO uptake in CGNs (**Figure 5J**, **Figure S19**).

### ASO-improved motor function and survival in *Tubb4a^D249N/KO^* mice

We next evaluated whether a single i.c.v. injection of the ASO candidate 18-3 effectively rescues motor phenotypes and improves survival in postnatal (P1-2) *Tubb4a^D249N/KO^* mice (Study design, **Figure 6A**). Remarkably, *Tubb4a^D249N/KO^* mice injected with Tubb4a ASO 18-3 at a dose of 20 µg/g show greatly extended survival (**Figure 6B**, Mean survival of P365-P370) compared to PBS-injected *Tubb4a^D249N/KO^* mice (Mean survival of P108-P110). We conducted accelerating rotarod, grip strength, and tremor assays to determine whether ASO 18-3 could improve motor function in *Tubb4a^D249N/KO^* mice. Tubb4a ASO-treated *Tubb4a^D249N/KO^* mice display significantly improved rotarod performance relative to untreated mice until P120, suggesting improved balance and coordination (**Figure 6C, Videos 1 and 2** show improved phenotype in treated *Tubb4a^D249N/KO^* mice versus untreated *Tubb4a^D249N/KO^* mice). Their grip strength and tremor assay performance were also notably enhanced (**Figure 6D-F)**. However, motor function in ASO-treated *Tubb4a* mutants starts to decline after P90, corresponding to a decrease in ASO efficacy and a return to baseline Tubb4a expression between 60-90 days following a single i.c.v. injection of ASO (**Figure 5D-F**). We also tested if post-natal ASO injection shows an improvement of H-ABC phenotypes in severe homozygous *Tubb4a^D249N/D249N^* mice. We observed significantly improved motor phenotypes and survival from ∼P32-P37 to ∼P50-P60 (**Figure S20A-C**). As *Tubb4a^D249N/KO^* mice are physiologically closer to the human disorder with the presence of one mutation copy, we used *Tubb4a^D249N/KO^* mice as a murine model for further preclinical studies.

**Figure 6:**
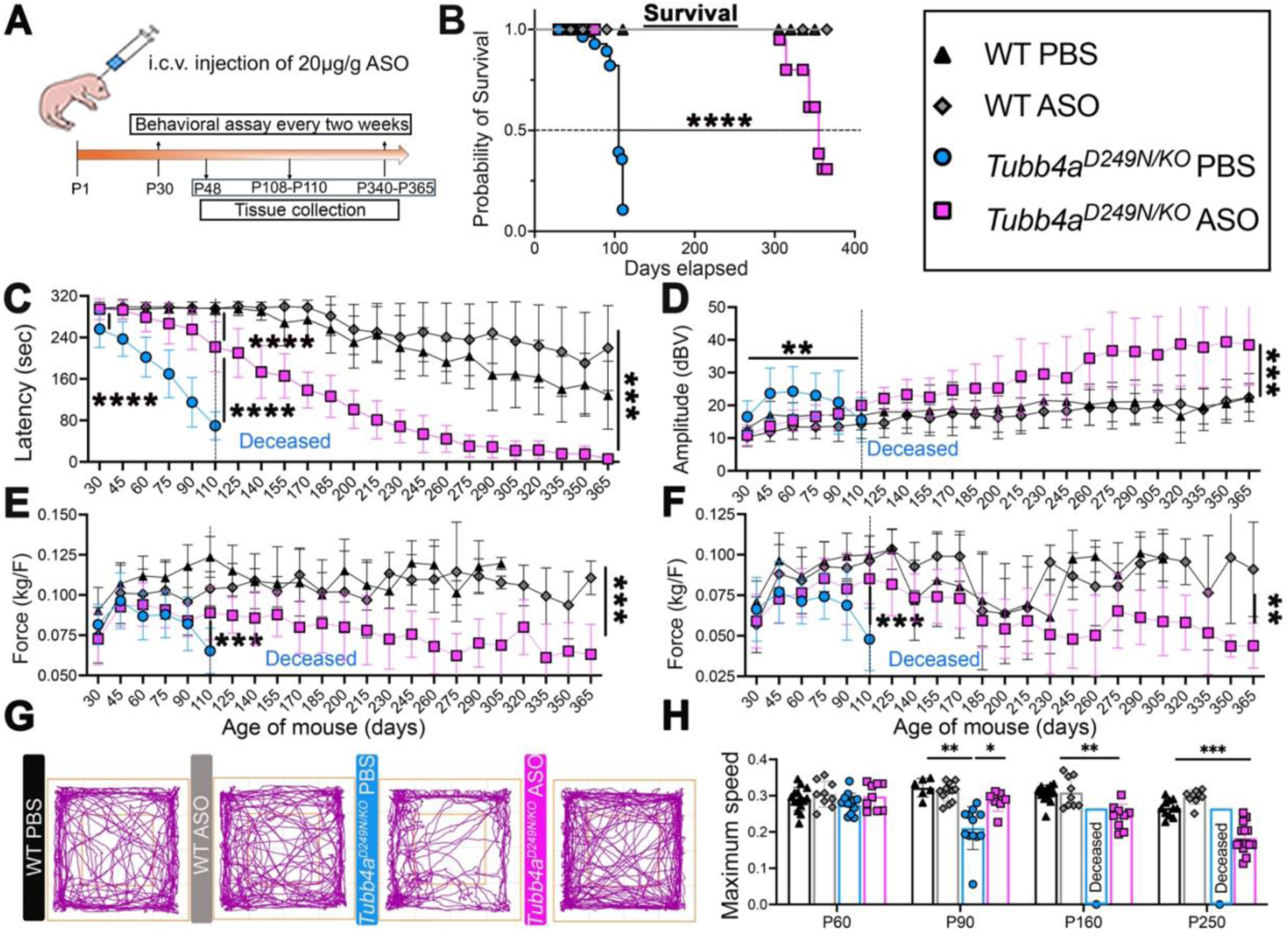
Survival and behavior in ASO-treated groups. Treatment groups include WT PBS, WT ASO, *Tubb4a^D249N/KO^* PBS-treated, and *Tubb4a^D249N/KO^* ASO-treated mice. **(A)** Experimental design. **(B)** Kaplan-Meier survival analysis of different treatment groups with n=25-40 mice per group. P-values were calculated using the log-rank test. **(C)** Graphical presentation of the accelerating rotarod test of treatment groups. **(D)** Graphical presentation of tremor amplitudes of treatment groups. (**E-F**) Graphical presentation of grip strength: Forelimb (E) and Hindlimb grip strength (F). n=20-25 mice were tested per group for rotarod, tremor assay, and grip strength. **(G)** Representative images of exploratory activity of the open field at P90. **(H)** Graphical presentation of the maximum speed of treatment groups. n=12-13 mice were tested per group. For behavioral tests, p-values are calculated using repeated measures ANOVA with mixed-effects analysis, representing the effects of the treatment groups. Graphs are represented as Mean (SD). Detailed p-values are provided in Table S19.

Of note, because untreated *Tubb4a^D249N/KO^* mice die at P108-P110, the surviving ASO-treated *Tubb4a^D249N/KO^* mice were compared with WT untreated and WT ASO-treated mice. Importantly, even after rotarod performance declines, Tubb4a ASO-treated *Tubb4a^D249N/KO^* mice remained mobile until P250 in open-field testing and performed similarly to WT PBS-treated mice (**Figure 6G-H)**. However, activity decreased at P300 (close to their end-stage). This data suggests that ASO 18-3 treatment delays the onset of motor deficits in H-ABC mice long after the ASO effect ceases to decrease Tubb4a expression levels. Of note, both Tubb4a ASO-treated *Tubb4a^D249N/KO^* and WT ASO-treated mice show increased exploratory activity relative to PBS-treated WT mice without changes in % freezing (data not shown) and average speed (**Figure 6H** and **Figure S21A).** Increased exploratory activity is often linked to anxiety, but the cause of this behavior in relation to ASO treatment remains unknown. Notably, WT ASO-treated mice also exhibit generalized astrogliosis, as indicated by an increase in *Gfap* transcripts via Nanostring **(Figure S22A-B**), which is also associated with heightened exploratory activity (34).

As we observed astrogliosis and microgliosis in the 18-3 injected mice, we investigated whether these phenomena are linked to the ASO sequence or ASO modifications. To assess this, we designed a scrambled ASO (non-targeting control (NTC)) that matches the ASO 18-3 modifications. We synthesized this NTC ASO and tested it *in vitro* in cortical neurons alongside three other ASO candidates (ASO 7-3, ASO 8-3, and ASO 18-3) for the downregulation of *Tubb4a* (**Figure S23A**). We injected NTC ASO postnatally at P1-2 at a dose of 20 µg/g in WT mice and assessed motor function at P30 compared to WT-PBS injected mice. We noted no abnormalities in behavior (**Figure S23B**). Next, we stained for GFAP and Iba1 and observed no astrogliosis or microgliosis in WT NTC ASO-injected mice relative to WT PBS-injected mice at P30 (**Figure S23C**). This indicates that the astrogliosis and microgliosis may be specific to the ASO 18-3 sequence. Therefore, we included WT-PBS, WT ASO-treated, and *Tubb4a^D249N/KO^*PBS mice as controls for subsequent studies.

### ASO-mediated suppression of Tubb4a effectively retains functional myelin and OLs

To assess myelination and related OL pathology after ASO 18-3 treatment, we measured myelin using *Mbp* transcripts and MBP protein, Eri-C staining, and EM in different brain regions. *Mbp* transcripts (detected by Nanostring panel) and MBP protein (by Western blot) are retained in ASO-treated *Tubb4a^D249N/KO^* mice in the striatum and cortex (**Figure S21B-C and Figure S22 for *Mbp* Nanostring**). *Mbp* mRNA is transported via MTs and then locally synthesized in OL processes (20–22). Since we observed increased *Mbp* mRNA expression in the ASO-treated *Tubb4a^D249N/KO^* mice using Nanostring (**Figures S21B-C and S22 for *Mbp* Nanostring)**, we then examined whether ASO treatment enhances *Mbp* mRNA transport to the myelin sheath. To investigate this, we used an RNAscope approach to assess *Mbp* mRNA distribution in OLs (labeled with ***Myrf* mRNA probe**) in the corpus callosum across the treatment groups. At P48, untreated *Tubb4a^D249N/KO^* mutants showed saturated *Mbp* mRNA in cell bodies with no or sparse *Mbp* mRNA (red) distribution on the myelin sheath (**Figure 7A; Figures S24 and S25**) as compared to WT controls, suggesting that the *Tubb4a* mutation has disrupted the transport of *Mbp* mRNA into the myelin compartment. Notably, ASO treatment in *Tubb4a^D249N/KO^*mice significantly enhanced the transport of *Mbp* mRNA on myelin sheaths compared to PBS-treated *Tubb4a^D249N/KO^* mice (**Figure 7A and 7D; Figures S24 and S25**). Our IHC data support increased MBP protein expression in the corpus callosum with ASO treatment (**Figure 7B and 7E**).

**Figure 7:**
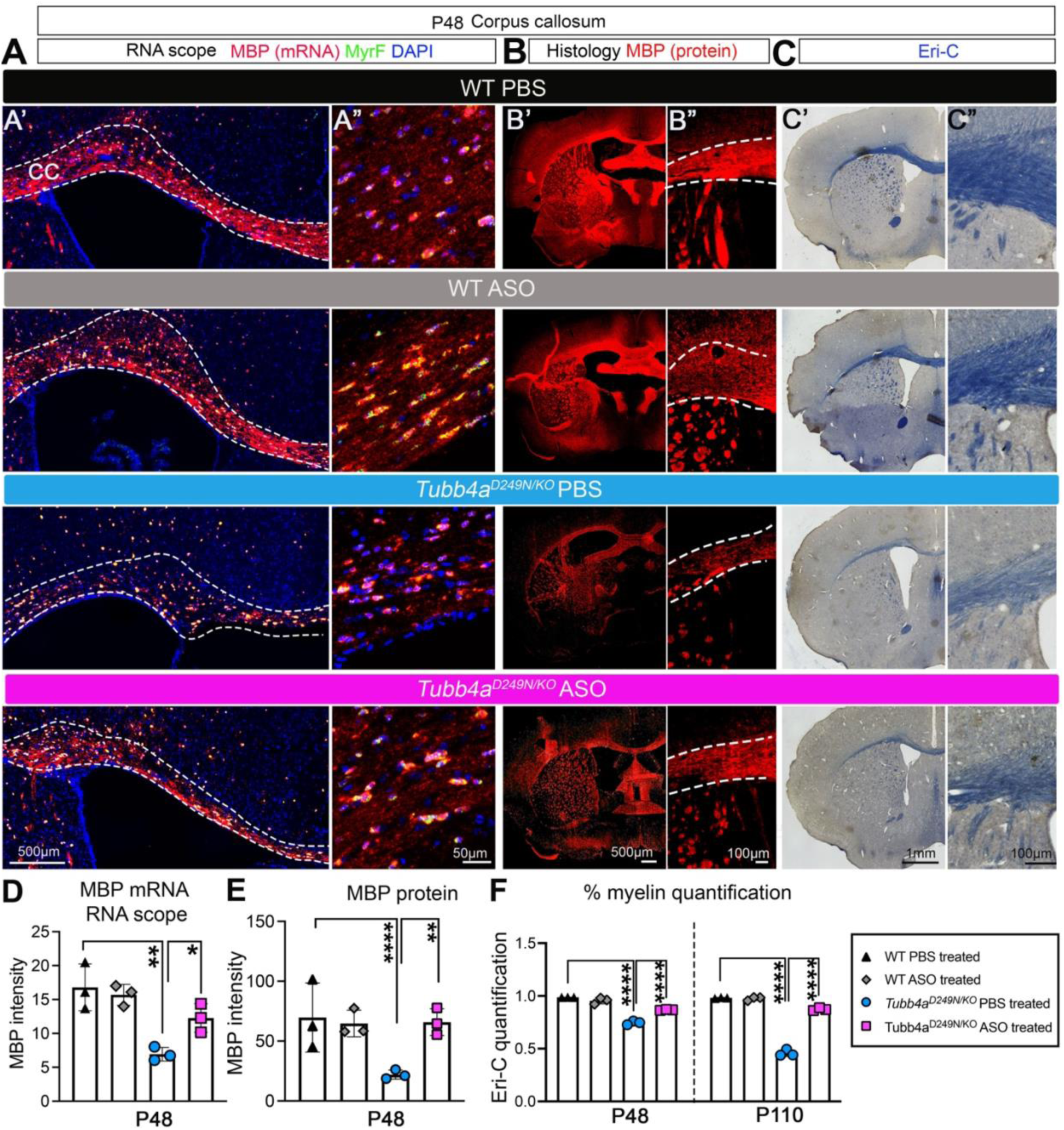
Myelin quantification in ASO-treated groups. Treatment groups include WT PBS, WT ASO, *Tubb4a^D249N/KO^* PBS-treated, and *Tubb4a^D249N/KO^*ASO-treated mice. **(A)** A’ and A’’ columns represent tiled and high magnification images of RNAscope of corpus callosum sections at age P48 representing *Mbp* mRNA (red), *Myrf* mRNA (green; representing OL cell bodies) and DAPI (blue) across treatment groups. Co-localized yellow puncta represent OLs. Unmerged individual channel images are provided in Figures S24 and S25. (**B**) B’ and B” columns represent tiled and high magnification images of MBP immunostaining in the forebrain at P48 across treatment groups. **(C)** C’ and C” columns represent tiled and high magnification images of Eri-C (myelin) staining in the forebrain at P48. **(D)** Graphical presentation of *Mbp* mRNA levels from RNAscope images across treatment groups. **(E)** Graphical presentation of *Mbp* intensity of corpus callosum (histology) across treatment groups. **(F)** Graphical presentation of myelin quantification (Eri-C) analysis across treatment groups. n=3 mice per treatment group. P-values are calculated using one-way or two-way ANOVA with Tukey corrections. *p<0.05, **p<0.01 ***p<0.001, ****p<0.0001. Graphs are represented as Mean (SD). Associated Figures include S20-25.

Next, we used EM to measure the g-ratio and % unmyelinated axons in the corpus callosum and optic nerve across all treatment groups to assess the myelin levels at high resolution. ASO-treated *Tubb4a^D249N/KO^* mice show increased myelination in both areas relative to untreated *Tubb4a^D249N/KO^* mice as measured by g-ratio and % unmyelinated axons (**Figure 8B-D; Figure S26A-E)**. Of note, myelin in treated *Tubb4a^D249N/KO^* mice remains less compact than untreated WT mice. Additionally, at higher resolution at ∼P110, myelin vacuolization is decreased in treated mice relative to untreated *Tubb4a^D249N/KO^* mice (**Figure 8E; Figures S26E and S27A-D**). We also observed decreased microglia and/or macrophages containing myelin debris in treated *Tubb4a^D249N/KO^* mice relative to untreated mutant mice **(Figure 8B; Figure S27A-D**). Similarly, myelin levels (Eri-C analysis) are enhanced in ASO-treated *Tubb4a^D249N/KO^*mice relative to PBS-treated *Tubb4a^D249N/KO^* mice in the corpus callosum, striatum, and cerebellar white matter (**Figures 7C and 7F, 8F and 8I; Figure S28A-D**); this effect persisted in forebrain but not in the cerebellum through one year without additional ASO dosing (**Figure S28A-D**). Similarly, ASO treatment restored mature OLs (ASPA+ Olig2+) in treated H-ABC mice (corpus callosum at P48 and P108-P110), unlike their significant depletion in untreated *Tubb4a^D249N/KO^*mice (**Figure 8H and 8J; Figure S29A**). Next, we quantified the proportion of pre-myelinating OLs (BCAS1+ Olig2+) across all treatment groups. We observed a significant reduction in BCAS1+ Olig2+ OLs in untreated *Tubb4a^D249N/KO^* mice compared to untreated WT mice (**Figure S29B-C**). With ASO treatment, we noted a restoration of BCAS1+ OLs (**Figure S29B-C**). Lastly, we quantified the number of OPCs across treatment groups. We previously showed that OPC numbers remained constant, but OPC proliferation increased in *Tubb4a^D249N/D249N^* mice throughout the disease (10). Consistent with these results, we noted no change in the number of NG2+ OPCs across all treatment groups (**Figure S29D-E**). To assess whether OPC proliferation is mitigated by ASO treatment, we conducted co-immunostaining of NG2+ and Ki67+ (a proliferation marker) cells. Qualitatively, we identified proliferating OPCs (NG2+ Ki67+ cells) in untreated *Tubb4a^D249N/KO^* mice but none in WT controls (**Figure S29F**). Similarly, we noted no NG2+ Ki67+ OPCs in treated *Tubb4a^D249N/KO^* mice relative to untreated *Tubb4a^D249N/KO^*mice (**Figure S29F**). Olig2+ cells remain comparable in all treatment groups (**Figure S29G**).

**Figure 8:**
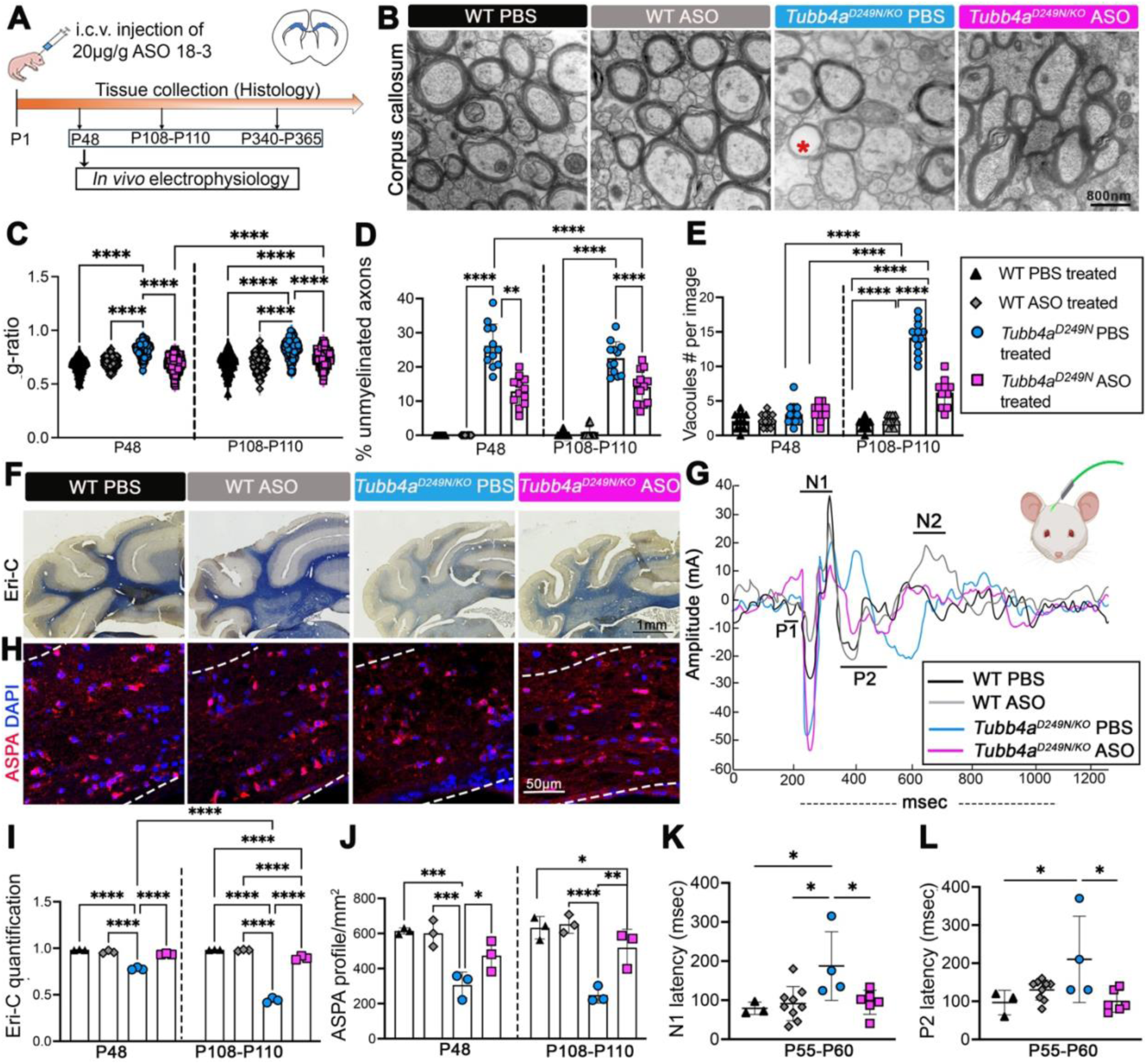
Neuropathological analysis in ASO-treated groups. Treatment groups include WT PBS, WT ASO, *Tubb4a^D249N/KO^*PBS-treated, and *Tubb4a^D249N/KO^* ASO-treated mice. **(A)** Experimental design. **(B)** Representative EM images of ASO-treated mice of the corpus callosum at P48. **(C)** Graphical presentation of g-ratio across different treatment groups. **(D)** Graphical presentation of % unmyelinated axons across different treatment groups. **(E)** Graphical presentation of the number of vacuoles (Vacuoles #). For EM data, n=3-4 animals are used, and at least 4-5 images per animal are used. At least 30-40 axons per image were used to measure the g-ratio. **(F)** Representative images of Eri-C (myelin) staining in the cerebellum at P48. **(G)** The mean of **the** visual evoked potential peaks of all groups was generated using MATLAB. **(H)** Representative immunofluorescent images of ASPA+ (red) DAPI+ (blue) cells in the corpus callosum at P48. **(I)** Graphical presentation of myelin quantification (Eri-C) analysis across treatment groups. **(J)** Graphical presentation of ASPA+ Olig2+ cells in the corpus callosum. **(K-L)** Graphical presentation of N1 and P2 latencies at P48. n=3-9 animals per treatment group. P-values are calculated using one-way or two-way ANOVA with Tukey corrections and represent the effects of treatment groups and ages. Detailed p-values are provided in Tables S21 and S22. *p<0.05, **p<0.01 ***p<0.001, ****p<0.0001. Graphs are represented as Mean (SD). Associated Figures include S26-31.

### ASO-mediated suppression of Tubb4a restores the electrophysiological response

To assess whether the restored myelin is functional, we implanted 250 μm electrodes over the visual cortex through the skull. Visual evoked potentials (VEPs) are used to evaluate the visual pathways and serve as a biomarker of remyelination (35). We measured the latencies and amplitudes of the VEP peaks including N1, N2, P1, and P2 (**Figure 8G and 8K-L**). We observed a delay in N1 latency in untreated *Tubb4a^D249N/KO^* mice compared to PBS-injected WT and ASO-treated WT mice. Notably, N1 latencies but not N1 amplitudes are significantly restored in ASO-treated *Tubb4a^D249N/KO^* mice at P48, suggesting that the preserved myelin is functional at this age (**Figure 8G and 8K, Figure S30**). Interestingly, in untreated *Tubb4a^D249N/KO^* mice, the P2 peak appears shifted, and P2 latencies are delayed (**Figure 8L**). ASO treatment restored P2 latencies in *Tubb4a^D249N/KO^* mice (**Figure 8G and 8L**).

### Tubb4a ASO suppression fails to reverse cerebellar granule neuron (CGN) death and loss

Since the death of CGNs is a cell-autonomous deficit in H-ABC (14), we evaluated whether ASO-treated *Tubb4a^D249N/KO^* mice show any signs of cerebellar neuronal rescue. We co-stained CGNs using NeuN and cleaved caspase-3 to assess neuronal cell death and count. We failed to see rescue at P48 of CGN cell death and cell numbers in treated *Tubb4a^D249N/KO^* mice (**Figure S31A-C**), suggesting that ASOs fail to target CGNs via i.c.v. route. Our spatial uptake of ASO by cerebellum tissue supports this data (**Figure S15D**). To test if an alternate administration route targets CGNs, we injected FAM-labelled ASO 18-3 in pups (P1-P3) via the cisterna magna route. We perfused and collected brains 30 days after injection and performed immunostaining for FAM and NeuN on the cerebellar sections. We noted no ASO 18-3 distribution in CGNs, but there was good distribution in Purkinje neurons (**Figure S31D**). This indicates that CGNs resist ASO uptake even via the cisterna magna route.

## Discussion

There is an unmet need for effective treatment strategies for individuals affected by *TUBB4A*-associated disorder. This disorder includes a broad spectrum of phenotypes: early infantile encephalopathy, late infantile motor phenotypes including H-ABC, and adult-onset dystonia. A recurring D249N heterozygous pathogenic variant in *TUBB4A* is frequently present in H-ABC, which is characterized by progressive loss of neurological skills in early childhood and eventual death. *Tubb4a^D249N/KO^* mice show mild ataxia early (P18-P21) with motor function progressively worsening until their humane endpoint at P108-P110. Although motor deficits are not apparent during the juvenile phase (P21-P30), these mice exhibit early myelination deficits at P21. Furthermore, the disease pathology affects not only OL lineage cells but also leads to the loss of CGNs starting at P48. The slower progression of the disease in *Tubb4a^D249N/KO^* mice makes them well-suited for testing therapies targeting TUBB4A.

This study evaluated the germline and therapeutic efficacy of Tubb4a suppression. We characterized the *Tubb4a^D249N/KO^* mouse model, which mirrors severe H-ABC manifestations with a slower disease progression. Importantly, a single i.c.v. injection of *Tubb4a*-targeted ASO in *Tubb4a^D249N/KO^* mice successfully improved survival, motor function, and myelination deficits. Our findings demonstrate that a single ASO injection in this disease model can elicit a sustained phenotypic improvement compared to untreated cohorts, with ∼10-20% myelin restoration. Importantly, we also observed that ASO treatment restores *Mbp* transcripts in the treated H-ABC mice. In OLs, *Mbp* mRNA granules are transported via microtubules and translated locally in OL processes, indicating that the transport mechanisms are restored after ASO therapy (36, 37). Moreover, we observed a significant improvement in cellular response, evoked potential latencies, and overall functional measures in the ASO-treated mouse model. For the visual evoked potential response in H-ABC mice, we observed a hyperresponsive N1 peak compared to WT controls. This may be linked to the primary auditory cortex or a lack of surround inhibition in the cortex, resulting in a large response. Interestingly, auditory impairment occurs in individuals affected with H-ABC (38). Importantly, treated H-ABC mice show improvement in P2 latencies relative to untreated mice, which represents GABA-mediated inhibition (39).

H-ABC leukodystrophy involves cell-autonomous deficits in OLs, CGNs, and striatal neurons, making it difficult to develop a treatment that targets all these cell types. Our spatial data indicates that ASOs are well-distributed in OLs, cortical, and striatal neurons but fail to target CGNs. CGNs are particularly resistant to ASO uptake via the i.c.v. or intrathecal route in rodents and primates, and our data are consistent with other reports (40, 41). Mortberg et al. tested four different 2’-MOE ASOs (ASO 18-3 has 2’MOE modifications), and all the ASOs showed limited CGN uptake. Importantly, we demonstrated that CGNs are also hard to target, even with direct cisterna magna injections. It remains to be seen whether different chemical backbones (such as locked nucleic acids or phosphorodiamidate morpholino oligomers) or simply repetitive administrations via i.c.v. route can improve CGN targeting. This could limit the efficacy of ASO-based therapeutics in H-ABC. However, we observe substantial therapeutic effects in our mice, even with a single administration, which primarily restores myelination. Therefore, even partial cellular rescue in H-ABC may lead to significant clinical benefits as therapies that more comprehensively target affected cells are developed. The ASO treatment might also offer improved therapeutic benefits in other *TUBB4A*-related disorders that only exhibit isolated hypomyelination and/or basal ganglia atrophy.

*TUBB4A* variants are monoallelic in affected individuals. The heterozygous *Tubb4a ^D249N/+^* murine model exhibits mild phenotypic effects with no behavioral abnormalities, making it less ideal for studying disease endpoints after ASO treatment. We also tested ASO treatment in *Tubb4a^D249N/D249N^* mice. These mice exhibit limited rescue, possibly because ASOs fail to target CGNs, and more than 60% of CGNs die in *Tubb4a^D249N/D249N^* mice by the end stage. Therefore, this model might underestimate the importance of the treatment benefits of this ASO drug. As a result, we mainly studied ASO treatment in *Tubb4a^D249N/KO^* mice, which more closely resemble the human heterozygous disease than homozygous *Tubb4a^D249N/D249N^* mice.

Several important questions remain, including whether repeated ASO treatments will lead to sustained improvement in pre-symptomatic mice. Future work will require determining the best dosing route, frequency, and detailed characterization of ASO pharmacokinetics and pharmacodynamics within the CNS. Notably, this Tubb4a anti-sense therapy is not specific to the recurring D249N variant and is suitable for broader applications in *TUBB4A*-associated leukodystrophy. Recent data indicate that germline deletion of *Tubb4a* in rodents recruits other tubulins, resulting in no abnormal phenotype (14). Further studies are necessary to understand how ASO influences and alters the β-tubulin composition. Mice treated with ASO 18-3 exhibited astrogliosis and microgliosis, indicating an activated immune response in the brain. The FAM-labeled 18-3 ASO was injected at a lower dose (10 µg/g) and achieved a strong OL distribution; it remains to be seen whether we can reduce the ASO dose or if alternative ASO designs can demonstrate similar disease mitigation without causing astrogliosis or microgliosis. Finally, testing ASO in additional models of milder and more severe *TUBB4A*-associated leukodystrophy with different pathogenic variants would help verify the general applicability of these findings.

Overall, Tubb4a ASO treatment effectively downregulates Tubb4a in H-ABC mouse models. Our study shows that although it does not target CGNs, ASO treatment maintains myelination, enhances motor and functional outcomes, and reverses many aspects of the disease phenotype. Our data indicate that Tubb4a ASOs are a promising therapeutic approach for H-ABC and *TUBB4A*-related leukodystrophy.

## Methods

### Sex as a biological variable

Our study included male and female findings, and similar findings were reported for both sexes.

### Animals

All animal protocols were approved by the Institutional Animal Care and Use Committee (IACUC) at the Children’s Hospital of Philadelphia. Equal number of females and males are included in this study. All mice were generated on the C57/BL6J background. *Tubb4a^KO/+^* mice (KOMP consortium) were intercrossed to obtain *Tubb4a^KO/KO^*. *Tubb4a^KO/+^* model was crossed with *Tubb4a^D249N/+^* to obtain *Tubb4a^D249N/KO^*mice. F1 litters were used for experiments. All animals were maintained in a temperature-controlled room, with a 12-hour light on/off cycle and free access to food and liquid. For genotyping, tail snips were collected and outsourced to TransnetYX, Inc (Cordoba, TN).

### Oligonucleotides design and synthesis

The ASO library was designed against the full mouse Tubb4a mRNA sequence. After ASO *in silico* modeling, the number of phosphorothioate bonds were modified to reduce potential toxicity and obtain multiple designs from the parent molecule. All ASOs were synthesized at BioSpring (BioSpring, Germany) and suspended in aliquots containing Phosphate buffer saline (PBS; Thermo Scientific, 1001-023) and then stored at −20 °C or −80°C.

### In vitro ASO screening

ASO *in vitro* screening was carried out using primary mouse cortical neurons. Embryos were removed at E15.5 from WT C57BL/6 females. Cortical tissue of each embryo was dissected on ice-cold Hank’s Balanced Salt Solution (Gibco, 14170-112). Pooled tissue was minced and digested with 0.05% Trypsin-EDTA (Gibco, 25300-054) at 37°C for 12 min. Cells were triturated, resuspended in modified neurobasal media [Glutamax (Gibco, 35050-061), 2% penicillin/streptomycin (Gibco, 15140-122), and B27 (Gibco, 17504-044)], and seeded at 0.25 × 10E6 cells/well in 6-well plates pre-coated with poly-L-lysine (Sigma, P1274-500MG). Neurons were treated with ASO doses on *in vitro* day 7 after culture and collected 7 days after treatment. The toxicity of each ASO was determined using the ApoTox-Glo triplex Assay (Promega, G6320) as per manufacturer instructions.

### Stereotactic i.c.v. injections. For Pups (P1-2)

Mouse pups were cryo anesthetized on ice on a cooled stage of a Digital Just for Mouse Stereotaxic Instrument (Stoelting, 51730D). The head of the anesthetized pup was securely placed flat between ear cups to ensure a level head position for injection. ASO (weight-based) or PBS (2µL) were loaded into a Hamilton 1700 syringe (Hamilton Company, CAL7635-01) with a 32-gauge, 0.5-inch needle (Hamilton Company, 7803-04) and secured by the Quintessential Stereotaxic Injector (QSI; Stoelting, 53311). The needle tip is aligned with the pups’ lambda suture, which is visible through the skin in neonates. This lambda point was used as the reference point (“zero”) to target the lateral ventricles: e.g., for right-sided injection; the syringe was moved to +1.50mm (anterior); −0.80mm (right); and −1.50mm (deep). Injections were pushed at a rate of 0.5µL/min. The needle was slowly retracted to avoid material leakage.

### Cisterna magna injection. For pups (P1-3)

Mouse pups were cryo-anesthetized in a petri dish surrounded by ice until all movements stopped. The head of the anesthetized pup was securely placed between the administer’s fingers to ensure a leveled head and neck were in position for injection. ASO FAM and ASO NTC (weight-based) or PBS (2µL) were loaded into a Hamilton 1700 syringe (Hamilton Company, CAL7635-01) with a 32-gauge, 0.5-inch needle (Hamilton Company, 7803-04). The needle tip was aligned with the pup’s neck below the skull; then the needle angle was increased to 50-60° to insert in the cerebrospinal fluid. The needle was slowly retracted after holding it in position for 60 seconds to avoid material leakage. After the injection pups were allowed to recover on a heat pad completely and returned to the cage.

### RNA extraction, qRT-PCR and Nanostring

To determine the expression of Tubb4a from mouse brain, tissues were dissected and snap frozen. RNA extraction, c-DNA, qRT-PCR and Nanostring details are provided in Supplemental Methods.

### Histology

Mice were anesthetized and transcardially perfused with an initial flush of PBS (Thermo Scientific, 1001-023) followed by 4% paraformaldehyde (PFA). Anesthetic methods varied by age with isoflurane gas/oxygen flow for adolescent mice (>P8) or a weight-based injection of a xylazine, ketamine, and PBS mixture for adult mice (>P21). The harvested whole brain tissue was post-fixed with 4% PFA in PBS overnight at 4°C, then rinsed in PBS and transferred to 30% sucrose in PBS at 4°C until the sample was no longer buoyant (>3 days), indicating penetrance of the sucrose solution. The whole brain was then embedded in a Tissue-Tek Optimum Cutting Temperature compound (Sakura, 4583) and cryosectioned at 20µm by microtome (Leica, CM3050S) into coronal sections. These sections were stored in free-floating PBS at 4 °C and transferred to a freezing medium composed of a mixture of DMEM containing HEPES (Gibco, 11885084 and Sigma, H7523, respectively) and glycerol (Sigma, G6279-500) for long-term storage at −20 °C. For myelin staining, Eriochrome Cyanine (Eri-C) staining was conducted using previously established methods (10). Sections were imaged by Leica CTR6000/B microscope and ImageJ software was used to quantify myelin levels in the corpus callosum and cerebellum (10).

### Myelin purification

The myelin fraction was extracted as published previously (42). Myelin was purified from the brains of P35-40 and P108-110 mice. Frozen whole forebrains were homogenized in 0.32M sucrose-containing protease inhibitor (Roche) using a TissueLyser II machine (Qiagen, 85300). After homogenizing the brains in 0.32LM sucrose solution, a crude myelin fraction was obtained by density gradient centrifugation over a 0.85LM sucrose. After washing and two osmotic shocks, the final myelin fraction was purified by second sucrose gradient centrifugation. Myelin fractions were washed and suspended in 1X Tris-buffered saline (137 mM NaCl, 20 mM Tris-HCl, pH 7.4, 4 °C) supplemented with protease inhibitor (Roche). The myelin fractions were then stored at −80°C until use.

### Electron microscopy

Mice were anesthetized with a weight-based injection of a xylazine, ketamine, and PBS mixture and transcardially perfused with an initial flush of PBS followed by a fixative solution composed of 2.5% glutaraldehyde, 2.0% paraformaldehyde in 0.1M sodium cacodylate buffer at pH 7.4. Optic nerve, corpus callosum, and cerebellar tissue were bluntly dissected and stored in the fixative solution. After overnight fixation at 4°C, tissues were washed with buffer and post-fixed in 2.0% osmium tetroxide for 1 hour at RT. Tissue was then rinsed in dH2O before en-bloc staining with 2% uranyl acetate and dehydrated through graded ethanol. The tissue was embedded in EMbed-812 (Electron Microscopy Sciences), and thin sections were stained with uranyl acetate and lead citrate. Sections were examined with a JEOL 1010 electron microscope fitted with a Hamamatsu digital camera and AMT Advantage NanoSprint500 software. The images for EM sections were assessed using ImageJ software. The inner and outer axonal area was measured for g-ratio analysis and quantified in 50 axons per animal with n = 3 per group.

### RNAscope

Mouse brains were harvested by standard perfusion, including overnight post-fixation, followed by 30% sucrose in PBS incubation for 3 days at 4°C, followed by paraffin embedding and sagittal sectioning. For Chromogenic InSitu Hybridization (CISH) staining, fresh slides were sectioned, air-dried, and baked within 48hrs of staining. Staining was performed on a Bond RXm automated staining system (Leica Biosystems). For CISH probe staining, a Mm-Tubb4 probe (Advanced Cell Diagnostics, 590508) was used along with RNAscope 2.5 LSx Reagent Kit Red (Advanced Cell Diagnostics, 322750) using standardized protocols from Advanced Cell Diagnostics. For Fluorescent InSitu Hybridization (FISH) staining, fresh slides were sectioned, air-dried, and baked within 48 hours of staining. Staining was performed on a Bond RXm automated staining system (Leica Biosystems). For FISH probe staining, Mm-Myrf and Mm-Mbp probes (Advanced Cell Diagnostics, 524068, 451498-C2) were used along with the RNAscope LS Multiplex Fluorescent Reagent Kit (Advanced Cell Diagnostics, 322800) using the Vivid 520 and Vivid 650 fluorophores. Standardized protocols from Advanced Cell Diagnostics were used except for the VIVID fluorophore dilutions, which were used at a 1:5K dilution.

### Immunoblotting

Brain tissues were dissected to protein levels in different brain regions (including cortex, striatum, and cerebellum) at different ages. Immunoblotting for myelin fraction and protein extracts was performed as explained in the Supplemental Methods.

### Immunofluorescence and image analysis

Brain sections were washed in PBS, blocked in 10% Normal Goat Serum (EMD Millipore, S26), 2% Bovine Serum Albumin (Sigma, A7030-100G), and 0.1% Triton X-100 (Sigma, T-9284) in PBS for 1 hour at room temperature (RT) and incubated in primary antibodies (See **Table S5**) overnight at 4°C. Following this incubation, sections were washed with 0.1% Triton X-100 (Sigma, T-9284) in PBS, incubated in fluorescent secondary antibodies for 1 hour, and protected from light at RT. In this study, AlexaFluor conjugated secondary antibodies were used at 1:1000 dilution. Sections were sequentially washed with 0.1% Triton X-100 (Sigma, T-9284) in PBS and mounted onto Superfrost Plus Microscope Slides (Fisher, 12-550-15) using ProLong Gold Antifade Mountant with DAPI (ThermoFisher, P36931).

### IHC analysis

Sections were imaged using a confocal microscope (Leica) or a Keyence BZ-X810 microscope. Neurons (NeuN and/or CTIP2-labelled neurons) and OLs (ASPA+ or Olig2+) were quantified using ImageJ or Keyence BZ-X800 Analyzer software. Eri-C quantification was conducted by measuring the ratio of myelinated area to total white area as published previously (10).

### RNA scope analysis

Colorimetric or fluorescent RNA scope images were imaged and stitched at 10X magnification. For measuring *Mbp* mRNA intensity, images of RNAscope sections were captured at 40X, and *Mbp* intensity was measured using ImageJ, as published previously.

### Striatal volume analysis

Eriochrome cyanine (Eri-C) stained 50 µm thick sections were analyzed for striatal volume measurement. WT, *Tubb4a^KO/KO^*, *Tubb4a^D249N/KO^*, *Tubb4a^D249N/+^*, *Tubb4a^D249N/D249N^* mice brain sections were analyzed at P32-P37 (end-stage of *Tubb4a^D249N/D249N^*) and P108-P110 (end-stage of *Tubb4a^D249N/KO^*). For stereological analyses, n=3 animals were chosen with 2-3 sections per brain of the same region rostro-caudally to eliminate bias. The striatum’s boundary included the corpus callosum border, and the external capsule was traced to its furthest ventral extent. The line then tracked to the inferior point of the lateral ventricle, omitting the anterior commissure, following along the lateral ventricle, and rejoining the corpus callosum (43). The area was quantified using ImageJ, and volume was analyzed, considering its thickness (50µm).

### Behavioral Assays

Rotarod, tremor assay, grip strength, and open field were conducted as published previously (10, 44, 45). The Supplemental method section includes the methods of behavioral assays.

### Electrode implantation surgeries; evoked potential stimuli presentation and data analysis

*Electrode Implantation, Evoked Visual and Auditory Stimuli:* Electrode implantation was performed on mice (46, 47) (See detailed method in Supplemental material)

### Figure preparation and statistical analysis

All figures are prepared using Adobe Photoshop, and all fluorescent images are subjected to exposure “1” to enhance the brightness. The mean at each time point for each mouse was obtained across three trials for behavior (rotarod and grip strength) analysis. All data are presented as the mean (SD) in figures. For mouse studies, ‘n’ represents the number of animals used per experiment unless indicated otherwise. Rotarod, tremor, and grip strength were evaluated in the *Tubb4a^D249N/KO^* mouse model for characterization and ASO treatment using mixed-effects analysis with random intercept model and linear assumption; separate models were fit prior to end-stage and post end-stage due to the discontinuation of the untreated, affected subgroup. Null hypotheses are that the estimate of each fixed effect (continuous time in days from P30, dichotomous genotype, dichotomous treatment, and interaction terms) in association to the behavior outcome is 0. Likelihood ratio testing was used to determine model fit. Estimated mean and 95% CI are reported for each subgroup from the model. Kaplan-Meier curves with log-rank testing were used to compare survival between groups. The null hypothesis for the log-rank test is that the probability of survival over time is the same for all subgroups. Median survival time is reported with 95% confidence intervals where applicable. Cox regression was used to evaluate the association of ASO treatment on the hazard of survival within *Tubb4a^D249N/KO^* mice. Graphical methods, such as log-log curves and Schoenfeld goodness-of-fit testing, were used to test the proportional hazards assumption. The null hypothesis is that hazard ratio for treatment is 1 or there is no difference in the risk of survival between treatment groups. Hazard ratio(s) of the model are reported with 95% confidence intervals. Comparisons in myelin quantification, NeuN, ASPA, NG2, Olig2 and cleaved caspase-3 counts, and fluorescent intensity were analyzed by mixed effects two-way ANOVA with multiple comparisons post-hoc Tukey tests. EM analysis for g-ratio and myelin analysis (Eri-C quantification) was conducted through two-way ANOVA, followed by a post-hoc Tukey test. Dose-response curves were calculated by Motulsky (4-parametric logistic), and non-linear regression, and the estimated effective concentration (EC) at 50% with 95% confidence intervals was reported. All statistical analyses were performed using Prism 7.0 (GraphPad Software) or R (version 4.4.0), with two-sided testing and an alpha level of 0.05 for statistical significance.

## Data and code availability

Data will be made available upon request.

## Acknowledgements

We would like to thank Dr. Judith B. Grinspan for providing hybridoma antibodies, the CHOP Pathology Core for technical support of RNAscope, and Elena Lysenko from Dr. Beverly Davidson’s laboratory for technical support for the tremor assay.

This study was supported by funds from Synaptix Bio company; Kamens Chair in Translational Neurotherapeutics; H-ABC Foundation UK; and Commonwealth Universal Research Enhancement Program (CURE) funding (SAP # 4100077047).

## Author Contributions

SS, JLH, and AV designed the study and drafted the manuscript. SS, JLH, PN, and AB conducted experiments, performed the analysis, and analyzed the data. SS and AV supervised, conceptualized, and coordinated the study. ADA conducted Nanostring, and AT provided her expertise and data analysis for Nanostring. SW helped in the statistical analysis of the data. MAJ provided help with the myelin extraction assay. JLH conducted and analyzed data for *in vivo* electrophysiology; EDM provided his expertise. SS, JLH, PN, EDM, and AV wrote and reviewed the manuscript.

## Declaration of generative AI and AI-assisted technologies

We have not used generative AI and AI-assisted technologies to write the manuscript. Grammarly was used to fix grammatical errors.

**Video 1** represents treated *Tubb4a^D249N/KO^*mice (blue label; Age: P134) versus untreated *Tubb4a^D249N/KO^* mice (yellow label; Age: P104).

**Video 2** represents treated *Tubb4a^D249N/KO^*mice (no label; Age: P128) versus untreated *Tubb4a^D249N/KO^* mice (blue label; Age: P97).

