## Supplemental Materials for "Therapeutic suppression of *Tubb4a* rescues H-ABC leukodystrophy"

**Supplemental Methods:**

**RNA extraction, qRT-PCR and Nanostring.** Approximately 50mg of mouse tissue was placed in 96-well plates (RNeasy 96 Universal Tissue Kit, Qiagen, 74881), placed in QIAzol Lysis Reagent (Qiagen, 79306), and disrupted by a TissueLyser II machine (Qiagen, 85300). Homogenized tissues were purified and eluted with RNAse Free Water. cDNA was generated from extracted RNA (High-Capacity RNA-to-cDNA Kit, Thermo Fisher) using Veriti 96-Well Thermal Cycler (Thermo Fisher). cDNA was mixed with TaqMan Gene Expression Master Mix (Thermo Fisher), Prime Time qPCR Assay Tubb4a and SFRS9 (Integrated DNA Technologies, Mm.PT.56a.9905332 and Mm SRFS9 455-588, respectively). Replicates of this solution were placed in a 384-well plate for QuantStudio FLEX 7 qRT-qPCR (Thermo Fisher). Mouse tissue RNA, 500ng, were hybridized with the nCounter CodeSets (nanoString, Seattle, USA) to a custom-designed Mouse Tubb4a panel, prepped on nCounter FLEX prep station (nanoString) and scanned the image on nCounter FLEX digital analyzer (nanoString). Raw counts were normalized to the internal positive controls and 4 housekeeping genes (Alas1, Hprt, Ppib and Tbp) by following the company's instruction.

**Immunoblotting.** The brain tissues were lysed in RIPA buffer (Thermo Fischer Scientific, USA) in the presence of protease and phosphatase inhibitors (Sigma-Aldrich, USA) and diluted with Laemmli buffer (Biorad, USA) prior to western blotting (10). The primary antibodies used for western blot are listed in Table 2. Images were scanned and analyzed by ImageJ software. For normalization with loading control, mouse anti-vinculin (1:3000, Sigma, Cat: V9131) was used after stripping the membrane (Thermo Fischer Scientific, USA). Protein molecular weight markers (Bio-Rad Precision Plus Protein™ Kaleidoscope™ Prestained Protein Standards, USA) was included in each blotting.

**Behavioral assays.** For all behavioral assays, mice underwent a period of acclimation for 30 minutes before beginning.

*Rotarod:* Rotarod assay was performed as published previously (10).

*Grip strength:* Forelimb and hindlimb grip strength of mice were tested using a grip strength meter (080312-3 Columbus Instruments, Columbus, OH, USA) (32). For forelimb testing, mice were held and allowed to grasp a horizontal metal bar with both paws. The mice were then steadily pulled away, and the pull force was recorded once the mice unclasped the metal bar. For the hindlimb grip strength measurements, mice were allowed to grab the horizontal bar with their hindlimb paws, and mice were pulled after a firmer grip of hind paws until their grasp broke. Measurements were not considered if the mice failed to use both fore- and hind paws turned backward during the pull or broke their grasps without resistance. Three consecutive trials for the forelimb strength and three for the hindlimb paws were conducted. The averaged values for the fore- and hindlimbs were taken for further analysis.

*Tremor assay:* Tremor assay was performed using San Diego Instruments' Startle Response System (SR-LAB) cabinets. Mice were placed in plexi-glass cylinders within these enclosed cabinets for a total of 8 minutes, which included 3 minutes of acclimation to their new environment and 5 minutes of recorded testing. Tremorous activity was recorded as signal intensities in dBVs (San Diego Instruments' Tremor Monitor Software). A standardized subset of measurements was averaged to compare tremorous phenotype between mouse genotypes and treatment groups.

*Open Field*: ANY-Maze software was used for all video recordings and data acquisition. In a 16-square open box, the four innermost boxes comprised the 'inner zone', while the 12 surrounding boxes comprised the 'outer zone'. Dimensions of the box were aligned and corrected at the beginning of each testing session, with background pictures taken between each mouse to ensure zone areas were kept consistent. Mice were allowed to freely explore their open environment for 10 minutes. The inner surfaces of the box were wiped down with 70% ethanol between each mouse to remove any confounding scents of previous mice. Data from ANY-Maze was analyzed to assess total distance traveled and maximum speed of ambulation.

**Electrode implantation and evoked potential measurements**

Animals were placed in custom-built cages and allowed to acclimate to the dark for 30 minutes prior to recording. The mice were presented with a 1Hz light flicker stimulus for 2 minutes generated by an Arduino Uno Board connected to an Atmel Atmega8/168/328 AVR microcontroller and FTDI FT232 breakout board (SparkFun Electronics). Light intensity was measured with a light meter model 840006C (Sper Scientific). After a 5-minute rest period, the mice were presented with 82Db, 15ms white noise stimulation train for 17 minutes generated by WaveForm Generator model 2414A (Tegam). Decibels were measured from the center of four cages using Sound Meter App version 1.7.5 SM-J337P, US (Smart Tools Co.). The evoked potentials were recorded using the 32-channel Intan extracellular amplifier.

*Analysis of Evoked Potentials:* Raw EVP data was imported into MATLAB R2021b (Mathworks Inc., Natick, MA) via read_Intan Gui (Intan Technologies, Los Angeles, CA). The EEG and EVP data underwent artifact rejection algorithm removing signal with root mean square amplitude of >200 or <30 and skew >0.4. For auditory and visual potential analysis, the leads over the auditory and visual cortex were averaged using in-house written scripts in Matlab that averaged the signals based on a TTL pulse generated with each stimulus. The average stimuli were presented via a marking GUI in Matlab, and a scorer marked regions around the N1, P1, and N2 peaks and the software chose the highest peak and measured amplitude and latency for both left and right auditory or visual channels for each mouse. The latencies were averaged for the left leads for both auditory and visual stimuli. For electrophysiology, the averaged latencies were plotted in MATLAB R2021b, and statistical analysis was conducted.

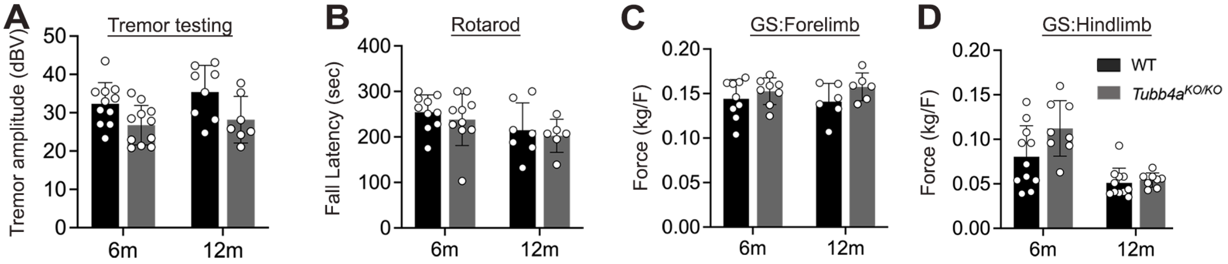

**Supplemental Figure 1** *Tubb4a*$^{KO/KO}$ motor behavior assays relative to WT at 6 and 12 months. **(A)** Tremor amplitude, **(B)** Rotarod, **(C)** Forelimb grip strength, and **(D)** Hindlimb grip strength. Repeated measures ANOVA was used to assess the motor deficits. Data is presented as Mean (SD). n=7-11 mice per genotype. Non-significant values are not denoted. See Supplemental Table 9 for a detailed statistical analysis of genotype and age interactions.

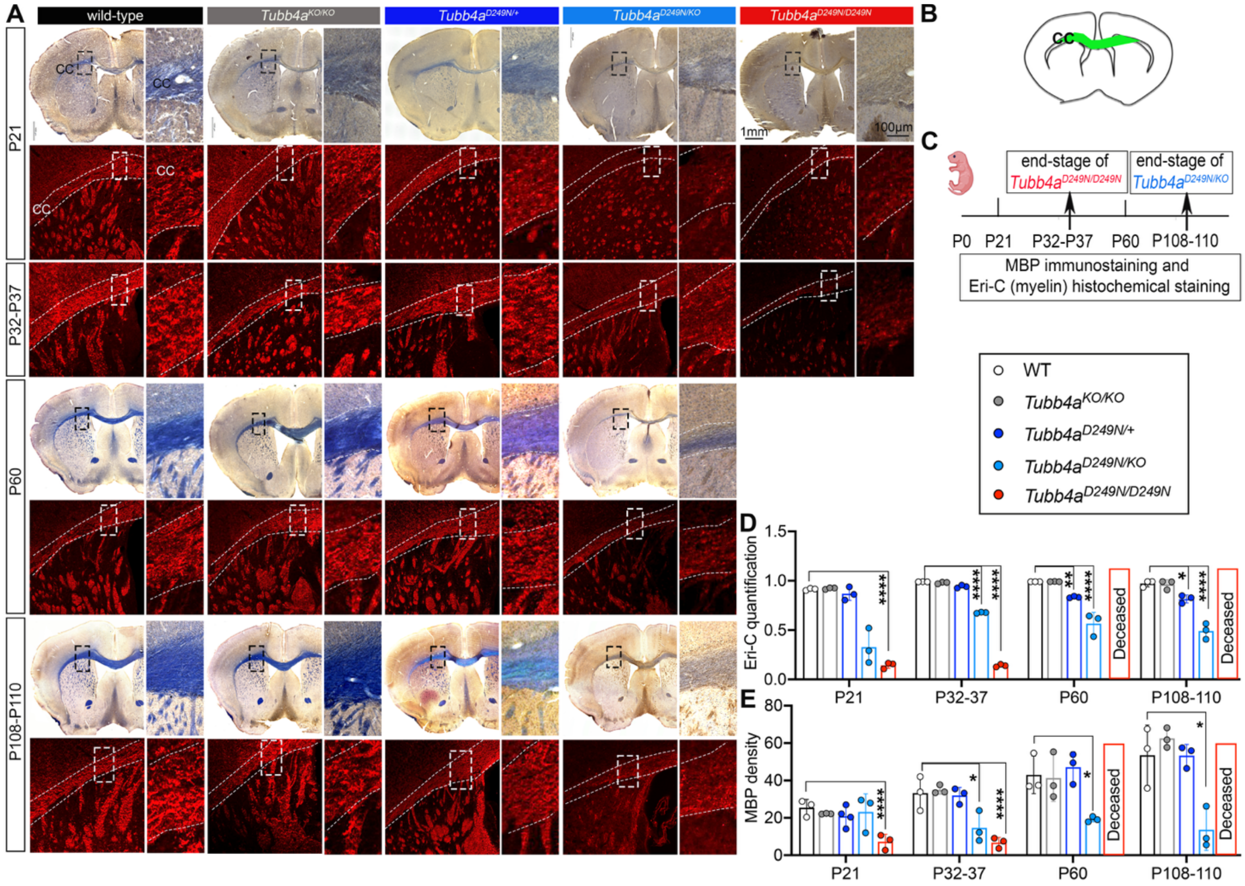

**Supplemental Figure 2** Myelin analysis in corpus callosum across all genotypes. **(A)** Representative histochemical images of Eri-C (blue; tiled images) and MBP (red) at P21, P32-P37, P60, P108-P110 (Eri-C images of P32-P37 are provided in Figure 1). **(B)** Coronal section of brain highlighting corpus callosum region that was used for analysis. **(C)** Survival timeline of *Tubb4a* mutant mice. **(D)** Graphical presentation of MBP density. **(E)** Graphical presentation of MBP density ≈. n=3-4 mice per genotype. Mixed effects two-way ANOVA was conducted to analyze the statistical significance. Data is presented as Mean (SD). Significant values are provided compared to WTs. Other age and genotype interactions are provided in the Supplemental Tables. *p<0.05, **p<0.01 ***p<0.001, ****p<0.0001. Non-significant values are not denoted. See Supplemental Table 11 for a detailed statistical analysis of genotype and age interactions.

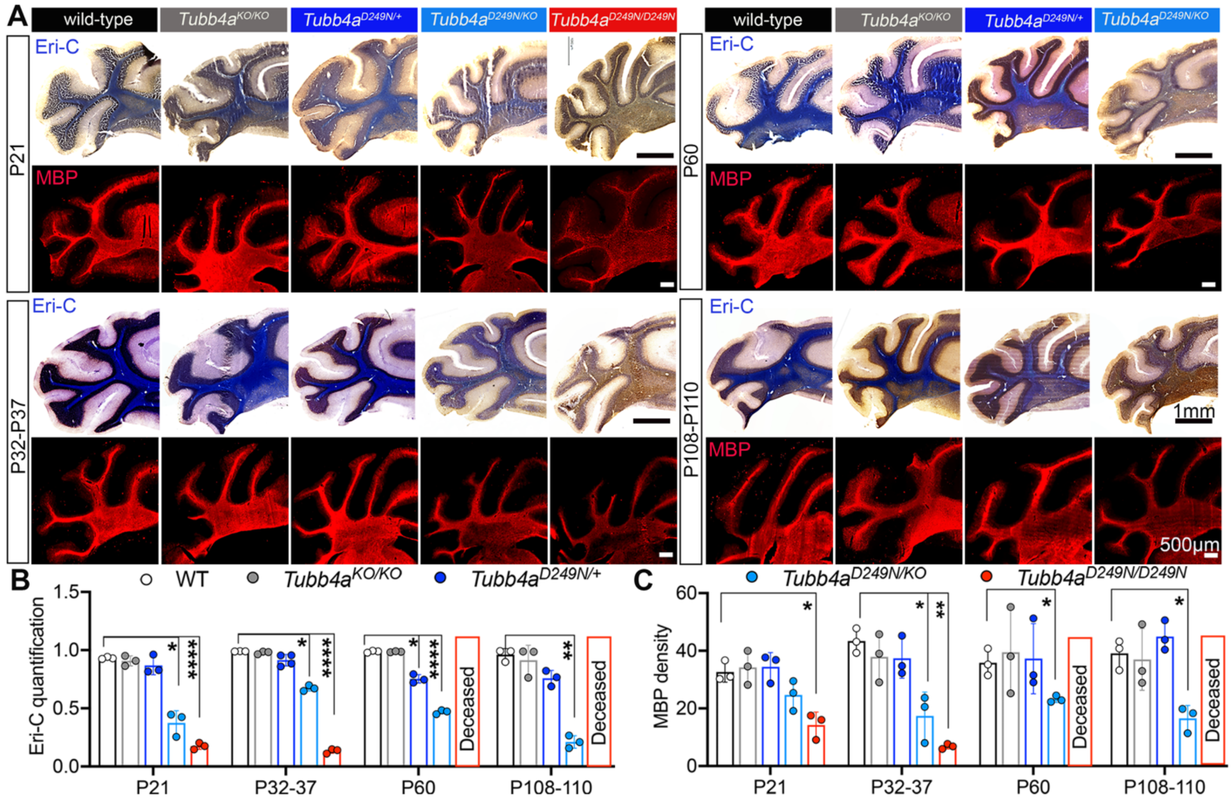

**Supplemental Figure 3** Myelin analysis in cerebellum across all genotypes. **(A)** Representative tiled histochemical images of Eri-C (blue) and MBP (red) at P21, P32-P37, P60, P108-P110. **(B-C)** Graphical presentation of Eri-C quantification (B) and MBP density (C). n=3-4 mice per genotype. Mixed effects two-way ANOVA was conducted to analyze the statistical significance. Data is presented as Mean (SD). Significant values are provided compared to WTs. Other age and genotype interactions are provided in Supplemental Table 12. *p<0.05, **p<0.01 ***p<0.001, ****p<0.0001. Non-significant values are not denoted.

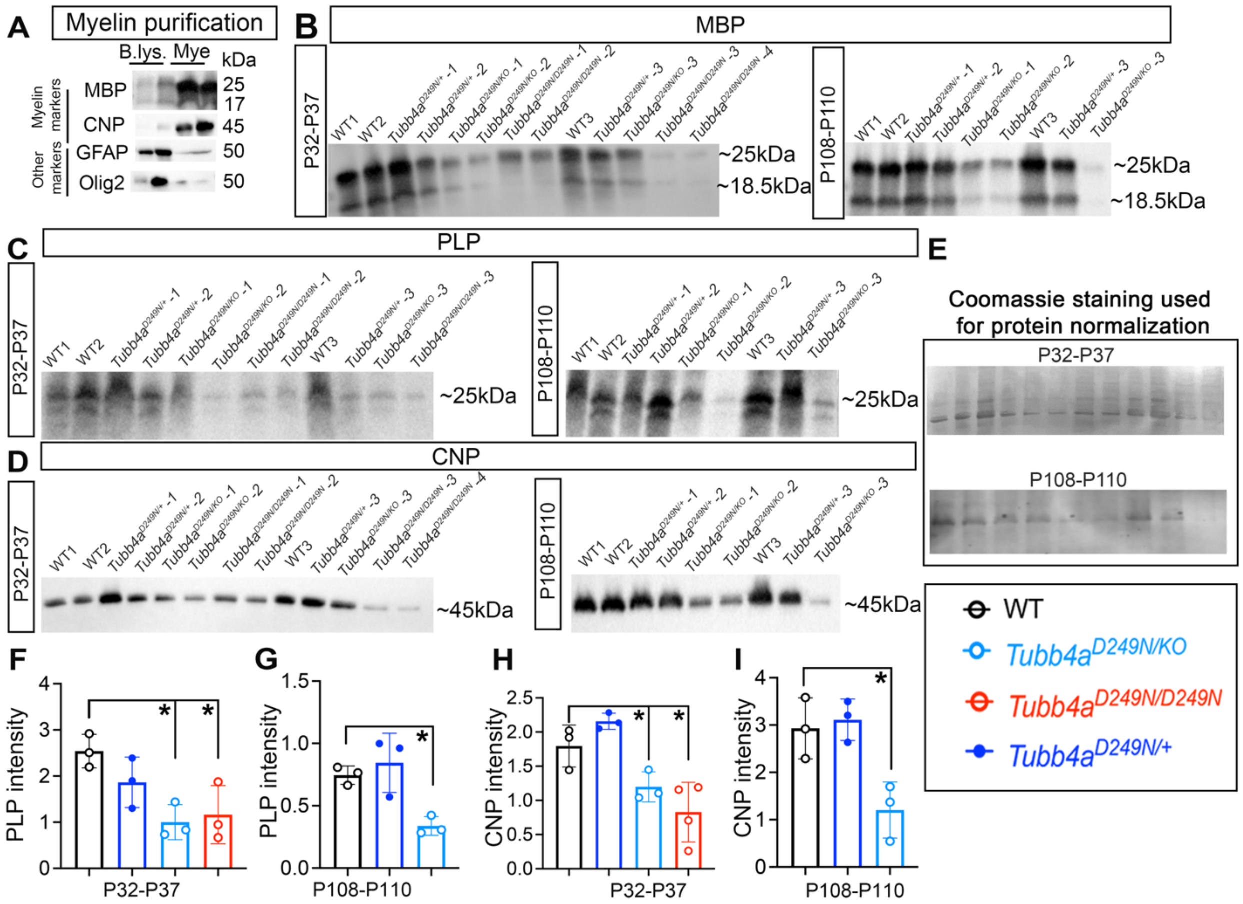

**Supplemental Figure 4** Immunoblotting analysis of extracted myelin fraction. **(A)** Images showing the successful purification of the myelin fraction (enriched with myelin proteins MBP and CNP). B.lys. = Brain lysate and Mye = Myelin fraction. **(B-D)** MBP (B; MBP graph is represented in Figure 1L), PLP (C), and CNP (D) immunoblotting images of pure myelin fraction at P32-37 and P108-P110. **(E)** Representative images of Coomassie staining that were used for protein normalization. **(F-G)** Plots of PLP band intensity of P32-37 and P108-P110, respectively. **(H-I)** Plots of CNP band intensity of P32-37 and P108-P110, respectively. n=3-4 mice per genotype and age. One-way ANOVA was carried out to analyze the statistical significance. Data is presented as Mean (SD). Significant values represent WT comparisons. See p-values in Supplemental Table 10. Non-significant values are not denoted. *p<0.05, **p<0.01 ***p<0.001, ****p<0.0001.

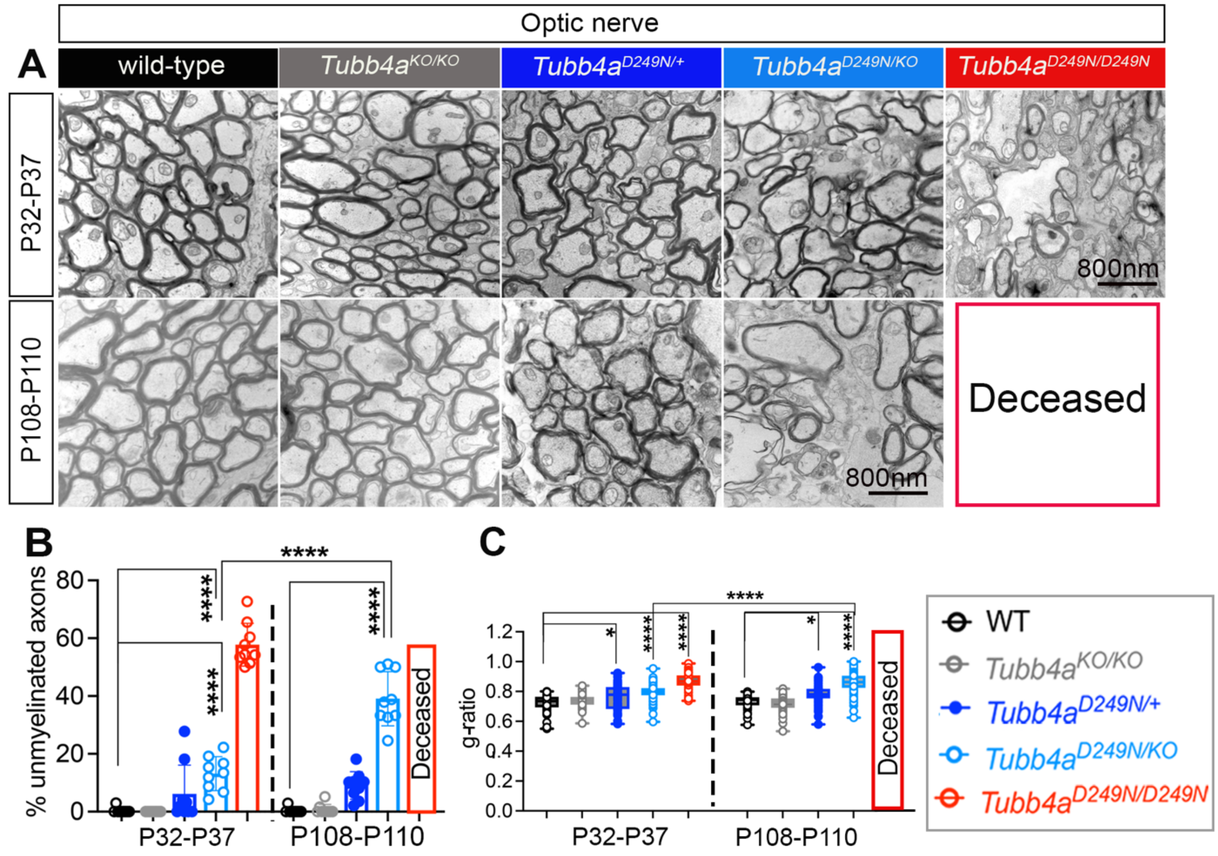

**Supplemental Figure 5** Myelin quantification at high resolution in optic nerve. **(A)** Representative electron micrographs of P32-P37 (end-stage of *Tubb4a*$^{D249N/D249N}$) and P108-P110 (end-stage of *Tubb4a*$^{D249N/KO}$). **(B)** Graphical presentation of % unmyelinated axons and **(C)** Graphical presentation of g-ratio at P32-P37 and P108-P110 in optic nerve. n=3 animals per genotype, with 40-60 axons per animal. Mixed effects two-way ANOVA was conducted to analyze the statistical significance. Data is presented as Mean (SD). Significant values are provided compared to WTs. Other age and genotype interactions are provided in the Supplemental Table 8. *p<0.05, **p<0.01 ***p<0.001, ****p<0.0001. Non-significant values are not denoted.

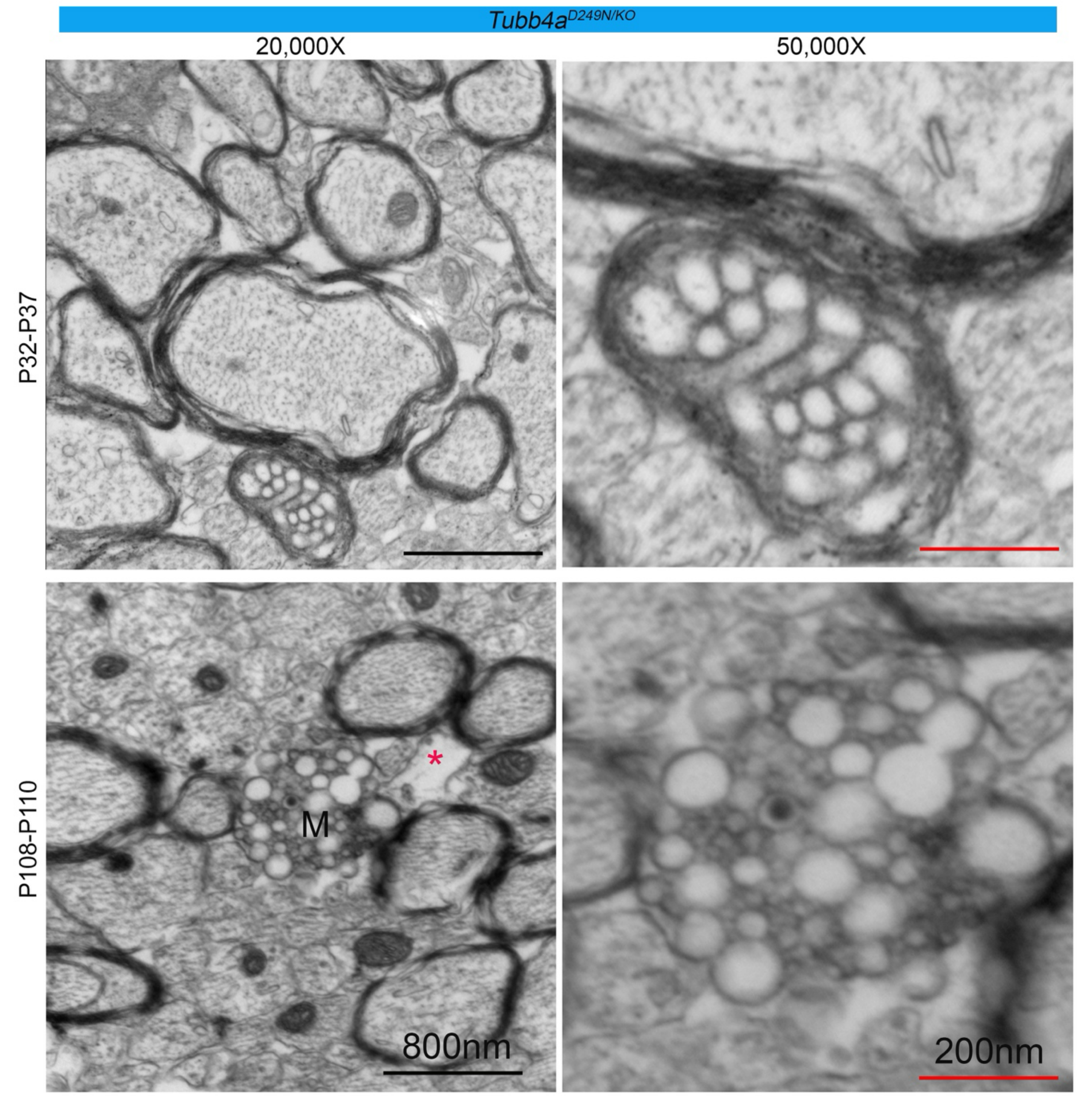

**Supplemental Figure 6** Representative electron micrograph images at P32-P37 and P108-P110 of corpus callosum. M- Phagocytosed myelin debris; * denotes the vacuole.

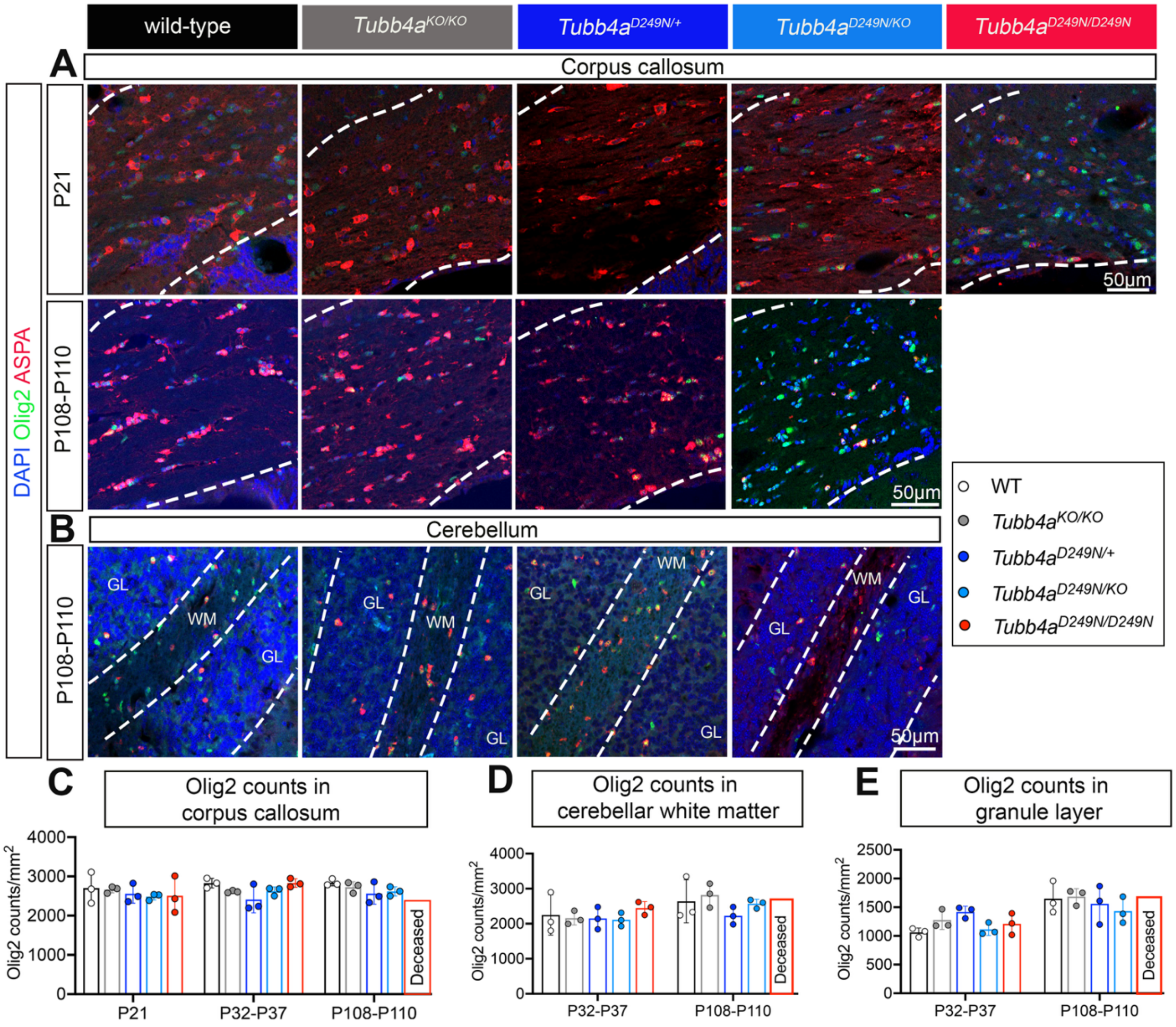

**Supplemental Figure 7** OLs count at different ages across all genotypes. (**A**) Representative images of ASPA+ (red), Olig2+ (green) and DAPI (blue) cells at P21 and P108-P110 in the corpus callosum (Representative images of P32-P37 are available in Figure 2). **(B)** Representative images of ASPA+ (red), Olig2+ (green) and DAPI (blue) cells at P108-P110 in the cerebellum (Representative images of P32-P37 are available in Figure 2). **(C-E)** Olig2 profile/mm$^2$ at different ages in the corpus callosum, cerebellar white matter and granule layer. n=3 mice per genotype and age. Mixed effects two-way ANOVA was carried out to analyze the statistical significance. Data is presented as Mean (SD). See Supplemental Table 14 for the p-values. Non-significant values are not denoted.

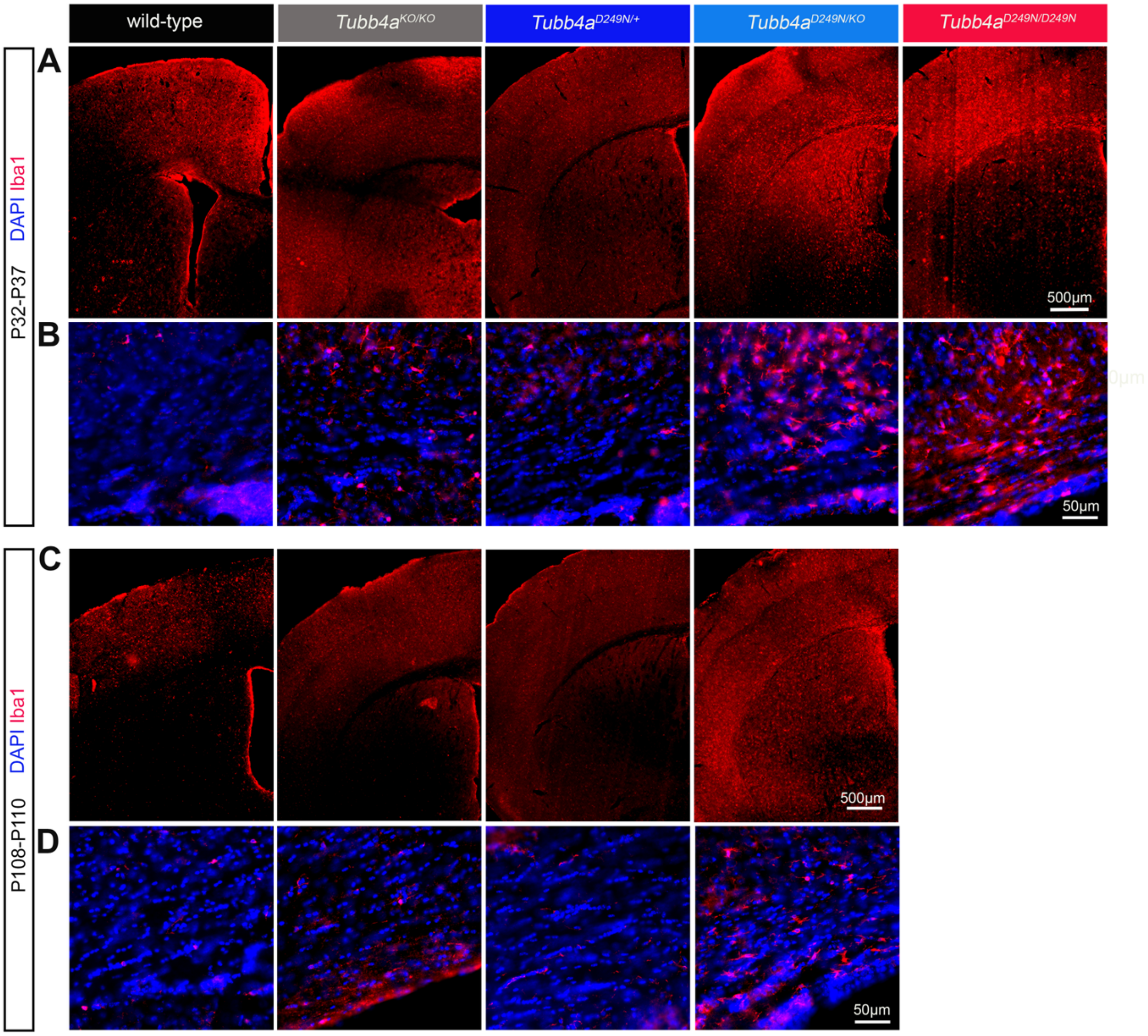

**Supplemental Figure 8** Microglia immunostaining (Iba1) of the forebrain and corpus callosum. **(A-B)** Representative images of Iba1 immunostaining at 4X and 20X at P32-P37; all sections are co-stained with GFAP from Figure 2E, and Keyence software was used to overlay the respective channels. **(C-D)** Representative images of Iba1 immunostaining at P108-P110; all sections in this image panel are co-stained with GFAP from supplemental figure 9A, and Keyence BZ software was used to overlay the respective channels.

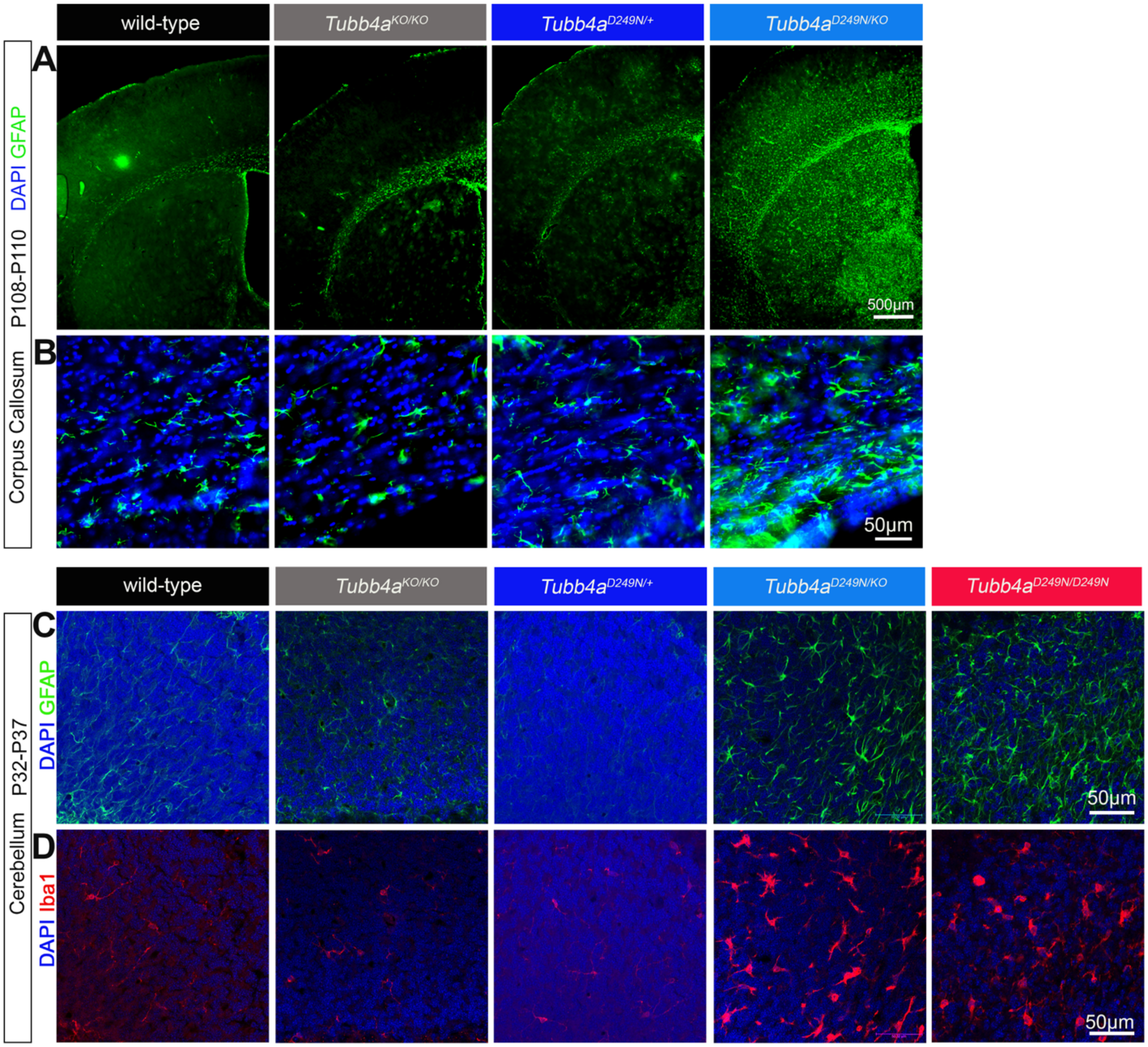

**Supplemental Figure 9** Astrocytes (GFAP) and microglia (Iba1) across all genotypes in the forebrain and cerebellum. **(A-B)** Representative images of GFAP immunostaining at low and high magnification at P108-P110; all sections in this image panel are co-stained with Iba1 from supplemental figure 8C, and Keyence BZ software was used to overlay the respective channels. **(C-D)** Representative GFAP (C) and Iba1 (D) immunostaining images of the same tissue section in each genotype at P32-P37 in the cerebellum and ImageJ software was used to overlay the respective channels.

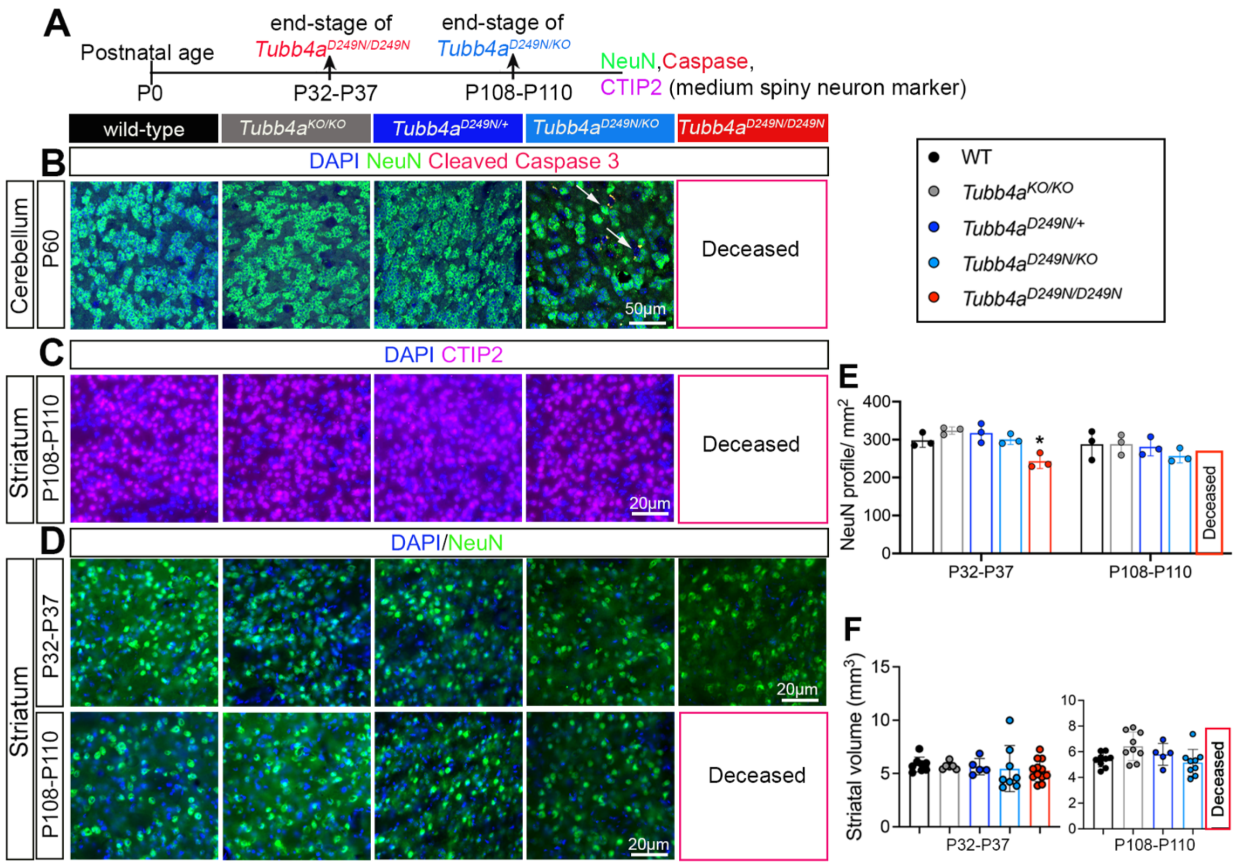

**Supplemental Figure 10** Neuronal (NeuN) profile across all genotypes in cerebellum and striatum. **(A)** Survival timeline of *Tubb4a* mutant mice. (**B**) Representative immunofluorescent images of NeuN+ (green), Caspase 3+ (red), with DAPI (blue, nucleic marker) at P60. Co-localized yellow cells are shown by arrow (NeuN+ and NeuN+ caspase+ double-positive cells quantification is provided in Figure 3). **(C)** Representative immunofluorescent images of CTIP2+ (magenta) and DAPI (blue, nucleic marker) at P108-P110 in the striatum (CTIP2+ quantification of P32-P37 is provided in Figure 3). **(D)** Representative immunofluorescent images of NeuN+ (green) and DAPI (blue, nucleic marker) at P32-P37 and P108-P110 in the striatum. **(E)** Graphical quantification of NeuN profile/mm$^2$ across all genotypes in the striatum**.** **(F)** Graphical quantification of Striatal volume across all genotypes. n=3 mice per genotype and age. Two-way ANOVA was carried out to analyze the statistical significance. Data is presented as Mean (SD). See Supplemental Table 16 for p-values. *p<0.05, **p<0.01 ***p<0.001, ****p<0.0001. Non-significant values are not denoted.

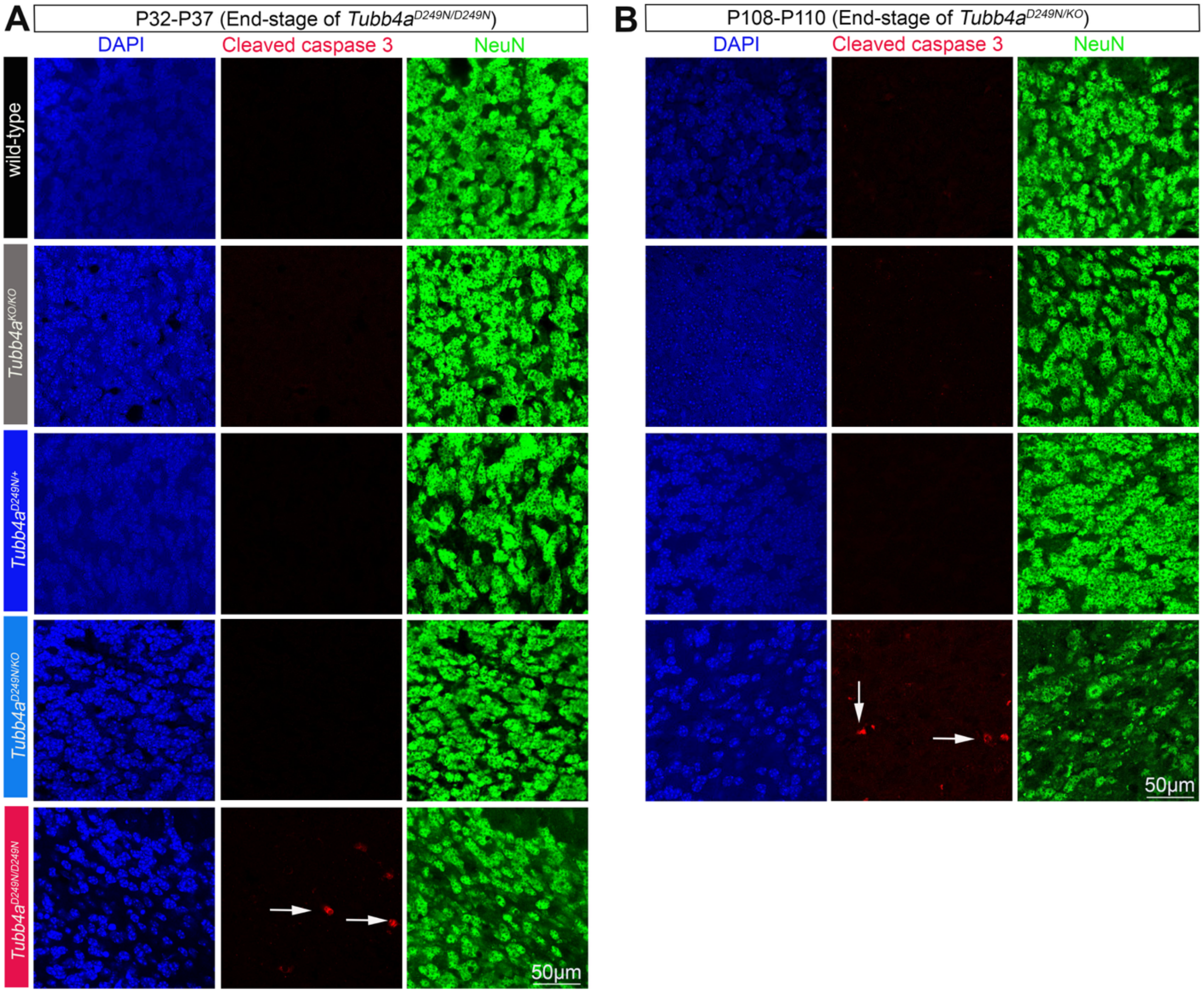

**Supplemental Figure 11** Representative unmerged images of each channel (DAPI (blue), cleaved Caspase 3+ (red), and NeuN+ (green)) of cerebellar sections at **(A)** P32-P37 and **(B)** P108-P110 from Figure 3D. Arrows show the cleaved caspase 3+ cells. The statistical analysis and the merged images are presented in Figure 3.

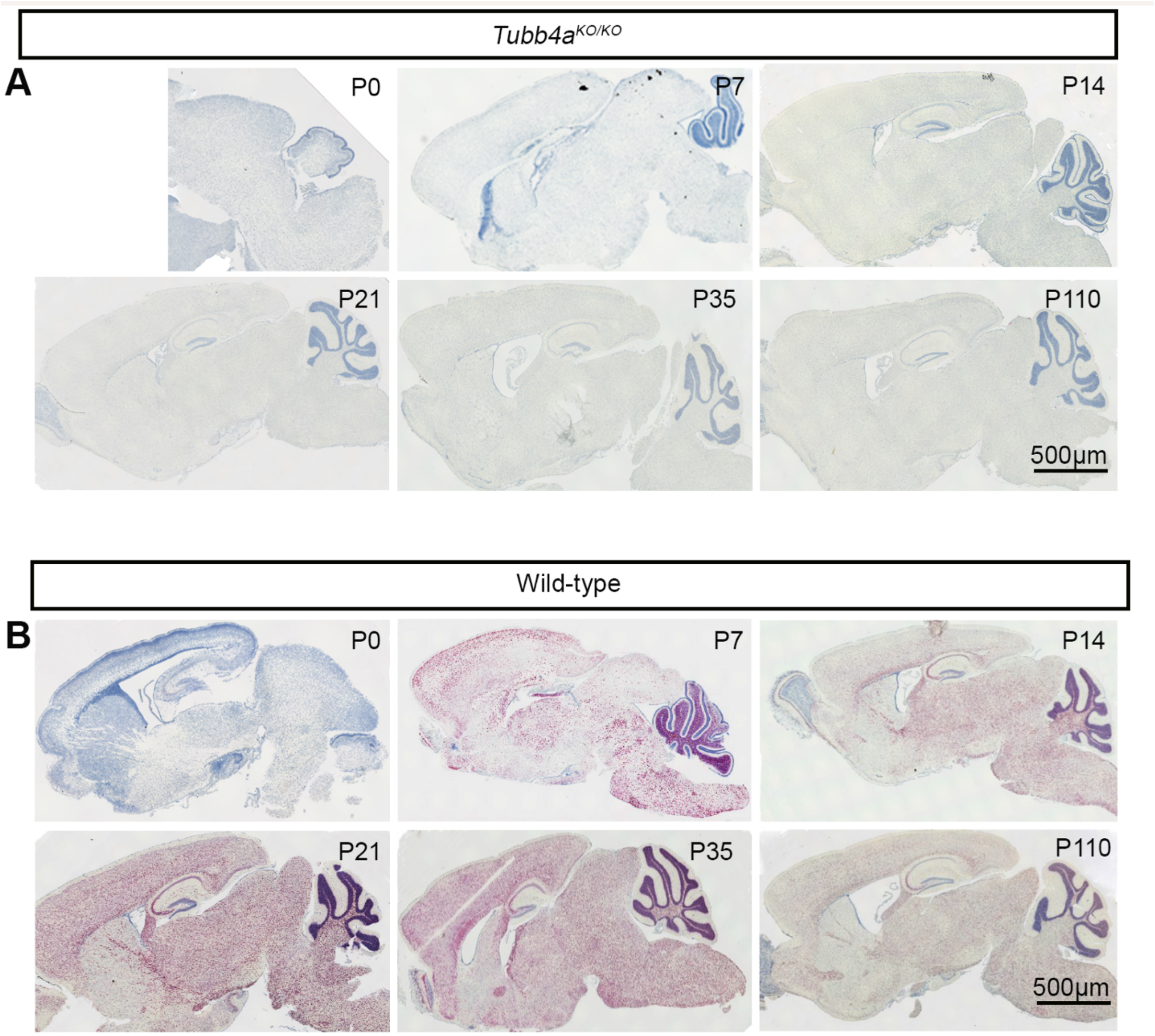

**Supplemental Figure 12** *Tubb4a* tiled RNAscope images **(A-B)** Tubb4a RNAscope tiled sagittal images at different ages in *Tubb4a*$^{KO/KO}$ and WT mice.

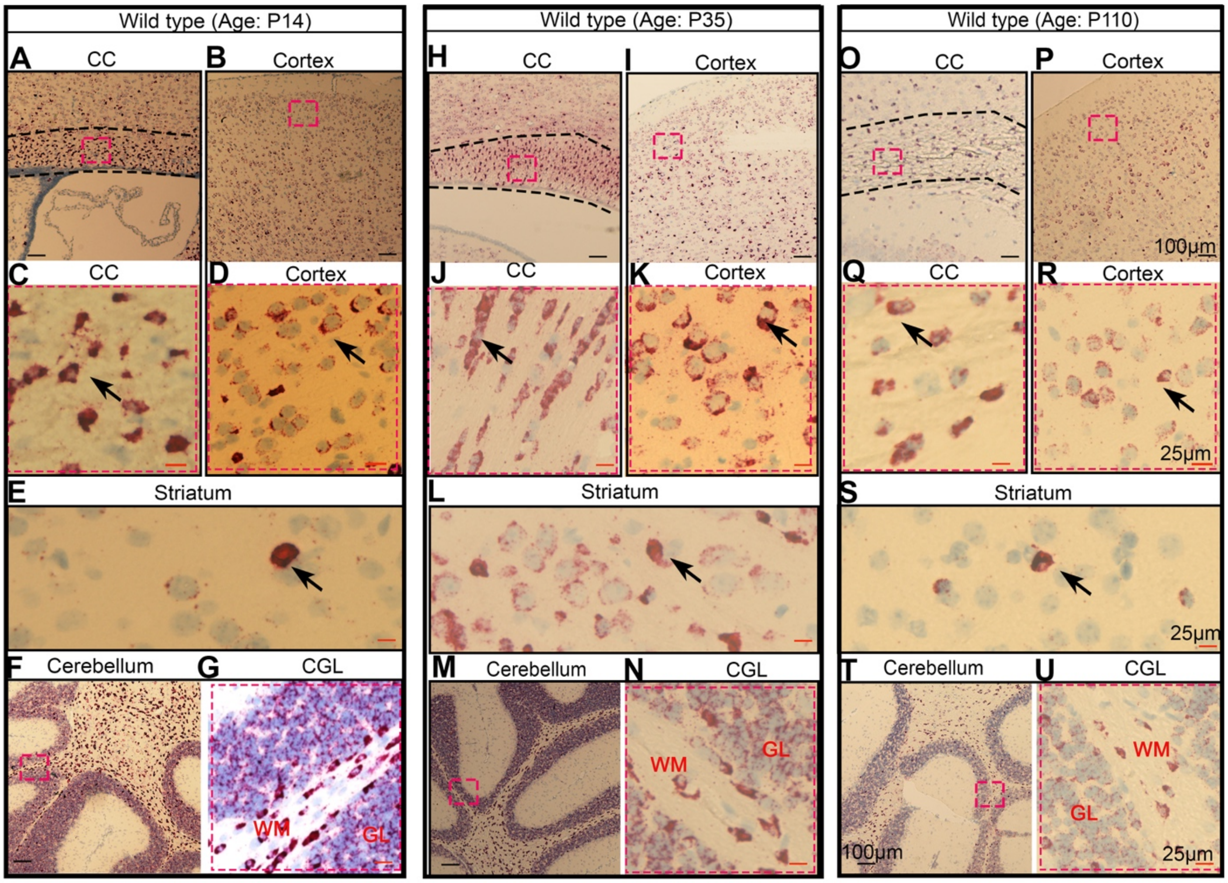

**Supplemental Figure 13** RNAscope in situ hybridization detecting *Tubb4a* transcripts at P14, P35, and P110 in WT sagittal brain sections. (**A-G**) Representative images of low [A-B, and F] and high [C-D, E and G] magnification of Tubb4a RNAscope at P14 of the cortex, corpus callosum (CC), cerebellum, and striatum. **(H-N)** Representative images of low [H-I, M] and high [J-K, L and N] magnification of RNAscope detecting *Tubb4a* transcripts at P35 of the cortex, CC, cerebellum, and striatum. **(O-U)** Representative images of low [O-P, and T] and high [Q-R, S and U] magnification of RNAscope detecting *Tubb4a* transcripts at P110 of the cortex, CC, cerebellum, and striatum. A few *Tubb4a* transcripts are shown by arrows in CC, cortex, and striatum. Tiled images of all corresponding ages are shown in Supplemental Figure 12. Tubb4a expression at P0, P7, and P21 are provided in Figure 4. WM= white matter; GL= granule layer

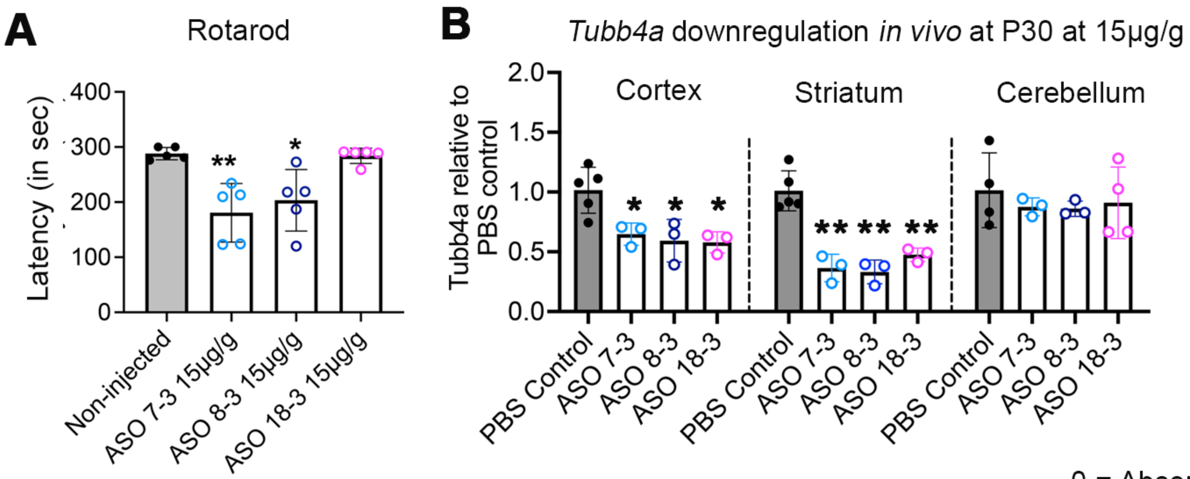

**C** Functional observation battery of ASO treated mice

0 = Absent
1= Present

| AUTONOMIC | mASO#18-3 (15µg/g) | mASO#18-3 (20µg/g) | mASO#18-3 (30µg/g) |
|---|---|---|---|
| Pupil dilator | 0 | 0 | 0 |
| Piloerection | 0 | 0 | 0 |
| Abnormal respiration | 0 | 0 | 0 |

| SENSORIMOTOR | mASO#18-3 (15µg/g) | mASO#18-3 (20µg/g) | mASO#18-3 (30µg/g) |
|---|---|---|---|
| Righting reflex | 0 | 0 | weak |
| Tremor | 0 | 0 | 0 |
| Hindlimb grasp | 0 | 0 | 1 |

| NEUROMUSCULAR | mASO#18-3 (15µg/g) | mASO#18-3 (20µg/g) | mASO#18-3 (30µg/g) |
|---|---|---|---|
| Body tone | 0 | 0 | 1 |
| Gait | 0 | 0 | 1 |
| Dystonia | 0 | 0 | 1 |
| Abormal posture or weight loss | 0 | 0 | 1 |
| Involuntary movements | 0 | 0 | 1 |
| Head bobbing | 0 | 0 | 0 |
| Head flicking | 0 | 0 | 0 |

**Supplemental Figure 14** *In vivo* ASO screening in WT mice. **(A)** Rotarod assay on day 30 (d30) after i.c.v. injection at P1. (**B**) *Tubb4a* downregulation (determined by qRT-PCR) at d30 after i.c.v. injection at P1. *Tubb4a* expression was normalized to the *sfrs9* gene, and the fold change of ASO-injected mice was measured by normalizing to PBS-injected mice. n=3-4 mice per condition. **(C)** Functional observation battery test to assess neurological scores on d30 after i.c.v. injection at P1, and average scores are presented. n=3-4 animals per dose. Two-way ANOVA was carried out to analyze the statistical significance. Data is presented as Mean (SD). *p<0.05, **p<0.001. Non-significant values are not denoted.

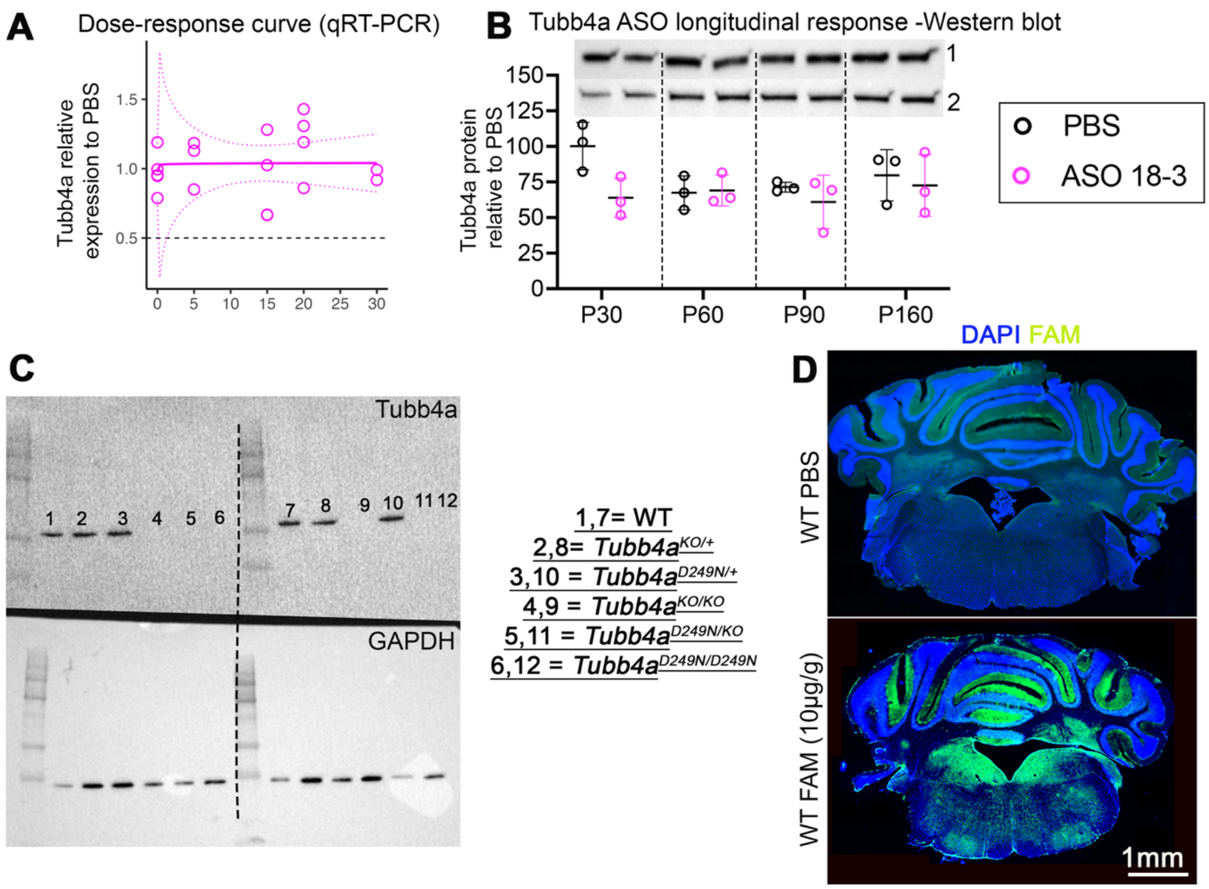

**Supplemental Figure 15** Dose-response curve of the cerebellum and Tubb4a Western blot **(A)** Dose-dependent *Tubb4a* expression of ASO 18-3 ASO relative to PBS-injected controls at P30 in the cerebellum. **(B)** Longitudinal Tubb4a protein levels in WT mice at P30, P60, P90, and P160 in the cerebellum, 1=Tubb4a; 2=vinculin (housekeeping protein). Tubb4a protein levels are normalized to vinculin. n=3-4 mice per condition. **(C)** Tubb4a Western blot image in different genotypes. Non-significant values are not denoted. **(D)** Tiled images of coronal forebrain and cerebellar sections at day 30 after i.c.v injection at P1 with either PBS or FAM tagged Tubb4a ASO 18-3 10µg/g dose.

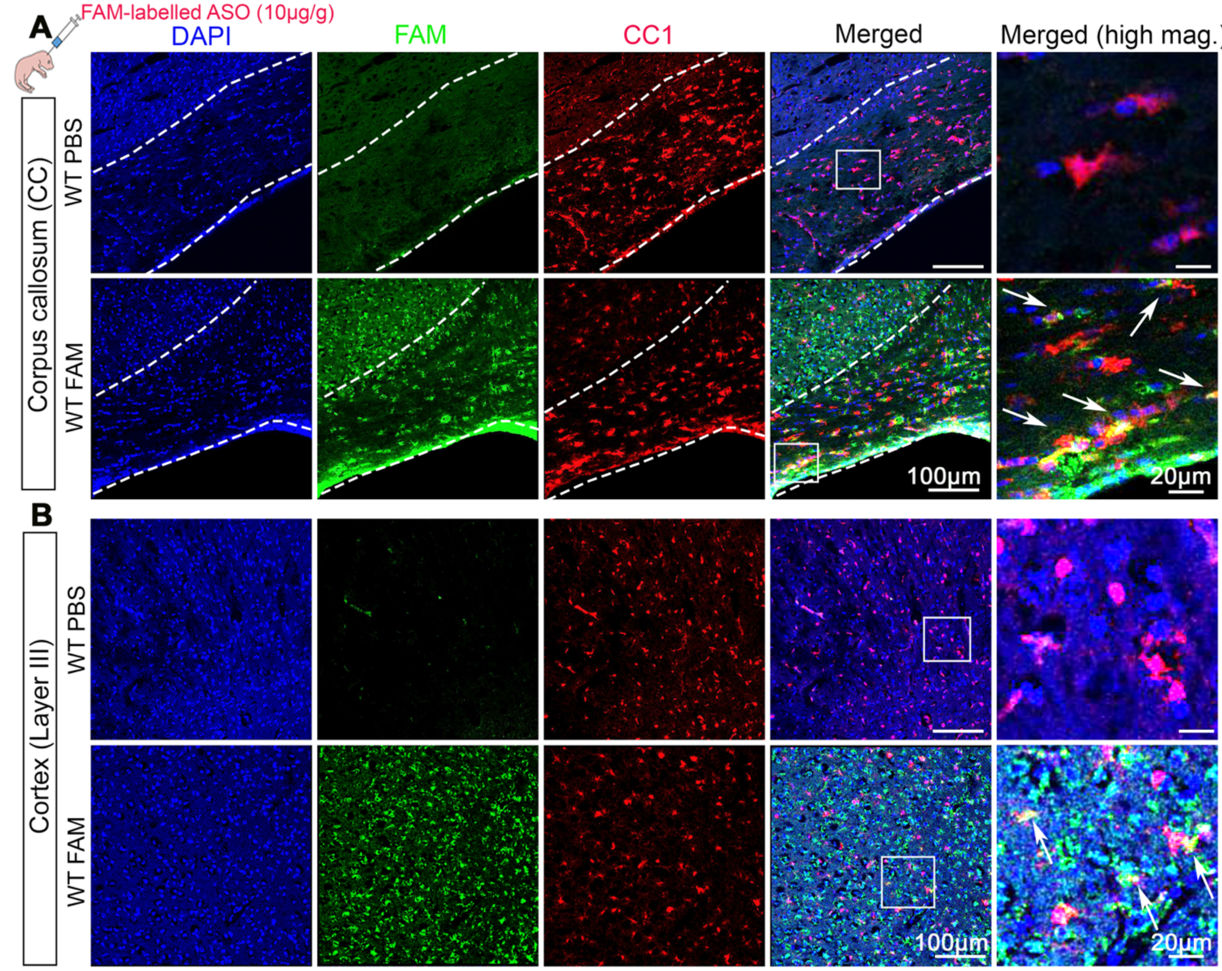

**Supplemental Figure 16** ASO cellular distribution in OLs (CC1 marker) in WT mice in corpus callosum and cortex. **(A)** Representative immunofluorescent images of CC1+ (red), FAM+ (green) and DAPI (blue) staining in the corpus callosum at day 30 after i.c.v. injection at P1 with either PBS or FAM-tagged Tubb4a ASO 18-3 10μg/g dose. These merged images are also provided in Figure 5I in WT FAM row. **(B)** Representative immunofluorescent images of CC1+ (red), FAM+ (green) and DAPI (blue) staining in the cortex at day 30 after i.c.v. injection at P1 with either PBS or FAM-tagged Tubb4a ASO 18-3 10μg/g dose. Arrows show co-localized puncta.

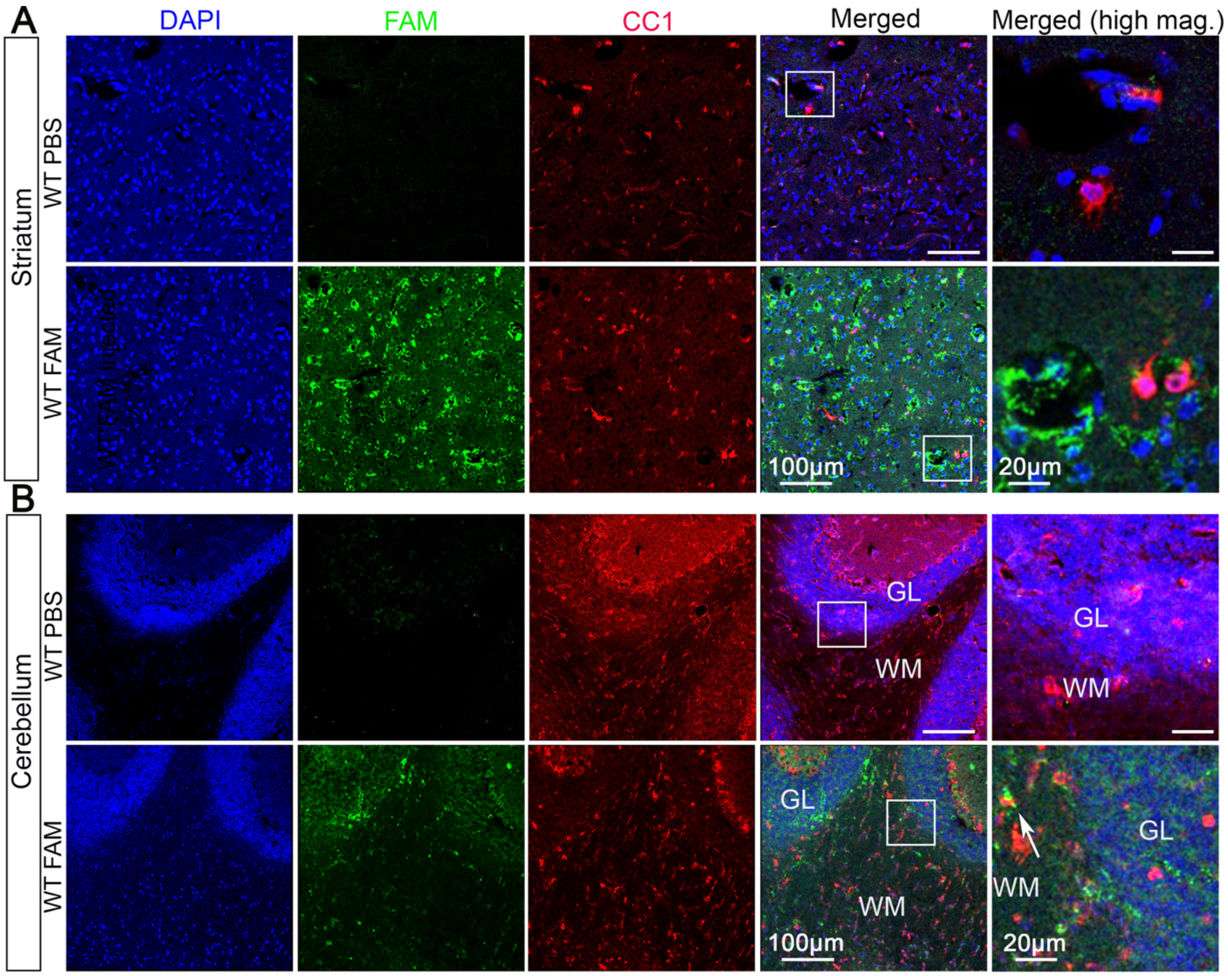

**Supplemental Figure 17** ASO cellular distribution in OLs (CC1 marker) in WT mice in striatum and cerebellum. **(A)** Representative immunofluorescent images of CC1+ (red), FAM+ (green) and DAPI (blue) staining in the striatum at day 30 after i.c.v injection at P1 with either PBS or FAM tagged Tubb4a ASO 18-3 10μg/g dose. **(B)** Representative immunofluorescent images of CC1+ (red), FAM+ (green) and DAPI (blue) staining in the cerebellum at day 30 after i.c.v injection at P1 with either PBS or FAM tagged Tubb4a ASO 18-3 10μg/g dose. Arrows show co-localized puncta.

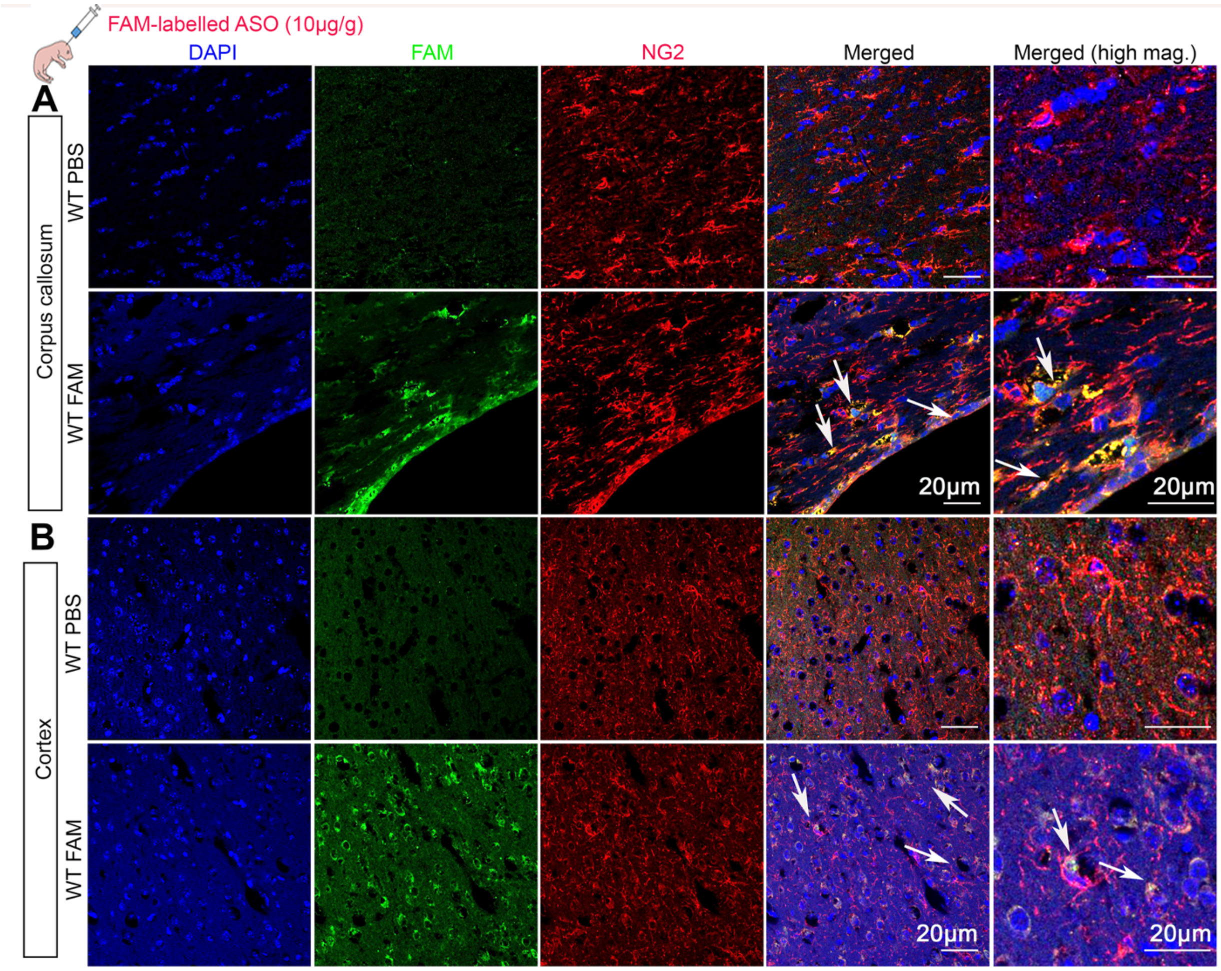

**Supplemental Figure 18** ASO cellular distribution in OPCs (NG2 marker) in WT mice in corpus callosum and cortex. **(A)** Representative immunofluorescent images of NG2+ (red), FAM+ (green) and DAPI (blue) staining in the corpus callosum at day 30 after i.c.v injection at P1 with either PBS or FAM tagged Tubb4a ASO 18-3 10µg/g dose. **(B)** Representative immunofluorescent images of NG2+ (red), FAM+ (green) and DAPI (blue) staining in the cortex at day 30 after i.c.v injection at P1 with either PBS or FAM tagged Tubb4a ASO 18-3 10µg/g dose. Arrows show FAM and NG2 co-localized cells.

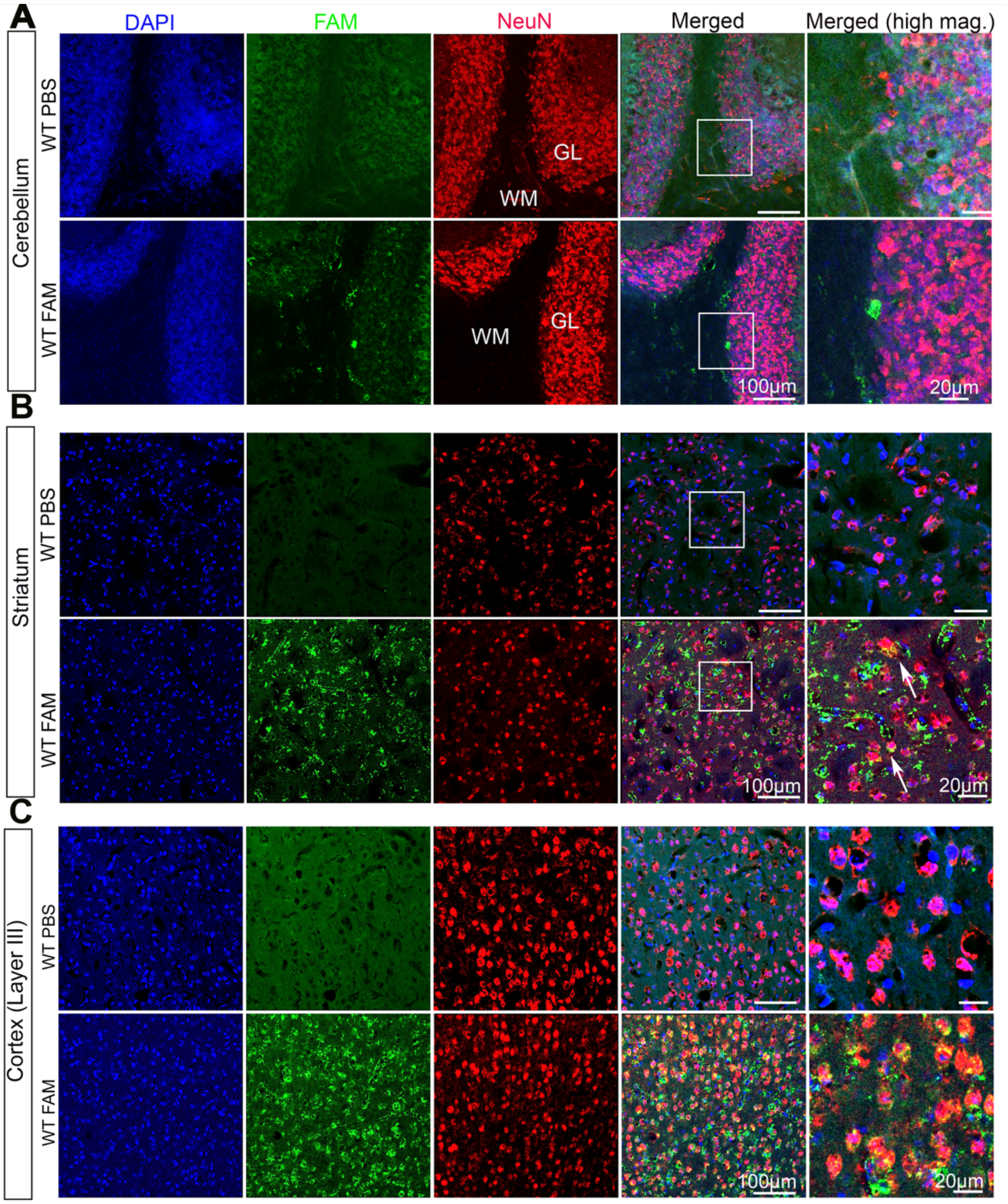

**Supplemental Figure 19** ASO cellular distribution in neurons (NeuN marker) in WT mice in cerebellum, striatum and cortex. **(A)** Representative immunofluorescent images of NeuN+ (red), FAM+(green) and DAPI (blue) staining in the cerebellum at day 30 after i.c.v injection at P1 with either PBS or FAM tagged Tubb4a ASO 18-3 10μg/g dose; **(B)** Representative immunofluorescent images of NeuN+ (red), FAM+(green) and DAPI (blue) staining in the striatum at day 30 after i.c.v injection at P1 with either PBS or FAM tagged Tubb4a ASO 18-3 10μg/g dose. **(C)** Representative immunofluorescent images NeuN+ (red), FAM+ (green) and DAPI (blue) staining in the cortex at day 30 after i.c.v injection at P1 with either PBS or FAM tagged Tubb4a ASO 18-3 10μg/g dose. Arrows show co-localized puncta.

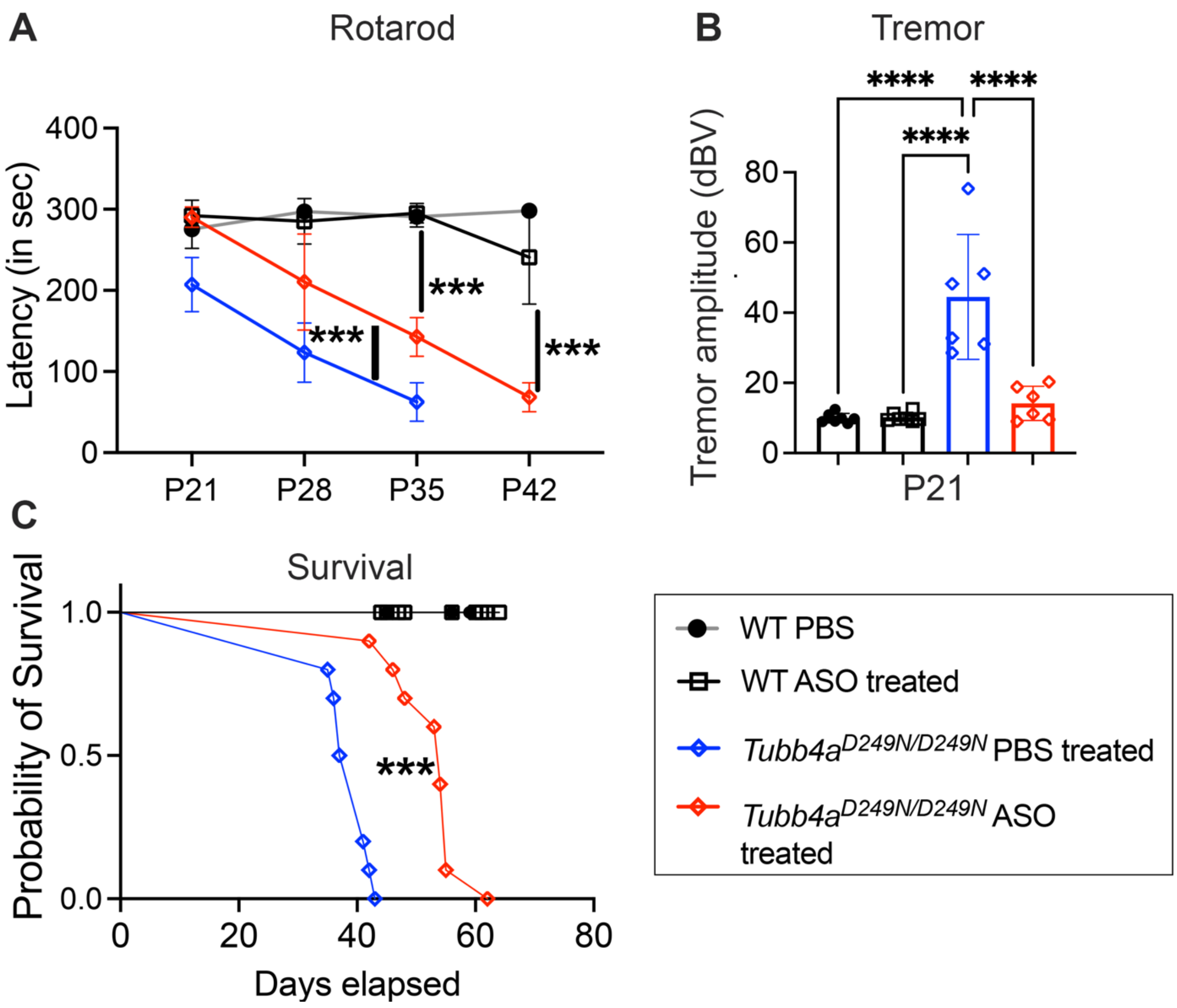

**Supplemental Figure 20** Survival and behavior in ASO-treated groups. Treatment groups include WT PBS, WT ASO, *Tubb4a*$^{D249N/D249N}$ PBS-treated, and *Tubb4a*$^{D249N/D249N}$ ASO-treated. **(A)** Graphical presentation of accelerating rotarod test of treatment groups. n=5-7 mice per treatment group. P-values are calculated using repeated measures ANOVA with mixed effects analysis. **(B)** Graphical presentation of tremor amplitudes of treatment groups. n=5-7 mice per treatment group. P-values are calculated using one-way ANOVA. **(C)** Kaplan-Meier survival analysis of treatment groups. n=8-10 mice per group. Data presented as Mean (SD). P-values calculated using log-rank test. *p<0.05, **p<0.01, ***p<0.001, ****P<0.0001.

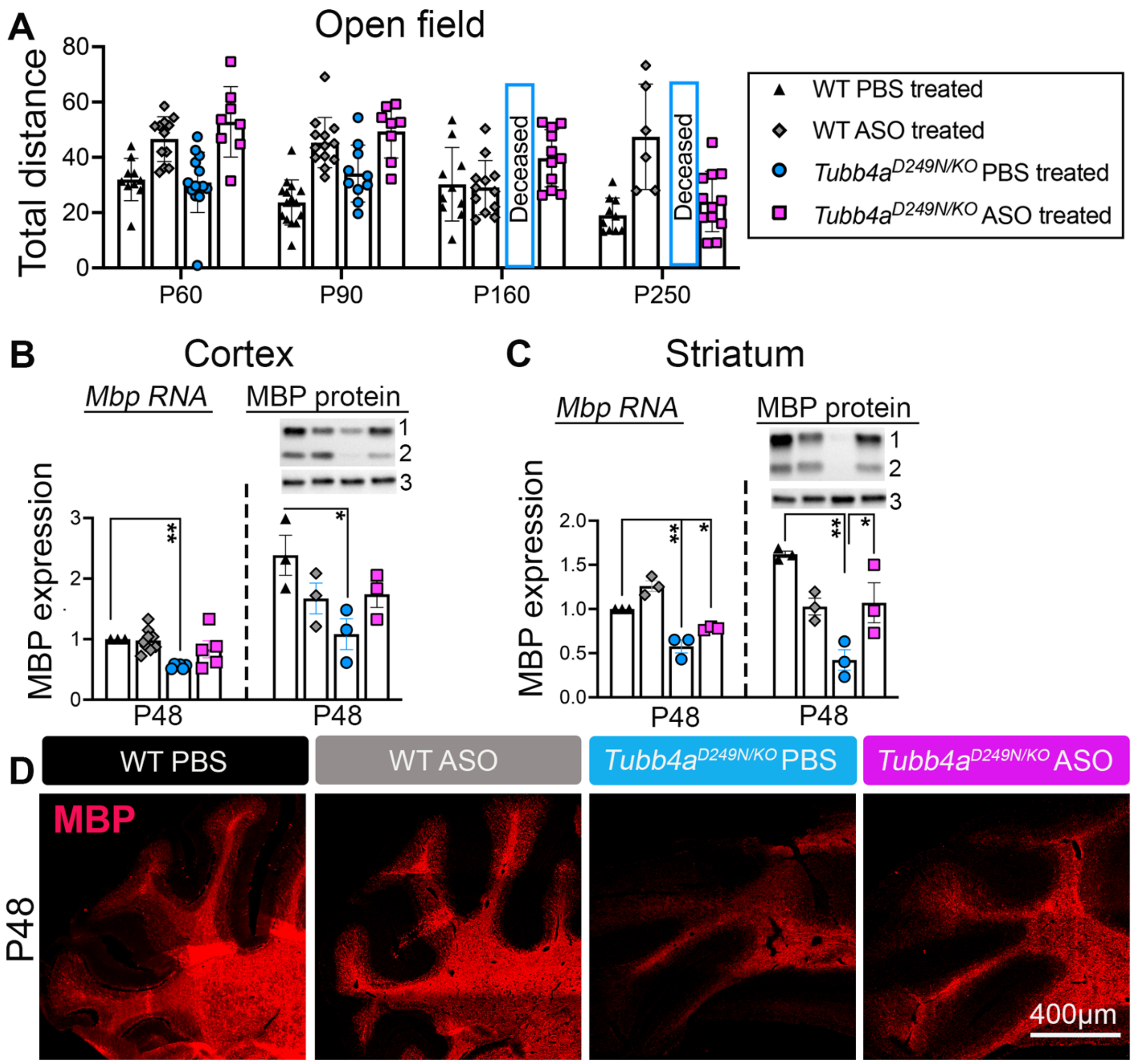

**Supplemental Figure 21** Open field test and MBP immunostaining across ASO treatment groups (A) Total distance traveled in open field test. n=10-15 mice per treatment group and age. (B) Graphical presentation of quantification of *Mbp* transcripts (using Nanostring) and MBP western blot in the cortex (See Nanostring in Supplemental Figure 22). (C) Graphical presentation of Mbp transcripts (via Nanostring) and MBP protein in the striatum. n=3-8 mice for Nanostring and MBP western blotting. GAPDH is used as a housekeeping protein to normalize the expression of MBP protein. 1= MBP ~25kDa; 2= MBP ~18.5kDa; and 3= GAPDH. Data is presented as Mean (SD). *p<0.05, **p<0.01, ***P<0.001, ****p<0.0001. (D) Representative immunofluorescent images of MBP (red) in the cerebellum at P48.

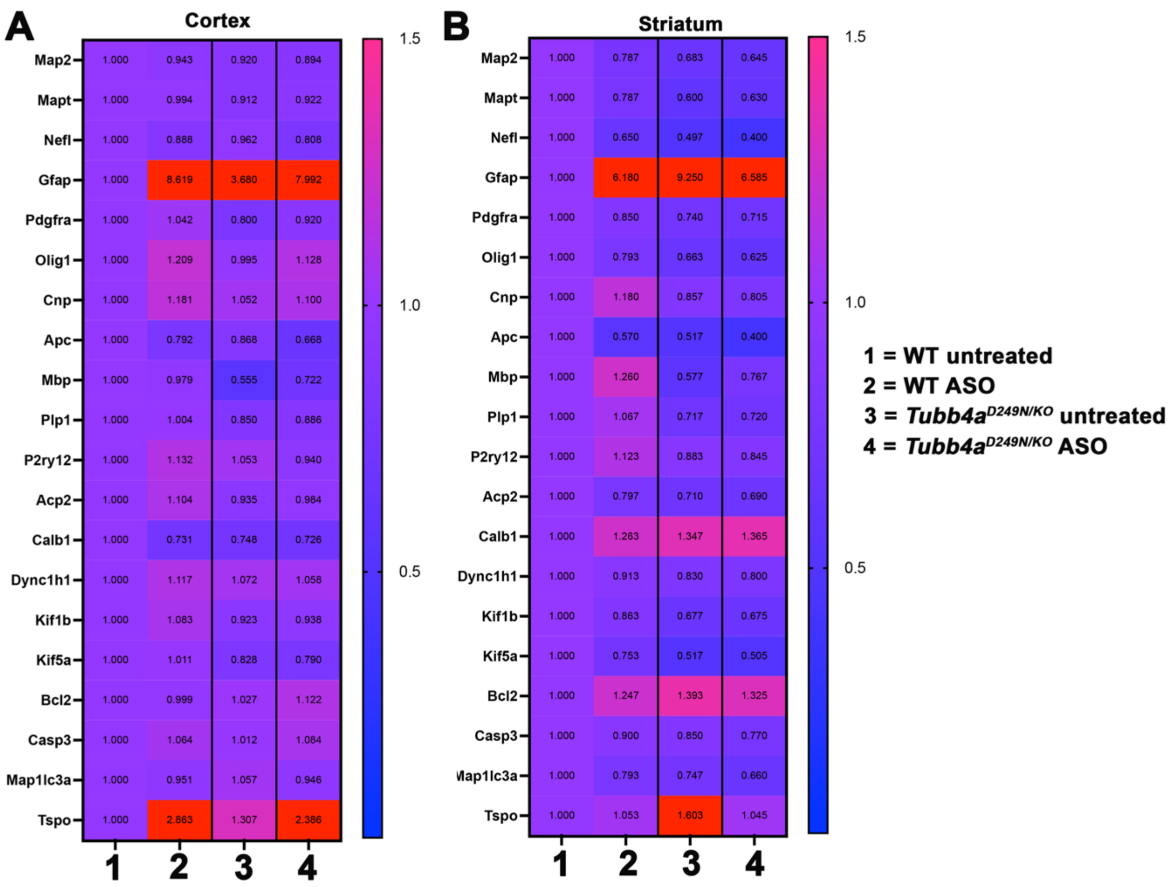

**Supplemental Figure 22** Nanostring panel

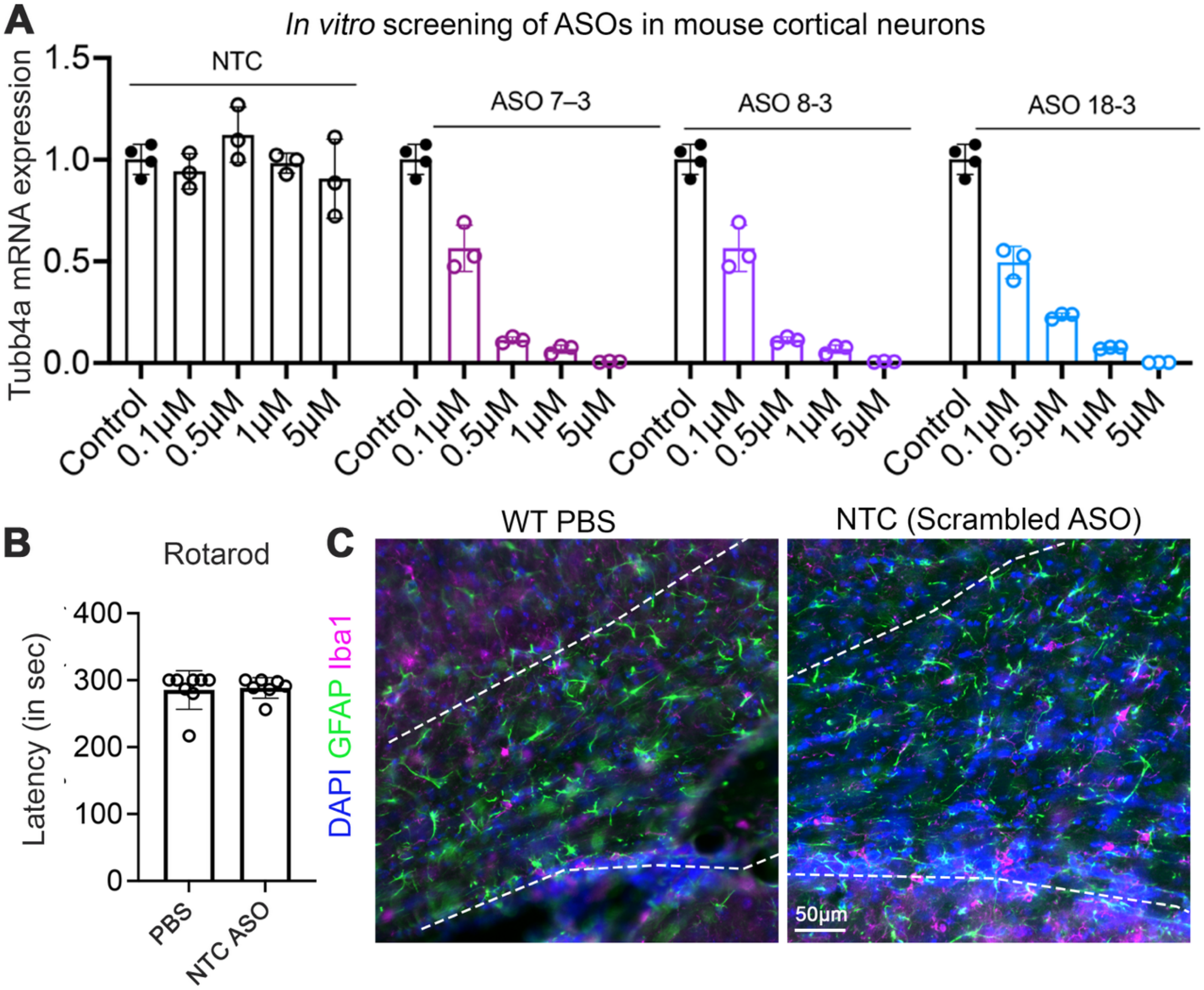

**Supplemental Figure 23 (A)** *In vitro* ASO screening with selected Tubb4a ASOs and scrambled ASO (NTC ASO) to assess *Tubb4a* downregulation. **(B)** Graphical presentation of accelerated Rotarod at P30 of WT PBS injected and scrambled (NTC) ASO injected mice (n=6-7 mice per group). A t-test was used to compare the two groups. Data is presented as Mean (SD). **(C)** GFAP (green), Iba1 (magenta), and DAPI (blue) immunostaining on brain sections at P30 (dotted lines show corpus callosum) injected with WT PBS and NTC injected mice.

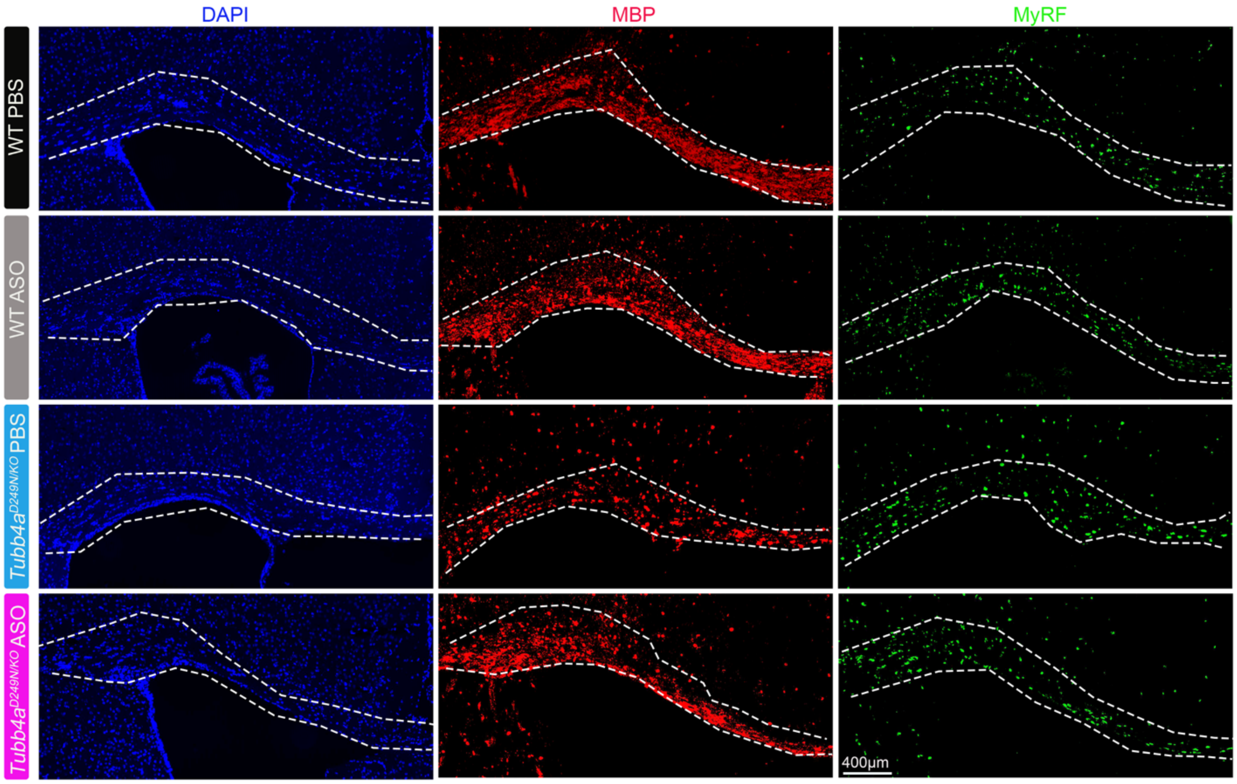

**Supplemental Figure 24** Stitched RNA scope images of *Mbp* and *Myrf* mRNA highlighting corpus callosum sections at P48. The merged pictures for this staining are provided in Figure 7A (A') column.

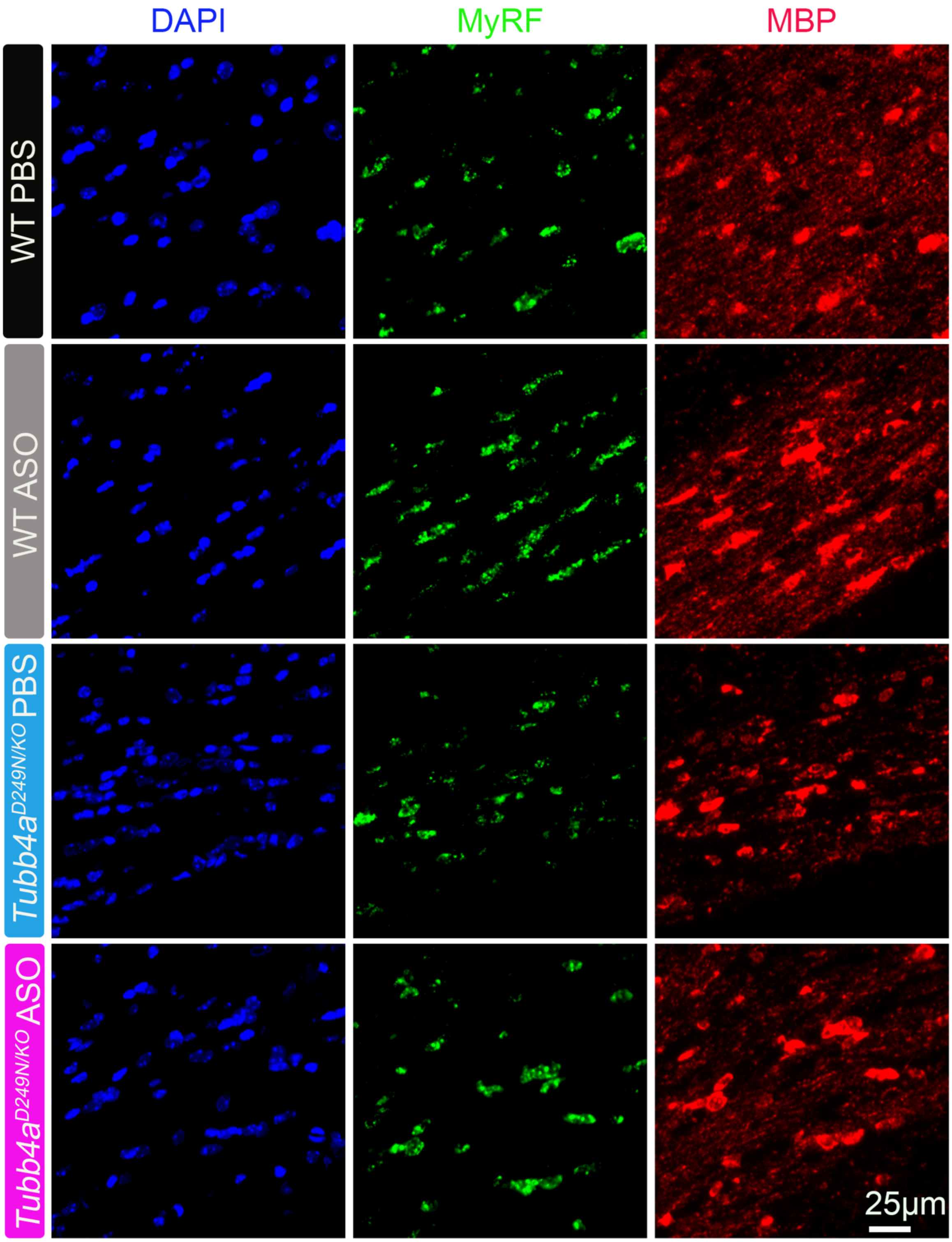

**Supplemental Figure 25** MBP (red) and Myrf (green) RNA scope unmerged images of corpus callosum sections across treatment groups from Figure 7 column A”. The graphical analysis and merged images are presented in Figure 7.

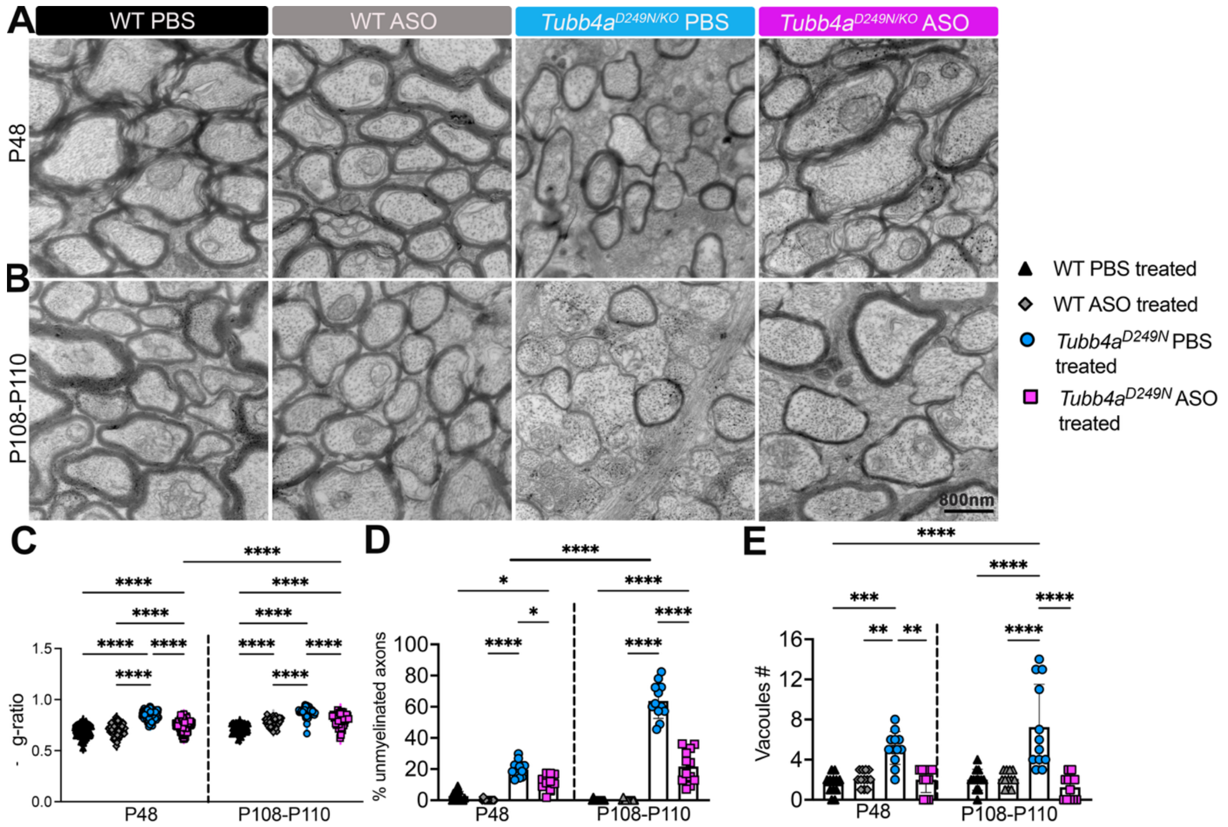

**Supplemental Figure 26** Optic nerve of ASO-treated *Tubb4a* mutant relative to controls. **(A-B)** Representative electron micrographs of the optic nerve in treatment groups at P48 and P108-P110. **(C)** g-ratio in treated groups at P48 and P108-P110 **(D)** % unmyelinated axons in treated groups. **(E)** Number of vacuoles (#) in treated groups at P48 and P108-P110. For EM data, n=3-4 animals are used, and at least 4-5 images per animal are used. For the g-ratio, at least 30-40 axons per image were used to measure the g-ratio. P-values are calculated using one-way or two-way ANOVA with Tukey corrections. Data is presented as Mean (SD). *p<0.05, **p<0.01 ***p<0.001, ****p<0.0001.

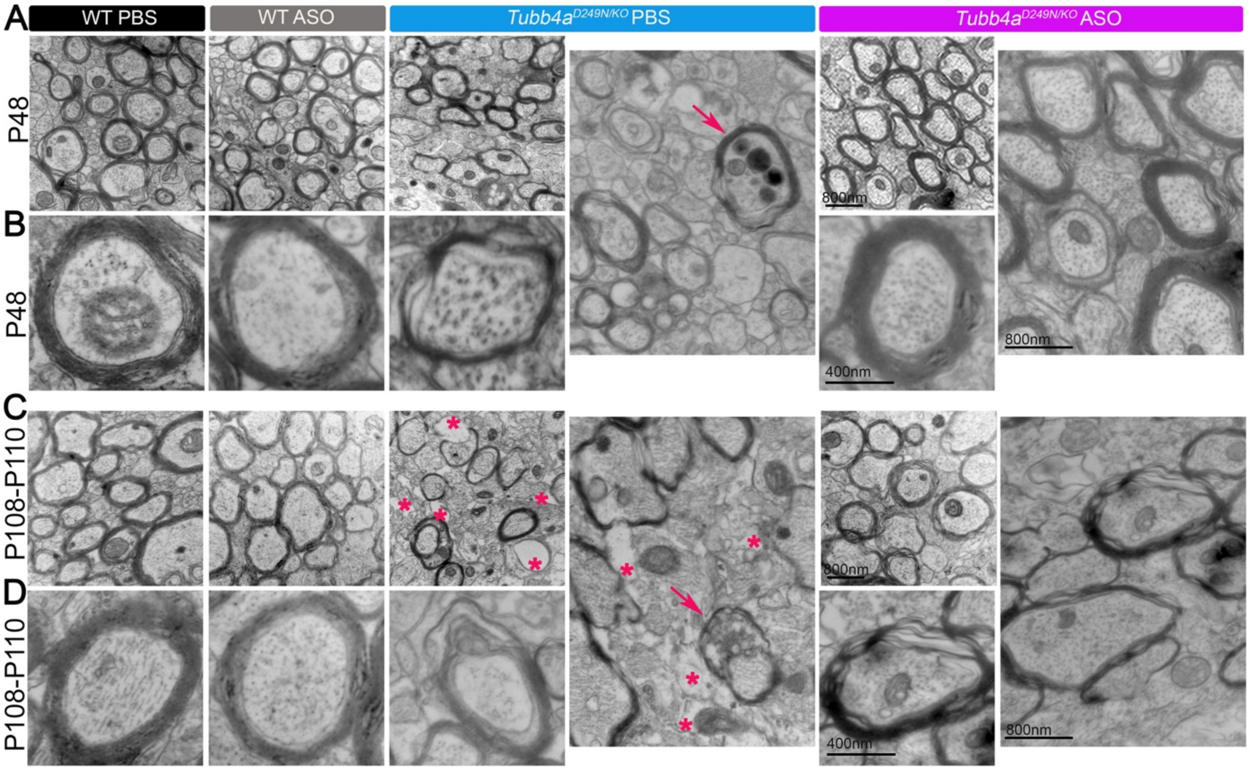

**Supplemental Figure 27** Corpus callosum of ASO-treated *Tubb4a* mutant relative to controls. This figure shows the abnormal vacuoles in untreated *Tubb4a*$^{D249N/KO}$ mice, and with ASO treatment, the number of vacuoles is reduced. Please see the statistical analysis in Figure 7E. **(A-B)** Representative electron micrographs at the low and higher magnification of the corpus callosum in treatment groups at P48. The *Tubb4a*$^{D249N/KO}$ *PBS* injected image is also presented in Figure 7 B. **(C-D)** Representative electron micrographs at the low and higher magnification of the corpus callosum in treatment groups at P108-P110. * denotes myelin vacuolization.

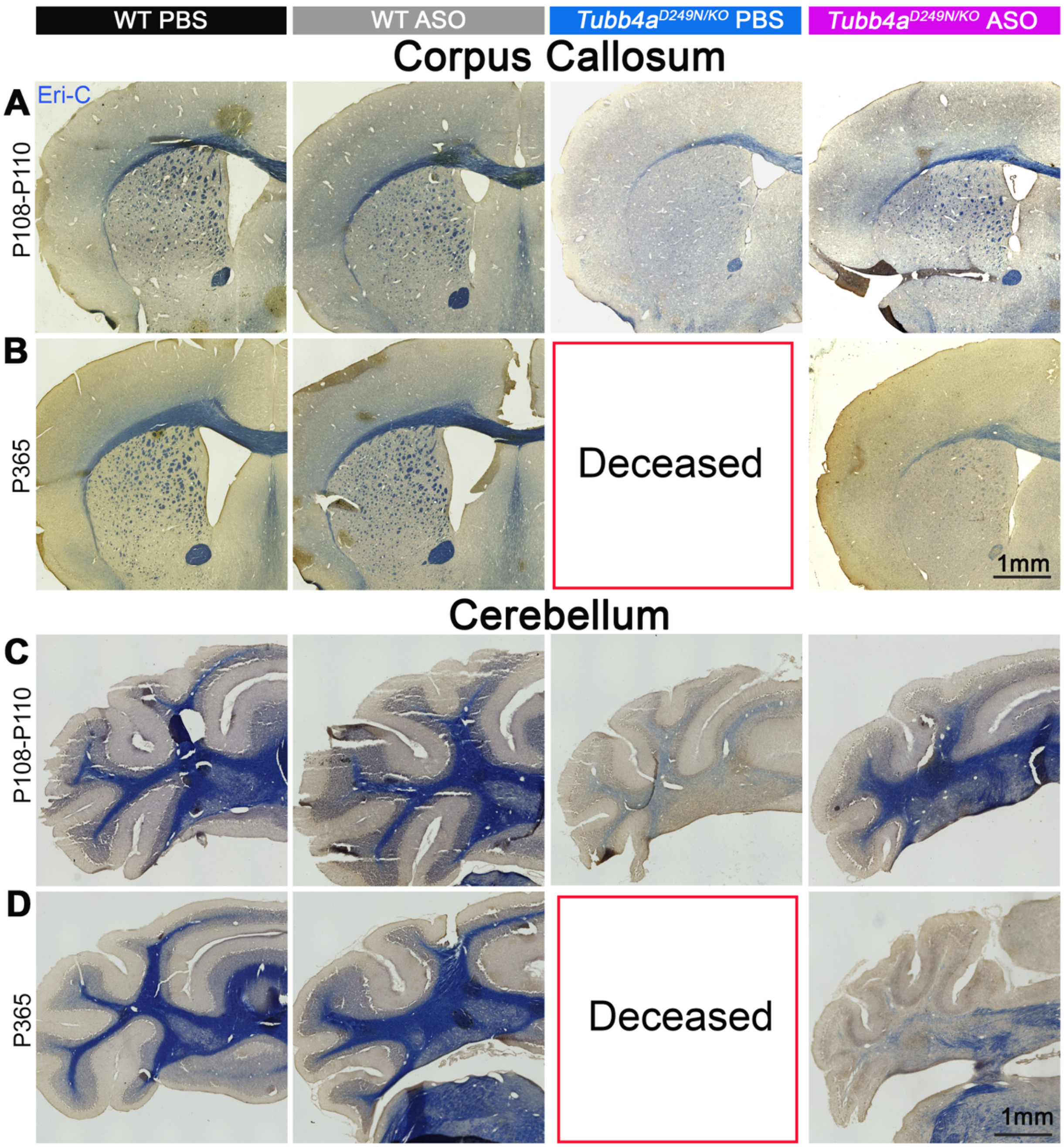

**Supplemental Figure 28** Eri-C staining in corpus callosum and cerebellum across different treatment groups. **(A-B)** Representative tiled histochemical images of Eri-C images of the forebrain at P108-P110 (A), and P365 (B). **(C-D)** Representative tiled histochemical images of Eri-C of the cerebellum at P108-P110 (A), and P365 (B).

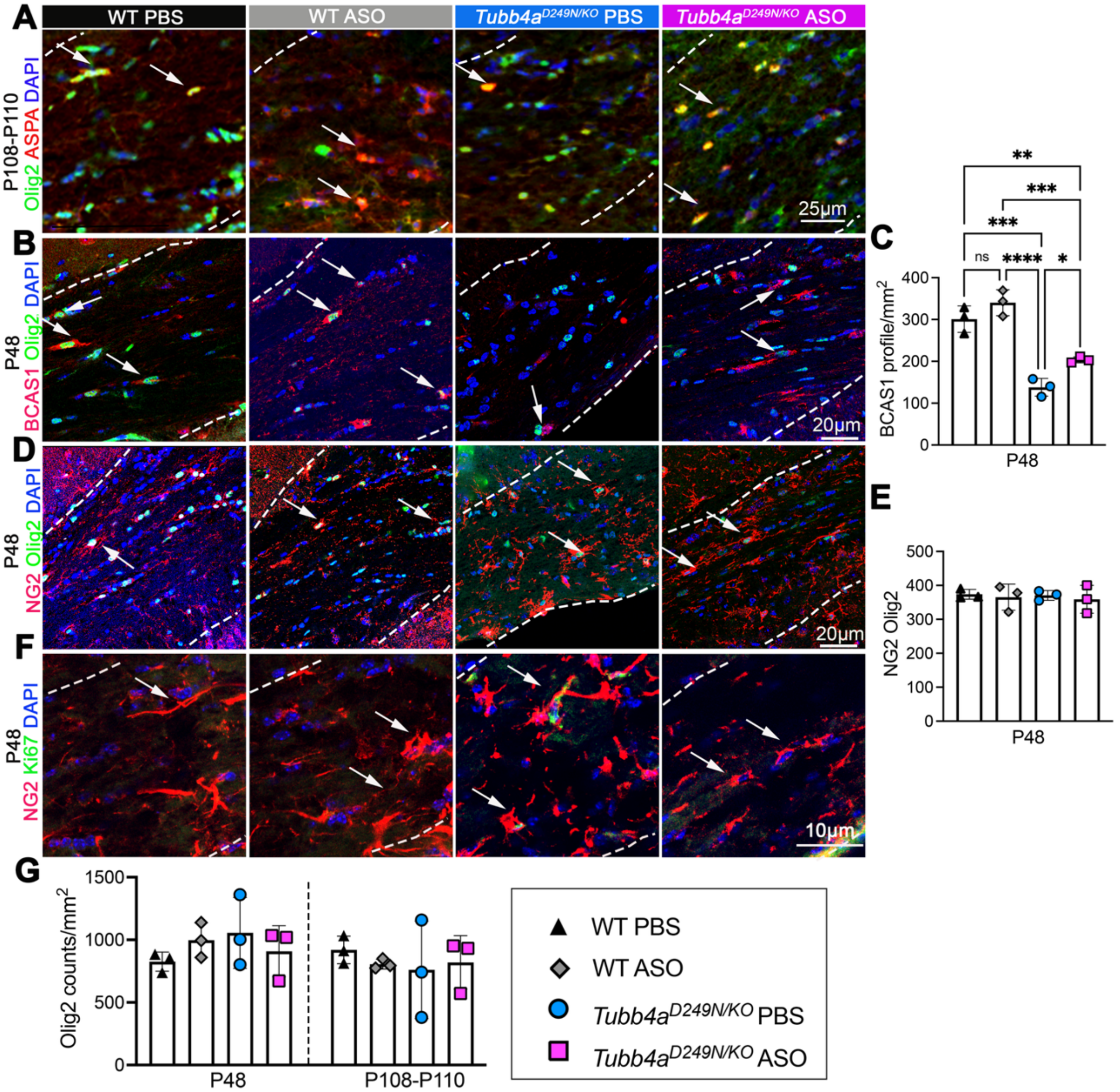

**Supplemental Figure 29** OL lineage cell staining in treatment groups in the corpus callosum. **(A)** Representative images of ASPA+ (red), Olig2 (green), and DAPI (blue) staining at P108-P110. **(B)** Representative images of BCAS1+ (red), Olig2+ (green) and DAPI (blue) staining at P48. **(C)** Graphical representation of BCAS1+Olig2+ counts/mm$^2$ at P48. **(D)** Representative images of NG2+ (red), Olig2+ (green) and DAPI (blue) staining at P48. **(E)** Graphical representation of NG2+Olig2+ counts/mm$^2$ at P48. n=3 mice per treatment group. One-way ANOVA was carried out to analyze the statistical significance. **(F)** Representative images of NG2+ (red), Ki67+ (green) and DAPI (blue) staining at P48. **(G)** Graph of Olig2 counts/mm$^2$ at P48 and P108-P110. n=3 mice per treatment group and age. Two-way ANOVA was carried out to analyze the statistical significance. Data is presented as Mean (SD). *p<0.05, **p<0.01 ***p<0.001, ****p<0.0001. Non-significant values are not denoted. Arrows represent the respective cell types.

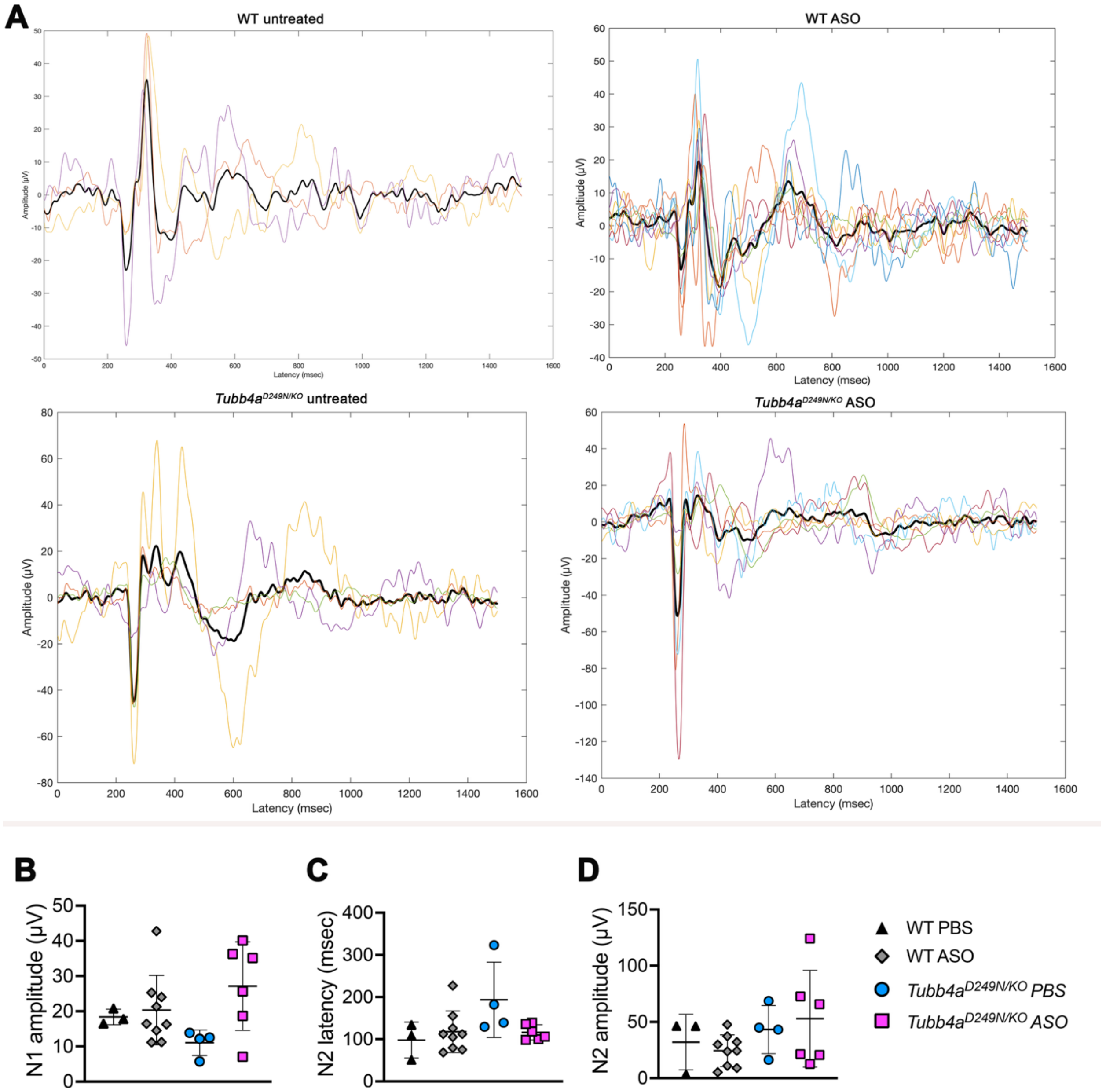

**Supplemental Figure 30** Evoked potential peaks and measurements of evoked potential latencies and amplitude of different treatment groups. **(A)** Evoked potential peaks of all the treatment groups. Graphical presentation of **(B)** N1 amplitude **(C)** N2 latencies **(D)** N2 amplitude. n=3-9 mice per treatment group. One-way ANOVA was carried out to analyze the statistical significance. Data is presented as Mean (SD).

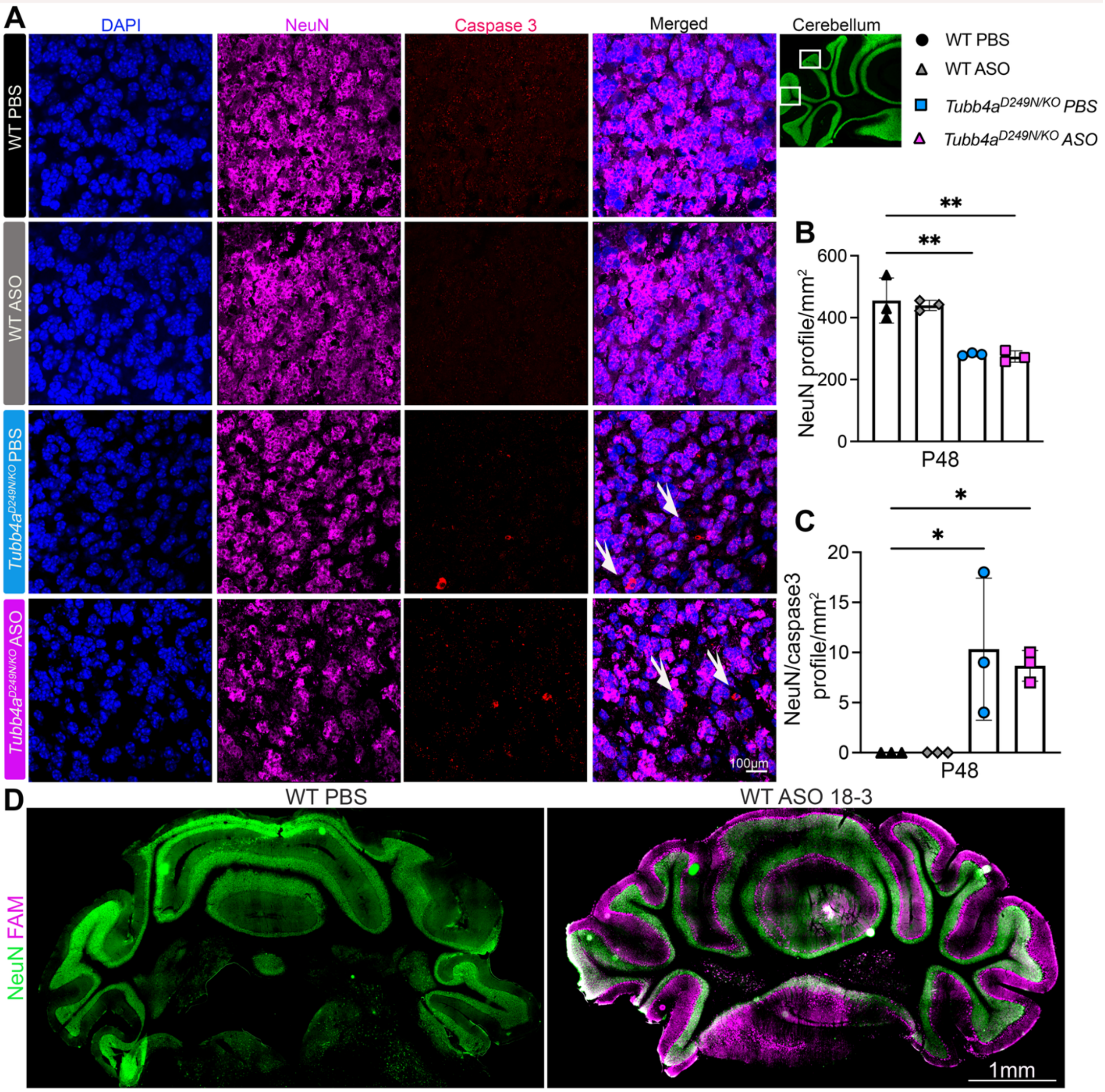

**Supplemental Figure 31** ASO treatment and cerebellar granule neuron outcomes. **(A)** Representative immunofluorescent images of NeuN+ (magenta), Caspase+ (red) and DAPI (blue) staining at P48. Arrows show co-localized NeuN+ and Caspase+ cells. **(B)** Graphical representation of NeuN+ DAPI+ profile/mm$^2$ at P48. **(C)** Graphical representation of NeuN+ Caspase+ DAPI+ profile/mm$^2$ at P48. n=3 mice per treatment group and age. One-way ANOVA was carried out to analyze the statistical significance. Data is presented as Mean (SD). *$p<0.05$, **$p<0.001$. **(D)** Representative stitched images of cerebellum with NeuN+ (green) and FAM+ (magenta) on day 30 after injection of FAM 18-3 ASO via cisterna magna at P1-2.

| Supplemental Table 1: ASO sequences | | | | | |
|---|---|---|---|---|---|
| **ASO#** | **ASO Gen#** | **ASO sequences** | **Gapmer** | **Total Length** | **Total PS bonds** |
| **ASO 1316** | IDT ASOs | @a*@c*@a*+t*a*c*g*g*c*t*g*t*c* @t*@t*@g* | 3-10-3 | 16 | 16 |
| **ASO 1** | Gen1 | +G*+C*+U*+U*g*c*a*g*g*t*g*c*a* +C*+G*+A*+U* | 3-10-3 | 17 | 17 |
| **ASO 2** | Gen1 | +U*+G*+U*+C*g*a*t*g*c*a*g*t*a* +G*+G*+U*+C* | 3-10-3 | 17 | 17 |
| **ASO 3** | Gen1 | +U*+G*+U*c*g*a*t*g*c*a*g*t*a*+ G*+G*+U* | 3-10-3 | 16 | 16 |
| **ASO 4** | Gen1 | +C*+C*+U*+C*g*t*t*g*t*c*g*a*t*+ G*+C*+A*+G* | 3-10-3 | 17 | 17 |
| **ASO 5** | Gen1 | +U*+U*+G*+A*g*g*t*c*c*c*c*g*t*+ A*+G*+G*+U* | 3-10-3 | 17 | 17 |
| **ASO 6** | Gen1 | +G*+C*+U*+C*g*t*c*t*a*c*c*t*c*+ C*+U*+U*+C* | 3-10-3 | 17 | 17 |
| **ASO 7** | Gen1 | +G*+C*+U*c*g*t*c*t*a*c*c*t*c*+C *+U*+U* | 3-10-3 | 16 | 16 |
| **ASO 8** | Gen1 | +U*+G*+C*t*c*g*t*c*t*a*c*c*t*+C* +C*+U* | 3-10-3 | 16 | 16 |
| **ASO 9** | Gen1 | +U*+C*+U*+C*g*t*c*c*a*t*g*c*c*+ U*+U*+C*+G* | 3-10-3 | 17 | 17 |
| **ASO 10** | Gen1 | +C*+C*+A*+U*c*t*c*g*t*c*c*a*t*+ G*+C*+C*+U* | 3-10-3 | 17 | 17 |
| **ASO 11** | Gen1 | +U*+C*+U*+U*c*g*a*a*c*t*c*g*c* +C*+C*+U*+C* | 3-10-3 | 17 | 17 |
| **ASO 12** | Gen1 | +C*+C*+U*c*c*t*c*t*t*c*g*a*a*+C *+U*+C* | 3-10-3 | 16 | 16 |
| **ASO 13** | Gen1 | +G*+G*+U*c*a*g*a*g*g*t*a*a*+G *+G*+C* | 3-10-3 | 15 | 15 |
| **ASO 14** | Gen1 | +U*+G*+C*+A*c*g*a*t*t*t*c*c*c*+ G*+C*+A*+U* | 3-10-3 | 17 | 17 |

| | | | | | |
|---|---|---|---|---|---|
| **ASO 15** | Gen1 | +G*+U*+U*+G*t*c*g*a*t*g*c*a*g* +U*+A*+G*+G* | 3-10-3 | 17 | 17 |
| **ASO 16** | Gen1 | +C*+C*+U*+C*+G*t*t*g*t*c*g*a*t* g*c*+A*+G*+U*+A* | 3-10-3 | 19 | 19 |
| **ASO 17** | Gen1 | +C*+U*+C*+G*t*t*g*t*c*g*a*t*g*+ c*+A*+G*+U* | 3-10-3 | 17 | 17 |
| **ASO 18** | Gen1 | +U*+G*+C*t*c*g*t*c*t*a*c*c*+U*+ C*+C* | 3-10-3 | 15 | 15 |
| **ASO 19** | Gen1 | +G*+U*+G*+C*t*g*t*t*g*c*c*g*a*t *+G*+A*+A*+G* | 3-10-3 | 18 | 18 |
| **ASO 20** | Gen1 | +G*+U*+G*c*t*g*t*t*g*c*c*g*a*+U *+G*+A* | 3-10-3 | 16 | 16 |
| **ASO 21** | Gen1 | +A*+U*+C*+U*+C*g*t*c*c*a*t*g*c *c*t*+U*+C*+G*+C* | 3-10-3 | 19 | 19 |
| **ASO 22** | Gen1 | +C*+C*+A*+U*c*t*c*g*t*c*c*a*t*g *+C*+C*+U*+U* | 3-10-3 | 18 | 18 |
| **ASO 23** | Gen1 | +U*+C*+C*+A*t*c*t*c*g*t*c*c*a*+ U*+G*+C*+C* | 3-10-3 | 17 | 17 |
| **ASO 24** | Gen1 | +U*+U*+C*g*a*a*c*t*c*g*c*c*c*+ U*+C*+U* | 3-10-3 | 16 | 16 |
| **ASO 25** | Gen1 | +C*+U*+C*+U*t*c*g*a*a*c*t*c*g* +C*+C*+C*+U* | 3-10-3 | 17 | 17 |
| **ASO 4A** | Gen 2 | +C*+C+U+cg*t*t*g*t*$^{m}$c*g*a*t*+G +C+A+G | 3-10-3 | 17 | 10 |
| **ASO 4B** | Gen 2 | +C*+C+U+cg*t*t*g*t*(5’me)c*g*a* t*+G+C+A*+G | 3-10-3 | 17 | 11 |
| **ASO 6A** | Gen 2 | +G*+C+U+cg*t*$^{m}$c*t*a*$^{m}$c*$^{m}$c*t*c* +C+U+U+C | 3-10-3 | 17 | 10 |
| **ASO 6B** | Gen 2 | +G*+C+U+Cg*t*$^{m}$c*t*a*$^{m}$c*$^{m}$c*t* c*+C+U+U* +C | 3-10-3 | 17 | 11 |
| **ASO 7A** | Gen 2 | +G*+C+U+cg*t*$^{m}$c*t*a*$^{m}$c*$^{m}$c*t* $^{m}$c+C+U+U | 3-10-3 | 16 | 10 |

| | | | | | |
|---|---|---|---|---|---|
| **ASO 7B** | Gen 2 | +G*+C+U+cg*t*$^{m}$c*t*a*$^{m}$c*$^{m}$c*t*$^{m}$c+C+U* +U | 3-10-3 | 16 | 11 |
| **ASO 6-1** | Gen 3 | +G*+C$^{m}$*+U*+C$^{m}$*g*t*c$^{m}$*t*a*c$^{m}$*c$^{m}$*t*c$^{m}$*+C$^{m}$*+U*+U*+C$^{m}$ | 4-9-4 | 16 | 16 |
| **ASO 6-2** | Gen 3 | +G*+C$^{m}$+U+C$^{m}$*g*t*c$^{m}$*t*a*c$^{m}$*c$^{m}$*t*c$^{m}$*+C$^{m}$+U+U*+C$^{m}$ | 4-9-4 | 12 | 12 |
| **ASO 6-3** | Gen 3 | +U*+G*+C$^{m}$*+U*+C$^{m}$*g*t*c$^{m}$*t*a*c$^{m}$*c$^{m}$*t*c$^{m}$*+C$^{m}$*+U*+U*+C$^{m}$*+A | 5-9-5 | 18 | 18 |
| **ASO 6-4** | Gen 3 | +U*+G+C$^{m}$+U+C$^{m}$*g*t*c$^{m}$*t*a*c$^{m}$*c$^{m}$*t*c$^{m}$*+C$^{m}$+U+U+C$^{m}$*+A | 5-9-5 | 12 | 12 |
| **ASO 7-1** | Gen 3 | +G*+C$^{m}$*+U*c$^{m}$*g*t*c$^{m}$*t*a*c$^{m}$*c$^{m}$*t*c$^{m}$*+C$^{m}$*+U*+U | 3-10-3 | 15 | 15 |
| **ASO 7-2** | Gen 3 | +G*@C$^{m}$*+U*c$^{m}$*g*t*c$^{m}$*t*a*c$^{m}$*c$^{m}$*t*c$^{m}$*+C$^{m}$*@U*+U | 3-10-3 | 15 | 15 |
| **ASO 7-3** | Gen 3 | +U*+G*+C$^{m}$*+U*c$^{m}$*g*t*c$^{m}$*t*a*c$^{m}$*c$^{m}$*t*c$^{m}$*+C$^{m}$*+U*+U*+C$^{m}$ | 4-10-4 | 17 | 17 |
| **ASO 7-4** | Gen 3 | +U*+G+C$^{m}$+U*c$^{m}$*g*t*c$^{m}$*t*a*c$^{m}$*c$^{m}$*t*c$^{m}$*+C$^{m}$+U+U*+C$^{m}$ | 4-10-4 | 13 | 13 |
| **ASO 8-1** | Gen 3 | +U*+G*+C$^{m}$*t*c$^{m}$*g*t*c$^{m}$*t*a*c$^{m}$*c$^{m}$*t*+C$^{m}$*+C$^{m}$*+U | 3-10-3 | 15 | 15 |
| **ASO 8-2** | Gen 3 | +U*@G+C$^{m}$*t*c$^{m}$*g*t*c$^{m}$*t*a*c$^{m}$*c$^{m}$*t*+C$^{m}$@C$^{m}$*U | 3-10-3 | 13 | 13 |
| **ASO 8-5 **** | Gen 3 | +U*+C$^{m}$*+U*+G*+C$^{m}$*t*c$^{m}$*g*t*c$^{m}$*t*a*c$^{m}$*c$^{m}$*t*+C$^{m}$*+C$^{m}$*+U*+U*+C$^{m}$ | 5-10-5 | 19 | 19 |
| **ASO 8-5**** | Gen 3 | +U*+C$^{m}$+U+G+C$^{m}$*t*c$^{m}$*g*t*c$^{m}$*t*a*c$^{m}$*c$^{m}$*t*+C$^{m}$+C$^{m}$+U+U*+C$^{m}$ | 5-10-5 | 13 | 13 |
| **ASO 18-1** | Gen 3 | +U*+G*+C$^{m}$*t*c$^{m}$*g*t*c$^{m}$*t*a*c$^{m}$*c$^{m}$*+U*+C$^{m}$*+C$^{m}$ | 3-10-3 | 14 | 14 |
| **ASO 18-2** | Gen 3 | @U*@G*@C$^{m}$*t*c$^{m}$*g*t*c$^{m}$*t*a*c$^{m}$*c$^{m}$*@U*@C$^{m}$*@C$^{m}$ | 3-10-3 | 14 | 14 |

| | | | | | |
|---|---|---|---|---|---|
| **ASO 18-3** | Gen 3 | +C$^{m}$*+U*+G*+C$^{m}$*t*c$^{m}$*g*t*c$^{m}$*t* a*c$^{m}$*c$^{m}$*+U*+C$^{m}$*+C$^{m}$*+U | 4-9-4 | 16 | 16 |
| **6-5** | Gen 4 | +G+C$^{m}$+U+C$^{m}$*g*t*c$^{m}$*t*a*c$^{m}$*c$^{m}$*t* c$^{m}$*+C$^{m}$+U+U+C$^{m}$ | 4-9-4 | 17 | 10 |
| **7-5** | Gen 4 | +G+C$^{m}$+Uc$^{m}$*g*t*c$^{m}$*t*a*c$^{m}$*c$^{m}$*t*c$^{m}$ *+C$^{m}$+U+U | 3-10-3 | 16 | 10 |
| **7-6** | Gen 4 | +U+G+C$^{m}$+Uc$^{m}$*g*t*c$^{m}$*t*a*c$^{m}$*c$^{m}$*t *c$^{m}$*+C$^{m}$+U+U+C$^{m}$ | 4-10-4 | 18 | 10 |
| **8-6** | Gen 4 | +U+G+C$^{m}$t*c$^{m}$*g*t*c$^{m}$*t*a*c$^{m}$*c$^{m}$*t* +C$^{m}$+C$^{m}$+U | 3-10-3 | 16 | 10 |
| **18-5** | Gen 4 | +U+G+C$^{m}$t*c$^{m}$*g*t*c$^{m}$*t*a*c$^{m}$*c$^{m}$*+ U+C$^{m}$+C$^{m}$ | 3-9-3 | 15 | 9 |
| **NTC (Scrambled ASO)** | Gen 4 | +T*+A *+* C$^{m}$g* c$^{m}$ *g* a* c$^{m}$ *t* t*a *t* +G*+C$^{m}$ *+G* +G | 3-9-4 | 16 | 10 |

Key: X*X = Phosphorothioate linkage; + = MOE; m=methyl; a c g t = DNA; A G C U = RNA; @ LNA

**Supplemental Table 2**: **Mouse-specific ASO screening via gymnosis on mouse cortical neurons (% Tubb4a downregulation determined by qRT-PCR).**

| # | ASO# | 1μM | 5μM | Apotox assay and visual observation |
|---|---|---|---|---|
| 1 | ASO 1 | 44.44±0.2 | 46.25±2.3 | No apoptosis |
| 2 | ASO 2 | 51.78±2.8 | 64.59±4.0 | No apoptosis |
| 3 | ASO 3 | 40.99±4.5 | 54.29±1.2 | No apoptosis |
| 4 | ASO 4 | 34.06±5.1 | 68.69±2.9 | No apoptosis |
| 5 | ASO 5 | -23.76±7.6 | 0.23±1.4 | No apoptosis |
| **6** | **ASO 6** | **70.0±3.2** | **95.4±1.6** | **No apoptosis** |
| **7** | **ASO 7** | **65.2±1.5** | **85.7±1.4** | **No apoptosis** |
| 8 | ASO 4A | 54.38±6.0 | 61.31±6.1 | No apoptosis |
| 9 | ASO 4B | 50.42±8.1 | 62.43±1.3 | No apoptosis |
| 10 | ASO 6A | 15.84±14.9 | 51.03±6.6 | No apoptosis |
| 11 | ASO 6B | 24.74±5.3 | 62.76±5.9 | No apoptosis |
| 12 | ASO 7A | 23.33±2.3 | 62.29±1.2 | No apoptosis |
| 13 | ASO 7B | 58.92±10.8 | 66.67±4.6 | No apoptosis |
| 14 | ASO 8 | 77.4±0.6 | 95.0±1.5 | Highly apoptotic |
| 15 | ASO 9 | 46.50±0.6 | 75.27±0.6 | No apoptosis |
| 16 | ASO 10 | 0.07±12.0 | 56.7±2.3 | No apoptosis |
| 17 | ASO 11 | 8.56±10.6 | 42.73±4.1 | No apoptosis |
| 18 | ASO 12 | | 61.58±4.5 | No apoptosis |
| 19 | ASO 13 | 32.4±16.0 | 61.9±6.2 | No apoptosis |
| 20 | ASO 14 | 8.99±2.1 | 38.19±1.3 | No apoptosis |
| 21 | ASO 15 | 28.25±1.9 | 51.46±2.4 | No apoptosis |
| 22 | ASO 16 | 63.65±0.3 | 79.77±0.5 | No apoptosis |
| 23 | ASO 17 | 37.67±1.1 | 64.46±2.7 | No apoptosis |
| 24 | ASO 18 | 61.52±4.1 | 89.44±0.5 | Highly apoptotic |
| 25 | ASO 19 | 18.37±3.1 | 49.00±4.9 | No apoptosis |
| 26 | ASO 20 | 26.59±6.6 | 48.69±3.5 | No apoptosis |
| 27 | ASO 21 | 38.84±2.3 | 64.95±1.4 | No apoptosis |
| 28 | ASO 22 | 35.19±4.5 | 71.21±3.0 | No apoptosis |
| 29 | ASO 23 | 23.99±2.9 | 67.58±5.1 | No apoptosis |
| 30 | ASO 6-1 | 23±1.01 | 56.6±1.5 | No apoptosis |
| 31 | ASO 6-2 | 32.9±1.5 | 71.7±4.3 | No apoptosis |
| 32 | ASO 6-3 | 26.9±1.3 | 62.9±2.7 | No apoptosis |
| 33 | ASO 6-4 | 0.08±1.01 | 56.3±0.9 | No apoptosis |
| 34 | ASO 7-1 | 40.0±1.8 | 86.3±4.3 | No apoptosis |
| **35** | **ASO 7-2** | **86.9±1.6** | **97.3±1.0** | **No apoptosis** |
| **36** | **ASO 7-3** | **58.8±2.2** | **96.2±0.7** | **No apoptosis** |
| 37 | ASO 7-4 | 52.6±5.4 | 68.9±6.4 | No apoptosis |
| 38 | ASO 8-1 | 58.3±7.6 | 64.3±8.7 | No apoptosis |
| **39** | **ASO 8-2** | **73.4±4.3** | **94.5±4.7** | **No apoptosis** |
| **40** | **ASO 8-3** | **66.1±1.2** | **87.0±2.7** | **No apoptosis** |
| 41 | ASO 8-4 | 17±10.8 | 54.7±8.9 | No apoptosis |
| **42** | **ASO 18-1** | **70.7±7.2** | **87.1±6.3** | **No apoptosis** |
| 43 | ASO 18-2 | 68.9±0.8 | 98.3±0.2 | Highly apoptotic |

| | | | | |
|---|---|---|---|---|
| 44 | ASO 18-3 | 66.4±2.9 | 85.8±4.8 | No apoptosis |
| 45 | ASO 299 | 30.1±1.9 | 35.1±2.9 | No apoptosis |
| 46 | ASO 1029 | 28.8±1.5 | 30.4±2.4 | No apoptosis |
| 47 | ASO 1316 | 24.5±2.2 | 50.4±2.6 | No apoptosis |
| 48 | ASO 1490 | 30.5±2.2 | 34.6±4.9 | No apoptosis |
| 49 | ASO 1747 | 26.4±8.9 | 48.6±2.6 | No apoptosis |
| 50 | ASO 1851 | 23.1±2.1 | 26.5±4.9 | No apoptosis |
| 51 | ASO 2080 | 24.8±1.8 | 20.5±0.9 | No apoptosis |
| 52 | ASO 1908 | 16.8±10.9 | 25.4±2.9 | No apoptosis |
| 53 | ASO 1988 | 8.8±0.9 | 9.8±0.3 | No apoptosis |

Selected ASOs are in "bold". Values are provided as Mean±SEM.

**Supplemental Table 3: % survival after 24h of ASO i.c.v. injection in wild-type pups (n=5 pups for each ASO) at 15μg/g ASO dose.**

| | **ASO 4*** | **ASO 6*** | **ASO 7*** | **ASO 7-2,** | **ASO 7-3** | **ASO 8-3** | **ASO 18-1** | **ASO 18-3** |
|---|---|---|---|---|---|---|---|---|
| **% survival in pups** | 0% | 50% | 50% | 100% | 100% | 100% | 100% | 100% |

*Pups were noted dead the next day in the cage, but PBS-injected mice survived in the same cohort.

**Supplemental Table 4: FOB at 30 days post-ASO injection in wild-type pups injected at P0-P2 (n=5 mice/group).**

| | ASO 7-2 (15μg/g) | ASO 7-3 (15μg/g) | ASO 8-3 (15μg/g) | ASO 18-1(15μg/g) | ASO 18-3(15μg/g) |
|---|---|---|---|---|---|
| **Ataxia** | Present | Absent | Absent | Present | Absent |
| **Hindlimb grasp** | Abnormal | Absent | Absent | Absent | Absent |
| **Body tone** | Abnormal | Absent | Absent | Absent | Absent |
| **Involuntary Movements** | erratic | Absent | Absent | Absent | Absent |
| **Dystonia** | Present | Absent | Absent | Absent | Absent |
| **Tremor** | Present | Absent | Absent | Absent | Absent |
| **Grooming** | Absent | Absent | Absent | Absent | Absent |
| **Hunching** | Hunched, poor trunk, slow movements | Absent | Absent | Absent | Absent |
| **Seizure** | Absent | Absent | Absent | Absent | Absent |
| **Exhibits Jumping Behavior** | Present | Absent | Absent | Absent | Absent |
| **Head Bobbing** | Present | Absent | Absent | Absent | Absent |
| **Head Flicking** | Present | Absent | Absent | Absent | Absent |
| **Tremor** | Present | Absent | Absent | Absent | Absent |
| **Abnormal Gait: ataxic** | Present | Mild | Mild | Present | Absent |
| **Abnormal Gait: hypotonic** | Present | Mild | Mild | Present | Absent |
| **Abnormal Gait: impaired** | Present | Mild | Mild | Present | Absent |
| **Hyperactive** | Present | Present | Present | Present | Absent |
| **Pupil dilator** | Absent | Absent | Absent | Absent | Absent |
| **Piloerection** | Absent | Absent | Absent | Absent | Absent |
| **Righting reflex** | Normal | Normal | Normal | Normal | Normal |
| **Weight loss** | Present | Present | Present | Present | Absent |

**Supplemental Table 5: Antibody resource**

| Antibody | Type/ Species | Source | Identifiers | Additional Information |
|---|---|---|---|---|
| Anti-Tubb4a | Rabbit | abcam | ab179509 | Western Blot Dilution 1:2000<br>IHC Dilution 1:250 and 1:500 |
| Anti-NG2 | Mouse | Millipore Sigma | MAB5384-I | IHC Dilution 1:250 |
| Anti-NG2 | Rabbit | US Biological | C5067-70D | IHC Dilution 1:250 |
| Anti-Olig2 | Mouse | Millipore Sigma | MABN50 | IHC Dilution 1:250 |
| Anti-Iba1 | Rabbit | FUJIFILM Wako Pure Chemical Corporation | 019-19741 | IHC Dilution 1:250 |
| Anti-ASPA | Rabbit | Genetex | GTX110699 | IHC Dilution 1:250 |
| Anti-NeuN | Mouse | Millipore Sigma | MAB377 | IHC Dilution 1:250 and 1:500 |
| Anti-MBP | Rabbit | Obtained from Dr. Judy Grinspan | Hybridoma source | IHC Dilution 1:1 |
| Anti-PLP | Rat | Obtained from Dr. Judy Grinspan | Hybridoma source | IHC Dilution 1:1 |
| Anti-Ki67 | Rabbit | BD Biosciences | 550609 | |
| Anti-GFAP | Rat | Obtained from Dr. Judy Grinspan | Hybridoma source | IHC Dilution 1:1 |
| Anti-Caspase | Rabbit | Cell Signaling Technology | 9579S | IHC Dilution 1:200 |
| Anti-CTIP2 | Rat | abcam | ab18465 | IHC Dilution 1:200 |
| Anti-GAPDH | Mouse | Millipore Sigma | MAB374 | Western Blot Dilution 1:5000 |

**Supplemental Table 6: p-values for genotype and age interaction**

| **Behavior** | **Figure 1C** | **Figure 1D** | **Figure 1E** | **Figure 1F** |
|---|---|---|---|---|
| Genotype Interaction | Tremor | Rotarod | GS Forelimb | GS Hindlimb |
| | P21 | | | |
| WT vs. *Tubb4a*$^{KO/KO}$ | 0.2210 | - | - | - |
| WT vs. *Tubb4a*$^{D249N/+}$ | 0.2523 | - | - | - |
| WT vs. *Tubb4a*$^{D249N/KO}$ | 0.8665 | - | - | - |
| WT vs. *Tubb4a*$^{D249N/D249N}$ | **<0.0001** | - | - | - |
| *Tubb4a*$^{KO/KO}$ vs. *Tubb4a*$^{D249N/+}$ | 0.0265 | - | - | - |
| *Tubb4a*$^{KO/KO}$ vs. *Tubb4a*$^{D249N/KO}$ | 0.1036 | - | - | - |
| *Tubb4a*$^{KO/KO}$ vs. *Tubb4a*$^{D249N/D249N}$ | **<0.0001** | - | - | - |
| | P30 | | | |
| WT vs. *Tubb4a*$^{KO/KO}$ | 0.9367 | 0.6466 | 0.8593 | 0.9998 |
| WT vs. *Tubb4a*$^{D249N/+}$ | 0.9579 | 0.2789 | 0.0994 | 0.0860 |
| WT vs. *Tubb4a*$^{D249N/KO}$ | **0.023** | 0.4294 | 0.9991 | 0.9995 |
| WT vs. *Tubb4a*$^{D249N/D249N}$ | 0.1364 | **<0.0001** | **<0.0001** | **0.036** |
| *Tubb4a*$^{KO/KO}$ vs. *Tubb4a*$^{D249N/+}$ | 0.9998 | 0.1266 | 0.0220 | 0.0362 |
| *Tubb4a*$^{KO/KO}$ vs. *Tubb4a*$^{D249N/KO}$ | **0.0417** | 0.1124 | 0.9867 | >0.9999 |
| *Tubb4a*$^{KO/KO}$ vs. *Tubb4a*$^{D249N/D249N}$ | 0.3137 | **<0.0001** | **<0.0001** | **0.0047** |
| | P60 | | | |
| WT vs. *Tubb4a*$^{KO/KO}$ | 0.9967 | 0.9864 | 0.8919 | 0.7837 |
| WT vs. *Tubb4a*$^{D249N/+}$ | 0.1121 | 0.9595 | 0.6359 | >0.9999 |
| WT vs. *Tubb4a*$^{D249N/KO}$ | **<0.0001** | **0.0011** | 0.4123 | **0.02** |
| WT vs. *Tubb4a*$^{D249N/D249N}$ | - | - | - | - |
| *Tubb4a*$^{KO/KO}$ vs. *Tubb4a*$^{D249N/+}$ | 0.2042 | 0.9120 | 0.2247 | 0.5754 |
| *Tubb4a*$^{KO/KO}$ vs. *Tubb4a*$^{D249N/KO}$ | **<0.0001** | **0.0023** | 0.7958 | 0.2252 |
| *Tubb4a*$^{KO/KO}$ vs. *Tubb4a*$^{D249N/D249N}$ | - | - | - | - |
| | P90 | | | |
| WT vs. *Tubb4a*$^{KO/KO}$ | >0.9999 | 0.6472 | 0.6146 | 0.9987 |
| WT vs. *Tubb4a*$^{D249N/+}$ | 0.1067 | 0.8097 | 0.1701 | 0.2200 |
| WT vs. *Tubb4a*$^{D249N/KO}$ | **<0.0001** | **0.0019** | **0.0350** | **0.0002** |
| WT vs. *Tubb4a*$^{D249N/D249N}$ | - | - | - | - |
| *Tubb4a*$^{KO/KO}$ vs. *Tubb4a*$^{D249N/+}$ | 0.1069 | 0.1716 | 0.170 | 0.4298 |
| *Tubb4a*$^{KO/KO}$ vs. *Tubb4a*$^{D249N/KO}$ | **<0.0001** | **0.0058** | 0.3617 | **0.0008** |
| *Tubb4a*$^{KO/KO}$ vs. *Tubb4a*$^{D249N/D249N}$ | - | - | - | - |
| Behavior | **Figure 1C** | **Figure 1D** | **Figure 1E** | **Figure 1F** |
| Age Interaction | Tremor | Rotarod | GS Forelimb | GS Hindlimb |
| | WT | | | |
| P21 vs. P30 | 0.0018 | - | - | - |
| P21 vs. P60 | 0.0012 | - | - | - |
| P21 vs. P90 | 0.0270 | - | - | - |
| P30 vs. P60 | 0.9998 | 0.9089 | 0.0011 | 0.0908 |
| P30 vs. P90 | 0.9995 | 0.6252 | 0.0024 | 0.1234 |
| P60 vs. P90 | 0.9959 | 0.5683 | 0.5021 | 0.9478 |
| | *Tubb4a*$^{KO/KO}$ | | | |
| P21 vs. P30 | 0.0074 | - | - | - |
| P21 vs. P60 | 0.0002 | - | - | - |
| P21 vs. P90 | 0.0120 | - | - | - |

| | | | | |
|---|---|---|---|---|
| P30 vs. P60 | 0.8947 | 0.3995 | 0.0016 | 0.6728 |
| P30 vs. P90 | 0.6751 | 0.0143 | 0.0372 | 0.1498 |
| P60 vs. P90 | 0.9574 | 0.0633 | 0.9239 | 0.0322 |
| | *Tubb4a*$^{D249N/KO}$ | | | |
| P21 vs. P30 | 0.0071 | - | - | - |
| P21 vs. P60 | <0.0001 | - | - | - |
| P21 vs. P90 | <0.0001 | - | - | - |
| P30 vs. P60 | 0.0055 | <0.0001 | 0.6501 | 0.1037 |
| P30 vs. P90 | 0.0378 | 0.0015 | 0.8817 | 0.0005 |
| P60 vs. P90 | 0.9972 | 0.3305 | 0.4432 | 0.0732 |
| | *Tubb4a*$^{D249N/+}$ | | | |
| P21 vs. P30 | 0.5554 | - | - | - |
| P21 vs. P60 | 0.0147 | - | - | - |
| P21 vs. P90 | 0.0494 | - | - | - |
| P30 vs. P60 | 0.0557 | 0.0983 | 0.0592 | 0.7581 |
| P30 vs. P90 | 0.0826 | 0.1069 | 0.0010 | 0.4221 |
| P60 vs. P90 | 0.6871 | 0.9440 | 0.1688 | 0.1609 |

**Supplemental Table 7: p-values for genotype and age of Figure 1G (Seizure score)**

| **Genotype Interaction** | **P34** | **P35** | **P36** | **P108** | **P109** | **P110** |
|---|---|---|---|---|---|---|
| WT vs. *Tubb4a*$^{D249N/KO}$ | - | - | - | **0.0132** | **0.0335** | 0.1364 |
| WT vs. *Tubb4a*$^{D249N/D249N}$ | 0.0701 | 0.2343 | **0.0439** | - | - | - |
| *Tubb4a*$^{KO/KO}$ vs. *Tubb4a*$^{D249N/KO}$ | - | - | - | **0.0132** | **0.0335** | 0.1364 |

| Age Interaction | ***Tubb4a*$^{D249N/KO}$** | ***Tubb4a*$^{D249N/D249N}$** |
|---|---|---|
| P34 vs. P35 | - | 0.3964 |
| P34 vs. P36 | - | 0.0781 |
| P34 vs. P108 | 0.0164 | - |
| P34 vs. P109 | 0.0415 | - |
| P34 vs. P110 | 0.1655 | - |
| P35 vs. P36 | - | 0.2916 |
| P35 vs. P108 | 0.0164 | - |
| P35 vs. P109 | 0.0415 | - |
| P35 vs. P110 | 0.1655 | - |
| P36 vs. P108 | 0.0164 | - |
| P36 vs. P109 | 0.0415 | - |
| P36 vs. P110 | 0.1655 | - |
| P108 vs. P109 | 0.4300 | - |
| P108 vs. P110 | 0.3487 | - |
| P109 vs. P110 | 0.4215 | - |

**Supplemental Table 8: p-values for Figure 1 and associated Supplemental Tables**

| EM | **Figure 1I** | **Figure 1J** | **Supp Fig 5B** | **Supp Fig 5C** |
|---|---|---|---|---|
| **Genotype Interaction** | % Unmyelinated axons_corpus callosum | g-ratio_ corpus callosum | % Unmyelinated axons_Optic nerve | g-ratio_ Optic nerve |
| | P32-37 | | | |
| WT vs. *Tubb4a*$^{KO/KO}$ | >0.9999 | 0.9954 | >0.9999 | >0.9999 |
| WT vs. *Tubb4a*$^{D249N/+}$ | 0.9988 | **0.0239** | 0.5237 | **0.0003** |
| WT vs. *Tubb4a*$^{D249N/KO}$ | **<0.0001** | **<0.0001** | **<0.0001** | **<0.0001** |
| WT vs. *Tubb4a*$^{D249N/D249N}$ | **<0.0001** | **<0.0001** | **<0.0001** | **<0.0001** |
| *Tubb4a*$^{KO/KO}$ vs. *Tubb4a*$^{D249N/+}$ | >0.9999 | 0.1694 | 0.5335 | **0.0002** |
| *Tubb4a*$^{KO/KO}$ vs. *Tubb4a*$^{D249N/KO}$ | **<0.0001** | **<0.0001** | **<0.0001** | **<0.0001** |
| *Tubb4a*$^{KO/KO}$ vs. *Tubb4a*$^{D249N/D249N}$ | **<0.0001** | **<0.0001** | **<0.0001** | **<0.0001** |
| | P108-P110 | | | |
| WT vs. *Tubb4a*$^{KO/KO}$ | >0.9999 | 0.3555 | >0.9999 | >0.9999 |
| WT vs. *Tubb4a*$^{D249N/+}$ | 0.9988 | **<0.0001** | 0.5237 | **0.0003** |
| WT vs. *Tubb4a*$^{D249N/KO}$ | **<0.0001** | **<0.0001** | **<0.0001** | **<0.0001** |
| WT vs. *Tubb4a*$^{D249N/D249N}$ | Deceased | | | |
| *Tubb4a*$^{KO/KO}$ vs. *Tubb4a*$^{D249N/+}$ | >0.9999 | **0.0021** | >0.9999 | **0.0002** |
| *Tubb4a*$^{KO/KO}$ vs. *Tubb4a*$^{D249N/KO}$ | **<0.0001** | **<0.0001** | **<0.0001** | **<0.0001** |
| *Tubb4a*$^{KO/KO}$ vs. *Tubb4a*$^{D249N/D249N}$ | Deceased | | | |
| EM | **Figure 1I** | **Supp Fig 1K** | **Supp Fig 5B** | **Supp Fig 5C** |
| Age Interaction | % Unmyelinated axons_corpus callosum | g-ratio_ corpus callosum | % Unmyelinated axons_Optic nerve | g-ratio_ Optic nerve |
| | WT | | | |
| P32-37 vs. P110 | **0.004** | 0.2727 | 0.9924 | 0.2988 |
| | *Tubb4a*$^{KO/KO}$ | | | |
| P32-37 vs. P110 | **0.0003** | **0.005** | 0.2325 | **0.01** |
| | *Tubb4a*$^{D249N/KO}$ | | | |
| P32-37 vs. P110 | **<0.0001** | **<0.0001** | **<0.0001** | **<0.0001** |
| | *Tubb4a*$^{D249N/+}$ | | | |
| P32-37 vs. P110 | 0.457 | **0.0008** | 0.4439 | 0.182 |

| Supplemental Table 9: p-values for Supplemental Figure 1 | | | | |
|---|---|---|---|---|
| **Behavior** | **Supp Fig 1A** | **Supp Fig 1B** | **Supp Fig 1C** | **Supp Fig 1D** |
| Genotype & Age Interaction | Tremor | Rotarod | GS Forelimb | GS Hindlimb |
| | | 6 months | | |
| WT vs. *Tubb4a*$^{KO/KO}$ | 0.0337 | 0.4804 | 0.3536 | 0.0219 |
| | | 12 months | | |
| WT vs. *Tubb4a*$^{KO/KO}$ | 0.0237 | 0.5399 | 0.1879 | 0.9541 |
| | | WT | | |
| 6m vs. 12m | 0.2663 | **0.0199** | 0.9444 | **0.0214** |
| | | *Tubb4a*$^{KO/KO}$ | | |
| 6m vs. 12m | 0.6185 | **0.0276** | 0.5378 | **0.0001** |

| Supplemental Table 10: p-values for Western blot associated with Figure 1 | | | |
|---|---|---|---|
| Immunoblot analysis of extracted myelin fraction | **Figure 1J** | **Supp Figure 4 I&J** | **Supp Figure 4 G&H** |
| | MBP band density | CNP band density | PLP band density |
| | | P32-P37 | |
| WT vs. *Tubb4a*$^{D249N/+}$ | 0.8558 | 0.3967 | 0.2852 |
| WT vs. *Tubb4a*$^{D249N/KO}$ | **0.0029** | 0.1050 | **0.0128** |
| WT vs. *Tubb4a*$^{D249N/D249N}$ | **0.0001** | **0.0076** | **0.0228** |
| | | P108-P110 | |
| WT vs. *Tubb4a*$^{D249N/+}$ | 0.9518 | 0.8956 | 0.6556 |
| WT vs. *Tubb4a*$^{D249N/KO}$ | **0.001** | **0.0172** | **0.0277** |
| WT vs. *Tubb4a*$^{D249N/D249N}$ | - | - | - |

| Supplemental Table 11: p-values for Supplemental Figure 2 | | | | |
|---|---|---|---|---|
| **Histology–Genotype Interaction** | **P21** | **P32-37** | **P60** | **P108-110** |
| MBP Density (**Supplemental Figure 2C**) | | | | |
| Corpus callosum | | | | |
| WT vs. *Tubb4a*$^{KO/KO}$ | 0.7264 | 0.9979 | 0.9998 | 0.8139 |
| WT vs. *Tubb4a*$^{D249N/+}$ | 0.9382 | 0.9997 | 0.9043 | >0.9999 |
| WT vs. *Tubb4a*$^{D249N/KO}$ | 0.9377 | **0.024** | **0.049** | **0.0500** |
| WT vs. *Tubb4a*$^{D249N/D249N}$ | **<0.0001** | **0.0083** | - | - |
| *Tubb4a*$^{KO/KO}$ vs. *Tubb4a*$^{D249N/+}$ | 0.9969 | 0.5468 | 0.6581 | 0.6552 |
| *Tubb4a*$^{KO/KO}$ vs. *Tubb4a*$^{D249N/KO}$ | 0.9969 | 0.2526 | 0.1384 | 0.0215 |
| *Tubb4a*$^{KO/KO}$ vs. *Tubb4a*$^{D249N/D249N}$ | 0.0671 | 0.0063 | - | - |
| Eri-C quantification (**Supplemental Figure 2D**) | | | | |
| Corpus callosum | | | | |
| WT vs. *Tubb4a*$^{KO/KO}$ | 0.8016 | 0.4503 | 0.8790 | 0.9980 |
| WT vs. *Tubb4a*$^{D249N/+}$ | 0.7725 | 0.0896 | **0.0030** | **0.0326** |
| WT vs. *Tubb4a*$^{D249N/KO}$ | 0.0867 | **<0.0001** | **<0.0001** | **<0.0001** |
| WT vs. *Tubb4a*$^{D249N/D249N}$ | **<0.0001** | **<0.0001** | - | - |
| *Tubb4a*$^{KO/KO}$ vs. *Tubb4a*$^{D249N/+}$ | 0.6803 | 0.1053 | 0.0025 | 0.0677 |
| *Tubb4a*$^{KO/KO}$ vs. *Tubb4a*$^{D249N/KO}$ | 0.0847 | **<0.0001** | **<0.0001** | **0.0094** |
| *Tubb4a*$^{KO/KO}$ vs. *Tubb4a*$^{D249N/D249N}$ | **<0.0001** | **<0.0001** | - | - |

| Age Interaction | **MBP Density** | **Eri-C quantification** |
|---|---|---|
| WT | | |
| P21 vs. P32-37 | 0.6066 | 0.0193 |
| P21 vs. P60 | 0.4967 | 0.0172 |
| P21 vs. P108-110 | 0.2138 | 0.2662 |
| P32-27 vs. P60 | 0.9190 | 0.1667 |
| P32-37 vs. P108-110 | 0.3824 | 0.8169 |
| P60 vs. P108-110 | 0.6805 | 0.7950 |
| *Tubb4a*$^{KO/KO}$ | | |
| P21 vs. P32-37 | 0.0266 | 0.0556 |
| P21 vs. P60 | 0.2741 | **0.0168** |
| P21 vs. P108-110 | 0.0141 | 0.6242 |
| P32-27 vs. P60 | 0.8288 | 0.2766 |
| P32-37 vs. P108-110 | 0.0146 | 0.9381 |
| P60 vs. P108-110 | 0.2156 | 0.7399 |
| *Tubb4a*$^{D249N/KO}$ | | |
| P21 vs. P32-37 | 0.8984 | 0.5443 |
| P21 vs. P60 | 0.9937 | 0.8578 |
| P21 vs. P108-110 | 0.6986 | 0.8163 |
| P32-27 vs. P60 | 0.9271 | **0.0080** |
| P32-37 vs. P108-110 | 0.9827 | **0.05** |
| P60 vs. P108-110 | 0.7059 | 0.8622 |
| *Tubb4a*$^{D249N/+}$ | | |
| P21 vs. P32-37 | 0.0998 | 0.1771 |
| P21 vs. P60 | 0.0445 | 0.5833 |
| P21 vs. P108-110 | **0.0056** | 0.6282 |
| P32-27 vs. P60 | 0.1857 | 0.4954 |
| P32-37 vs. P108-110 | 0.0312 | 0.1533 |
| P60 vs. P108-110 | 0.7329 | 0.8342 |

| Supplemental Table 12: p-values for Supplemental Figure 3 | | | | |
|---|---|---|---|---|
| Histology–Genotype Interaction | **P21** | **P32-37** | **P60** | **P108-110** |
| MBP Density | | | | |
| Cerebellum | | | | |
| WT vs. *Tubb4a*$^{KO/KO}$ | 0.9334 | 0.6992 | 0.9595 | 0.9790 |
| WT vs. *Tubb4a*$^{D249N/+}$ | 0.9933 | 0.8425 | 0.9995 | 0.5380 |
| WT vs. *Tubb4a*$^{D249N/KO}$ | 0.1767 | **0.0075** | **0.023** | **0.0442** |
| WT vs. *Tubb4a*$^{D249N/D249N}$ | **0.0445** | **0.0121** | - | - |
| *Tubb4a*$^{KO/KO}$ vs. *Tubb4a*$^{D249N/+}$ | >0.9999 | >0.9999 | 0.9993 | 0.4686 |
| *Tubb4a*$^{KO/KO}$ vs. *Tubb4a*$^{D249N/KO}$ | **0.0080** | **0.06150** | 0.5421 | 0.4393 |
| *Tubb4a*$^{KO/KO}$ vs. *Tubb4a*$^{D249N/D249N}$ | **0.0117** | **0.0572** | - | - |
| Eri-C quantification | | | | |
| Cerebellum | | | | |
| WT vs. *Tubb4a*$^{KO/KO}$ | 0.8592 | 0.6043 | 0.9994 | 0.9673 |
| WT vs. *Tubb4a*$^{D249N/+}$ | 0.7016 | 0.1901 | **0.0191** | 0.0718 |
| WT vs. *Tubb4a*$^{D249N/KO}$ | **0.0343** | **0.0062** | **<0.0001** | **0.0022** |
| WT vs. *Tubb4a*$^{D249N/D249N}$ | **<0.0001** | **<0.0001** | - | - |
| *Tubb4a*$^{KO/KO}$ vs. *Tubb4a*$^{D249N/+}$ | 0.9329 | 0.2666 | 0.0230 | 0.4902 |
| *Tubb4a*$^{KO/KO}$ vs. *Tubb4a*$^{D249N/KO}$ | **0.0216** | **0.0014** | **<0.0001** | **0.0323** |
| *Tubb4a*$^{KO/KO}$ vs. *Tubb4a*$^{D249N/D249N}$ | **0.0003** | **<0.0001** | - | - |
| Age Interaction | **MBP Density** | | **Eri-C quantification** | |
| WT | | | | |
| P21 vs. P32-37 | 0.0178 | | 0.0216 | |
| P21 vs. P60 | 0.1887 | | 0.0033 | |
| P21 vs. P108-110 | 0.2666 | | 0.7555 | |
| P32-27 vs. P60 | **0.0035** | | 0.9740 | |
| P32-37 vs. P108-110 | 0.6386 | | 0.7914 | |
| P60 vs. P108-110 | 0.7722 | | 0.7531 | |
| *Tubb4a*$^{KO/KO}$ | | | | |
| P21 vs. P32-37 | 0.9462 | | 0.2477 | |
| P21 vs. P60 | 0.8862 | | 0.2277 | |
| P21 vs. P108-110 | 0.8315 | | 0.9996 | |
| P32-27 vs. P60 | 0.9953 | | 0.5393 | |
| P32-37 vs. P108-110 | 0.9994 | | 0.8340 | |
| P60 vs. P108-110 | 0.9567 | | 0.7879 | |
| *Tubb4a*$^{D249N/KO}$ | | | | |
| P21 vs. P32-37 | 0.7936 | | 0.0920 | |
| P21 vs. P60 | 0.9489 | | 0.5780 | |
| P21 vs. P108-110 | 0.8107 | | 0.9150 | |
| P32-37 vs. P60 | 0.9821 | | **0.0330** | |
| P32-37 vs. P108-110 | 0.9990 | | 0.1298 | |
| P60 vs. P108-110 | 0.9378 | | 0.4038 | |
| *Tubb4a*$^{D249N/+}$ | | | | |
| P21 vs. P32-37 | 0.8598 | | 0.5770 | |
| P21 vs. P60 | 0.9677 | | 0.3932 | |
| P21 vs. P108-110 | 0.4595 | | 0.681 | |
| P32-27 vs. P60 | >0.9999 | | 0.1236 | |
| P32-37 vs. P108-110 | 0.6852 | | **0.0306** | |
| P60 vs. P108-110 | 0.7797 | | 0.9983 | |
| *Tubb4a*$^{D249N/D249N}$ | | | | |
| P21 vs. P32-37 | 0.3038 | | 0.3339 | |
| P21 vs. P60 | - | | - | |

| | | |
|---|---|---|
| P21 vs. P108-110 | - | - |
| P32-27 vs. P60 | - | - |
| P32-37 vs. P108-110 | - | - |
| P60 vs. P108-110 | - | - |

| Supplemental Table 13: p-values for Figure 2 | | | |
|---|---|---|---|
| Histology – ASPA Count/mm$^2$ | **P21** | **P32-P37** | **P108-P110** |
| **Figure 2B**: Corpus Callosum | | | |
| WT vs. *Tubb4a*$^{KO/KO}$ | >0.9999 | 0.6865 | 0.9992 |
| WT vs. *Tubb4a*$^{D249N/+}$ | 0.7141 | 0.6207 | 0.8056 |
| WT vs. *Tubb4a*$^{D249N/KO}$ | 0.2510 | **0.0474** | **0.0464** |
| WT vs. *Tubb4a*$^{D249N/D249N}$ | **0.0002** | **<0.0001** | Deceased |
| *Tubb4a*$^{KO/KO}$ vs. *Tubb4a*$^{D249N/+}$ | 0.8592 | 0.3872 | 0.6632 |
| *Tubb4a*$^{KO/KO}$ vs. *Tubb4a*$^{D249N/KO}$ | 0.2479 | **0.0695** | **0.0366** |
| *Tubb4a*$^{KO/KO}$ vs. *Tubb4a*$^{D249N/D249N}$ | 0.0063 | **0.0400** | Deceased |
| **Figure 2C**: Cerebellum White Matter | | | |
| WT vs. *Tubb4a*$^{KO/KO}$ | - | 0.9955 | 0.1722 |
| WT vs. *Tubb4a*$^{D249N/+}$ | - | >0.9999 | 0.9713 |
| WT vs. *Tubb4a*$^{D249N/KO}$ | - | 0.9925 | **0.0106** |
| WT vs. *Tubb4a*$^{D249N/D249N}$ | - | **0.0309** | Deceased |
| *Tubb4a*$^{KO/KO}$ vs. *Tubb4a*$^{D249N/+}$ | - | >0.9999 | >0.9999 |
| *Tubb4a*$^{KO/KO}$ vs. *Tubb4a*$^{D249N/KO}$ | - | 0.9755 | 0.1845 |
| *Tubb4a*$^{KO/KO}$ vs. *Tubb4a*$^{D249N/D249N}$ | - | 0.1551 | Deceased |
| **Figure 2D**: Cerebellum granule layer | | | |
| WT vs. *Tubb4a*$^{KO/KO}$ | - | 0.6865 | 0.9992 |
| WT vs. *Tubb4a*$^{D249N/+}$ | - | 0.6563 | 0.9998 |
| WT vs. *Tubb4a*$^{D249N/KO}$ | - | 0.9987 | **0.0347** |
| WT vs. *Tubb4a*$^{D249N/D249N}$ | - | **0.0309** | Deceased |
| *Tubb4a*$^{KO/KO}$ vs. *Tubb4a*$^{D249N/+}$ | - | >0.9999 | 0.9940 |
| *Tubb4a*$^{KO/KO}$ vs. *Tubb4a*$^{D249N/KO}$ | - | 0.8322 | 0.0347 |
| *Tubb4a*$^{KO/KO}$ vs. *Tubb4a*$^{D249N/D249N}$ | - | 0.0019 | Deceased |
| Histology – ASPA Count/mm$^2$ | **Figure 2B** | **Figure 2C** | **Figure 2D** |
| Age Interaction | Corpus Callosum | Cerebellum White Matter | Cerebellum granule layer |
| WT | | | |
| P21 vs. P32-37 | 0.1351 | - | - |
| P21 vs. P108-110 | 0.1405 | - | - |
| P32-37 vs. P108-110 | 0.3603 | 0.9819 | 0.4994 |
| *Tubb4a*$^{KO/KO}$ | | | |
| P21 vs. P32-37 | 0.1248 | - | - |
| P21 vs. P108-110 | 0.1358 | - | - |
| P32-37 vs. P108-110 | 0.5134 | 0.4604 | 0.7047 |
| *Tubb4a*$^{D249N/KO}$ | | | |
| P21 vs. P32-37 | 0.9800 | - | - |
| P21 vs. P108-110 | 0.9224 | - | - |
| P32-37 vs. P108-110 | 0.1679 | 0.08 | **0.0202** |
| *Tubb4a*$^{D249N/D249N}$ | | | |
| P21 vs. P32-37 | 0.0710 | - | - |
| P21 vs. P108-110 | Deceased | - | - |
| P32-37 vs. P108-110 | Deceased | Deceased | Deceased |

| | | | |
|---|---|---|---|
| | ***Tubb4a*$^{D249N/+}$** | | |
| **P21 vs. P32-37** | 0.8448 | - | - |
| **P21 vs. P108-110** | 0.2143 | - | - |
| **P32-37 vs. P108-110** | 0.1873 | 0.6997 | 0.4128 |

| **Supplemental Table 14: p-values for Supplemental Figure 7** | | | |
|---|---|---|---|
| **Histology –Olig2 Count/mm$^2$** | **P21** | **P32-P37** | **P108-P110** |
| **Figure S7C**: Corpus Callosum | | | |
| WT vs. *Tubb4a*$^{KO/KO}$ | 0.9997 | 0.2571 | 0.7440 |
| WT vs. *Tubb4a*$^{D249N/+}$ | 0.9775 | 0.4506 | 0.5529 |
| WT vs. *Tubb4a*$^{D249N/KO}$ | 0.8693 | 0.3528 | 0.2299 |
| WT vs. *Tubb4a*$^{D249N/D249N}$ | 0.9771 | >0.9999 | Deceased |
| *Tubb4a*$^{KO/KO}$ vs. *Tubb4a*$^{D249N/+}$ | 0.9369 | 0.8400 | 0.8669 |
| *Tubb4a*$^{KO/KO}$ vs. *Tubb4a*$^{D249N/KO}$ | 0.2269 | 0.9984 | 0.8317 |
| *Tubb4a*$^{KO/KO}$ vs. *Tubb4a*$^{D249N/D249N}$ | 0.9697 | 0.1963 | Deceased |
| **Figure S7D**: Cerebellum White Matter | | | |
| WT vs. *Tubb4a*$^{KO/KO}$ | - | 0.9975 | 0.9695 |
| WT vs. *Tubb4a*$^{D249N/+}$ | - | 0.9970 | 0.6120 |
| WT vs. *Tubb4a*$^{D249N/KO}$ | - | 0.9884 | 0.9992 |
| WT vs. *Tubb4a*$^{D249N/D249N}$ | - | 0.9582 | Deceased |
| *Tubb4a*$^{KO/KO}$ vs. *Tubb4a*$^{D249N/+}$ | - | >0.9999 | 0.2784 |
| *Tubb4a*$^{KO/KO}$ vs. *Tubb4a*$^{D249N/KO}$ | - | 0.9998 | 0.9067 |
| *Tubb4a*$^{KO/KO}$ vs. *Tubb4a*$^{D249N/D249N}$ | - | 0.8444 | Deceased |
| **Figure S7E**: Cerebellum granule layer | | | |
| WT vs. *Tubb4a*$^{KO/KO}$ | - | 0.6803 | 0.9995 |
| WT vs. *Tubb4a*$^{D249N/+}$ | - | 0.2223 | 0.9790 |
| WT vs. *Tubb4a*$^{D249N/KO}$ | - | 0.9975 | 0.6720 |
| WT vs. *Tubb4a*$^{D249N/D249N}$ | - | 0.8877 | Deceased |
| *Tubb4a*$^{KO/KO}$ vs. *Tubb4a*$^{D249N/+}$ | - | 0.9004 | 0.9334 |
| *Tubb4a*$^{KO/KO}$ vs. *Tubb4a*$^{D249N/KO}$ | - | 0.9935 | 0.5420 |
| *Tubb4a*$^{KO/KO}$ vs. *Tubb4a*$^{D249N/D249N}$ | - | 0.3593 | Deceased |
| Histology – ASPA Count/mm$^2$ | **Figure 7C** | **Figure 7D** | **Figure 7E** |
| Age Interaction | Corpus Callosum | Cerebellum White Matter | Cerebellum granule layer |
| | WT | | |
| P21 vs. P32-37 | 0.8999 | - | - |
| P21 vs. P108-110 | 0.8332 | - | - |
| P32-37 vs. P108-110 | 0.9940 | 0.1922 | 0.003 |
| | *Tubb4a*$^{KO/KO}$ | | |
| P21 vs. P32-37 | 0.7225 | - | - |
| P21 vs. P108-110 | 0.8449 | - | - |
| P32-37 vs. P108-110 | 0.4322 | 0.03 | 0.02 |
| | *Tubb4a*$^{D249N/KO}$ | | |
| P21 vs. P32-37 | 0.0145 | - | - |
| P21 vs. P108-110 | 0.5619 | - | - |
| P32-37 vs. P108-110 | 0.9983 | 0.1238 | 0.05 |
| | *Tubb4a*$^{D249N/D249N}$ | | |
| P21 vs. P32-37 | 0.4289 | - | - |
| P21 vs. P108-110 | Deceased | - | - |
| P32-37 vs. P108-110 | Deceased | Deceased | Deceased |
| | ***Tubb4a*$^{D249N/+}$** | | |

| | | | |
|---|---|---|---|
| **P21 vs. P32-37** | 0.8644 | - | - |
| **P21 vs. P108-110** | 0.9580 | - | - |
| **P32-37 vs. P108-110** | 0.9980 | 0.8039 | 0.369 |

| Supplemental Table 15: p-values for Figure 3 | | | |
|---|---|---|---|
| **Histology** | **P32-P37** | **P60** | **P108-P110** |
| **Figure3C:** NeuN profile/mm$^2$ | | | |
| WT vs. *Tubb4a*$^{KO/KO}$ | 0.3879 | 0.5657 | 0.8577 |
| WT vs. *Tubb4a*$^{D249N/+}$ | 0.9985 | 0.6079 | 0.0616 |
| WT vs. *Tubb4a*$^{D249N/KO}$ | 0.8418 | **<0.0001** | **0.0002** |
| WT vs. *Tubb4a*$^{D249N/D249N}$ | **0.00109** | - | - |
| *Tubb4a*$^{KO/KO}$ vs. *Tubb4a*$^{D249N/+}$ | 0.4117 | 0.9237 | 0.9685 |
| *Tubb4a*$^{KO/KO}$ vs. *Tubb4a*$^{D249N/KO}$ | 0.1466 | **0.0475** | **0.0332** |
| *Tubb4a*$^{KO/KO}$ vs. *Tubb4a*$^{D249N/D249N}$ | **0.0100** | - | - |
| **Figure3E:** NeuN+ Caspase+ profile/mm$^2$ | | | |
| WT vs. *Tubb4a*$^{KO/KO}$ | 0.0250 | 0.8410 | 0.9935 |
| WT vs. *Tubb4a*$^{D249N/+}$ | 0.2775 | 0.4759 | 0.3718 |
| WT vs. *Tubb4a*$^{D249N/KO}$ | 0.9924 | **<0.0001** | **0.0013** |
| WT vs. *Tubb4a*$^{D249N/D249N}$ | **<0.0001** | - | - |
| *Tubb4a*$^{KO/KO}$ vs. *Tubb4a*$^{D249N/+}$ | 0.2182 | **<0.0001** | **0.0023** |
| *Tubb4a*$^{KO/KO}$ vs. *Tubb4a*$^{D249N/KO}$ | 0.4864 | **<0.0001** | **0.0013** |
| *Tubb4a*$^{KO/KO}$ vs. *Tubb4a*$^{D249N/D249N}$ | **<0.0001** | - | - |
| **Figure3G:** CTIP2 profile/mm$^2$ | | | |
| WT vs. *Tubb4a*$^{KO/KO}$ | 0.9999 | - | 0.9892 |
| WT vs. *Tubb4a*$^{D249N/+}$ | 0.6390 | - | 0.9766 |
| WT vs. *Tubb4a*$^{D249N/KO}$ | >0.9999 | - | 0.1357 |
| WT vs. *Tubb4a*$^{D249N/D249N}$ | **0.0014** | - | - |
| *Tubb4a*$^{KO/KO}$ vs. *Tubb4a*$^{D249N/+}$ | 0.5528 | - | 0.8298 |
| *Tubb4a*$^{KO/KO}$ vs. *Tubb4a*$^{D249N/KO}$ | 0.9994 | - | 0.0561 |
| *Tubb4a*$^{KO/KO}$ vs. *Tubb4a*$^{D249N/D249N}$ | **0.0019** | - | - |
| **Histology** | **Figure3C** | **Figure3E** | **Figure3G** |
| Age Interaction | NeuN profile/mm$^2$ | NeuN+ Caspase+ profile/mm$^2$ | CTIP2 profile/mm$^2$ |
| WT | | | |
| P32-37 vs. P60 | 0.9496 | 0.8839 | - |
| P32-37 vs. P108-110 | 0.8693 | **0.0359** | **<0.0001** |
| P60 vs. P108-110 | 0.0152 | 0.8745 | - |
| *Tubb4a*$^{KO/KO}$ | | | |
| P32-37 vs. P60 | 0.2433 | 0.6452 | - |
| P32-37 vs. P108-110 | 0.3403 | 0.0621 | <0.0001 |
| P60 vs. P108-110 | 0.8949 | 0.1303 | - |
| *Tubb4a*$^{D249N/KO}$ | | | |
| P32-37 vs. P60 | <0.0001 | 0.0166 | - |
| P32-37 vs. P108-110 | 0.0021 | 0.0941 | 0.0143 |
| P60 vs. P108-110 | 0.0416 | 0.0646 | - |
| *Tubb4a*$^{D249N/+}$ | | | |
| P32-37 vs. P60 | 0.7134 | 0.7154 | - |
| P32-37 vs. P108-110 | 0.1046 | 0.9557 | 0.0037 |
| P60 vs. P108-110 | 0.7579 | 0.6244 | - |

| Supplemental Table 16: p-values for Supplemental Figure 10E | | |
|---|---|---|
| Histology – NeuN Count/mm$^2$ | **P32-P37** | **P108-P110** |
| WT vs. *Tubb4a*$^{KO/KO}$ | 0.6479 | >0.9999 |
| WT vs. *Tubb4a*$^{D249N/+}$ | 0.8320 | 0.9963 |
| WT vs. *Tubb4a*$^{D249N/KO}$ | >0.9999 | 0.4883 |
| WT vs. *Tubb4a*$^{D249N/D249N}$ | **0.023** | - |
| *Tubb4a*$^{KO/KO}$ vs. *Tubb4a*$^{D249N/+}$ | 0.9966 | 0.9949 |
| *Tubb4a*$^{KO/KO}$ vs. *Tubb4a*$^{D249N/KO}$ | 0.6967 | **0.0475** |
| *Tubb4a*$^{KO/KO}$ vs. *Tubb4a*$^{D249N/D249N}$ | **0.00100** | - |
| WT | | |
| P32-37 vs. P108-P110 | 0.5696 | |
| *Tubb4a*$^{KO/KO}$ | | |
| P32-37 vs. P108-P110 | **0.0623** | |
| *Tubb4a*$^{D249N/KO}$ | | |
| P32-37 vs. P108-P110 | **0.0267** | |
| *Tubb4a*$^{D249N/D249N}$ | | |
| P32-37 vs. P108-P110 | Deceased | |
| *Tubb4a*$^{D249N/+}$ | | |
| P32-37 vs. P108-P110 | **0.0578** | |

| Supplemental Table 17: p-values for Figure 5C | | | |
|---|---|---|---|
| **Western Blot** | **Cortex** | **Striatum** | **Cerebellum** |
| p-values with PBS | | | |
| P30 | 0.018 | **0.0007** | 0.2520 |
| **p-values for Figure 5D, E and G** | | | |
| Longitudinal response in WT-qRT-PCR | **Cortex** | **Striatum** | **Cerebellum** |
| p-values compared to PBS | | | |
| PBS | >0.9999 | >0.9999 | >0.9999 |
| P30 | **<0.0001** | **<0.0001** | 0.4945 |
| P60 | **0.0026** | **0.0016** | 0.4099 |
| P90 | **0.0018** | **<0.0001** | 0.0704 |
| P160 | 0.4268 | 0.0859 | >0.9999 |
| Longitudinal response in WT-Western Blot | **Cortex** | **Striatum** | **Cerebellum** |
| p-values compared to PBS | | | |
| P30 | **0.014** | **0.029** | 0.0544 |
| P60 | **0.045** | **0.041** | 0.5558 |
| P90 | 0.9854 | 0.0967 | 0.4747 |
| P160 | 0.9732 | 0.6662 | 0.8405 |
| Figure 5G | **Cortex** | **Striatum** | **Cerebellum** |
| p-values compared to *Tubb4a*$^{D249N/KO}$ PBS | | | |
| *Tubb4a*$^{D249N/KO}$ *ASO* | **0.0006** | **0.0003** | **0.018** |

| Supplemental Table 19: p values for Figure 6C-F (behavior) | | | | |
|---|---|---|---|---|
| **Behavior** | **Tremor (Figure 6C)** | **Rotarod (Figure 6D)** | **Forelimb (Figure 6E)** | **Hindlimb (Figure 6F)** |
| Up to end stage (P30-P110**)** | | | | |
| WT baseline (intercept) | <0.0001 | <0.0001 | <0.0001 | <0.0001 |
| Time days (after P30) | 0.0024 | 0.8360 | <0.0001 | <0.0001 |
| Treatment (ASO) | 0.0033 | 0.7230 | 0.0071 | 0.0012 |
| Genotype (*Tubb4a*$^{D249N/KO}$) | <0.0001 | 0.0000 | 0.1012 | 0.0516 |
| Treatment vs Genotype | 0.0066 | 0.0001 | 0.8275 | 0.5236 |
| Time days (after P30) vs Treatment | 0.9085 | 0.8880 | 0.0500 | 0.1278 |
| Time days (after P30) vs Genotype | 0.5395 | <0.0001 | <0.0001 | <0.0001 |
| Time days (after P30) Treatment vs Genotype | 0.0045 | <0.0001 | 0.0001 | 0.0155 |
| After end stage (P110-P365) *no *Tubb4a*$^{D249N/KO}$ PBS | | | | |
| Time days (after P110) | 0.0006 | <0.0001 | 0.0199 | <0.0001 |
| Treatment | <0.0001 | <0.0001 | 0.0000 | 0.1351 |
| Genotype | 0.3023 | 0.0828 | 0.4909 | 0.0003 |
| Time days (after P110) vs Treatment | 0.2312 | <0.0001 | 0.0368 | 0.1558 |
| Time days (after P110) vs Genotype | <0.0001 | <0.0001 | <0.0001 | <0.0001 |

| Supplemental Table 20: p-values for Figure 8 (EM) | | | |
|---|---|---|---|
| **EM** | **Figure 8C** | **Figure 8D** | **Figure 8E** |
| **Genotype Interaction** | g-ratio_CC | % Unmyelinated axons_CC | Vacuoles #_CC |
| | P48 | | |
| WT PBS vs. WT ASO | 0.7472 | >0.9999 | 0.9922 |
| WT PBS vs. *Tubb4a*$^{D249N/KO}$ PBS | **<0.0001** | **<0.0001** | 0.3402 |
| WT PBS vs. *Tubb4a*$^{D249N/KO}$ ASO | **<0.0001** | **<0.0001** | 0.1190 |
| WT ASO vs. *Tubb4a*$^{D249N/KO}$ PBS | **<0.0001** | **<0.0001** | 0.5011 |
| WT ASO vs. *Tubb4a*$^{D249N/KO}$ ASO | **<0.0001** | **<0.0001** | 0.2098 |
| *Tubb4a*$^{D249N}$PBS vs. *Tubb4a*$^{D249N/KO}$ ASO | **<0.0001** | **<0.0001** | 0.9433 |
| | P108-P110 | | |
| WT PBS vs. WT ASO | 0.9915 | 0.9643 | 0.8322 |
| WT PBS vs. *Tubb4a*$^{D249N/KO}$ PBS | **<0.0001** | **<0.0001** | **<0.0001** |
| WT PBS vs. *Tubb4a*$^{D249N/KO}$ ASO | **<0.0001** | **<0.0001** | **<0.0001** |
| WT ASO vs. *Tubb4a*$^{D249N/KO}$ PBS | **<0.0001** | **<0.0001** | **<0.0001** |
| WT ASO vs. *Tubb4a*$^{D249N/KO}$ ASO | **<0.0001** | **<0.0001** | **<0.0001** |
| *Tubb4a*$^{D249N}$PBS vs. *Tubb4a*$^{D249N/KO}$ ASO | **<0.0001** | **<0.0001** | **<0.0001** |
| Age Interaction | g-ratio_CC | % Unmyelinated axons_CC | Vacuoles #_CC |
| | WT PBS treated | | |
| P48 vs. P108-P110 | 0.7388 | 0.8623 | 0.4043 |
| | WT ASO treated | | |
| P48 vs. P108-P110 | 0.2262 | 0.5152 | 0.7773 |
| | *Tubb4a*$^{D249N}$ PBS treated | | |
| P48 vs. P108-P110 | **0.0350** | **0.018** | **<0.0001** |
| | *Tubb4a*$^{D249N}$ ASO treated | | |
| P48 vs. P108-P110 | 0.8140 | 0.3798 | **<0.0001** |

| Supplemental Table 21: p-values for Figure 8 (Histology) | | |
|---|---|---|
| **Histology** | **Figure 8I (cerebellum)** | **Figure 8J** |
| **Genotype Interaction** | % Eri-C | ASPA profile/mm$^2$ |
| | **P48** | |
| WT PBS vs. WT ASO | 0.1650 | 0.9967 |
| WT PBS vs. *Tubb4a*$^{D249N/KO}$ PBS | **<0.0001** | **0.009** |
| WT PBS vs. *Tubb4a*$^{D249N/KO}$ ASO | **<0.0001** | **0.163** |
| WT ASO vs. *Tubb4a*$^{D249N/KO}$ PBS | **<0.0001** | **0.0014** |
| WT ASO vs. *Tubb4a*$^{D249N/KO}$ ASO | **0.0002** | 0.2295 |
| *Tubb4a*$^{D249N}$PBS vs. *Tubb4a*$^{D249N/KO}$ ASO | **<0.0001** | **0.040** |
| | P108-P110 | |
| WT PBS vs. WT ASO | 0.9915 | 0.9643 |
| WT PBS vs. *Tubb4a*$^{D249N/KO}$ PBS | **<0.0001** | **0.001** |
| WT PBS vs. *Tubb4a*$^{D249N/KO}$ ASO | **<0.0001** | 0.2314 |
| WT ASO vs. *Tubb4a*$^{D249N/KO}$ PBS | **<0.0001** | **0.006** |
| WT ASO vs. *Tubb4a*$^{D249N/KO}$ ASO | **<0.0001** | 0.1273 |
| *Tubb4a*$^{D249N}$PBS vs. *Tubb4a*$^{D249N/KO}$ ASO | **<0.0001** | **0.01** |
| **Age Interaction** | % Eri-C | ASPA profile/mm$^2$ |
| | WT PBS treated | |
| P48 vs. P108-P110 | 0.4641 | 0.8623 |
| | WT ASO treated | |
| P48 vs. P108-P110 | 0.021 | 0.5152 |
| | *Tubb4a*$^{D249N}$ PBS treated | |
| P48 vs. P108-P110 | **<0.0001** | **0.018** |
| | Tubb4a$^{D249N}$ ASO treated | |
| P48 vs. P108-P110 | 0.4473 | 0.3798 |

| Supplemental Table 22: p-values for Histology | | | |
|---|---|---|---|
| **Supplemental Figure 29C** | | **Supplemental Figure 29E** | |
| **Histology – BCAS1 profile/mm$^2$** | **P48** | **Histology – NG2 Olig2 profile/mm$^2$** | **P48** |
| WT PBS vs. WT ASO | 0.2852 | WT PBS vs. WT ASO | 0.9840 |
| WT PBS vs. Tubb4a$^{D249N/KO}$ PBS | **0.0002** | WT PBS vs. Tubb4a$^{D249N/KO}$ PBS | 0.9980 |
| WT PBS vs. Tubb4a$^{D249N/KO}$ ASO | **0.0060** | WT PBS vs. Tubb4a$^{D249N/KO}$ ASO | 0.9266 |
| WT ASO vs. Tubb4a$^{D249N/KO}$ PBS | **<0.0001** | WT ASO vs. Tubb4a$^{D249N/KO}$ PBS | 0.9965 |
| WT ASO vs. Tubb4a$^{D249N/KO}$ ASO | **0.0007** | WT ASO vs. Tubb4a$^{D249N/KO}$ ASO | 0.9936 |
| Supplemental Figure 31B | | **Supplemental Figure 31C** | |
| Histology – NeuN profile/mm$^2$ | **P48** | **Histology – NeuN/caspase3 profile/mm$^2$** | **P48** |
| WT PBS vs. WT ASO | 0.9243 | WT PBS vs. WT ASO | >0.9999 |
| WT PBS vs. Tubb4a$^{D249N/KO}$ PBS | 0.0015 | WT PBS vs. Tubb4a$^{D249N/KO}$ PBS | 0.0206 |
| WT PBS vs. Tubb4a$^{D249N/KO}$ ASO | 0.0011 | WT PBS vs. Tubb4a$^{D249N/KO}$ ASO | 0.0467 |

### Western blot images for Figure 1L and associated Supplemental Figure 4

Full western blots for Figure 1K and Supplemental Figure 4B
P32-P37

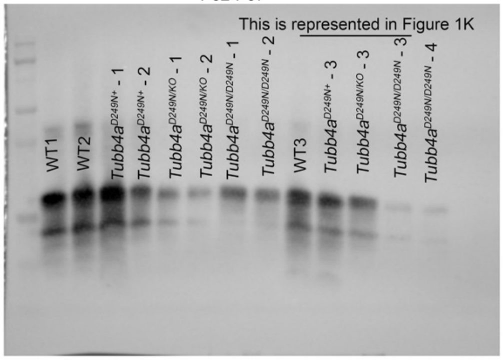

Full western blots for Figure 1K and Supplemental Figure 4B
P108-P110

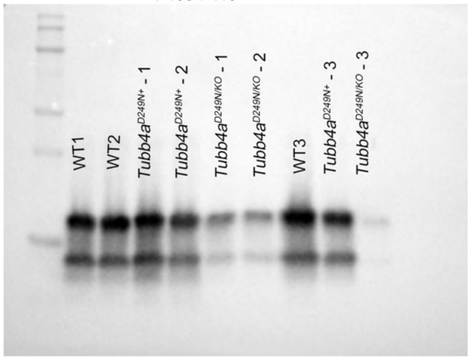

Full western blots for Supplemental Figure 4C

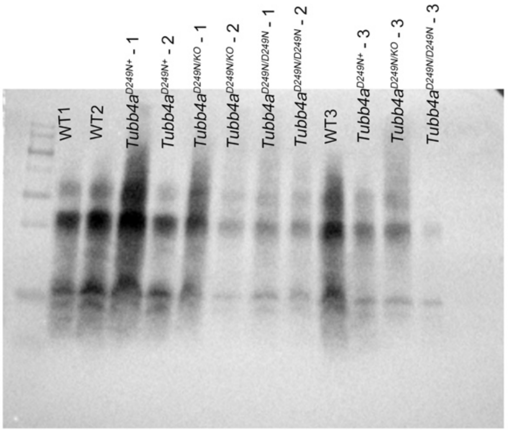

Full western blots for Supplemental Figure 4C

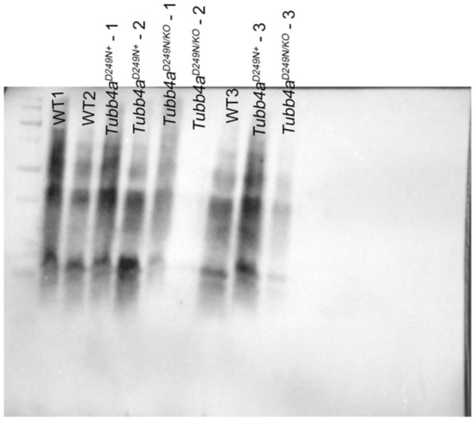

Full western blots for Supplemental Figure 4D

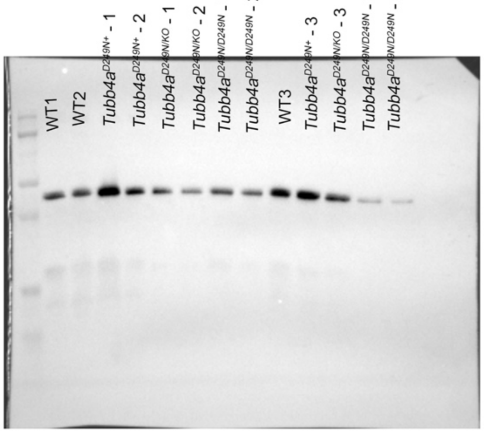

Full western blots for Supplemental Figure 4D

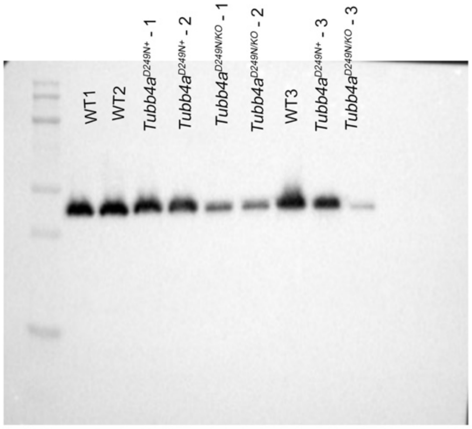

### Western blot images for Figure 5C

| P30 –Loading pattern | | | | | | | | | |
|---|---|---|---|---|---|---|---|---|---|
| 1 | 2 | 3 | 4 | 5 | 6 | 7 | 8 | 9 | 10 |
| Ladder | PBS | PBS | PBS | PBS | ASO18-3 | ASO18-3 | ASO18-3 | ASO18-3 | ASO18-3 |

Western blot of the cortex

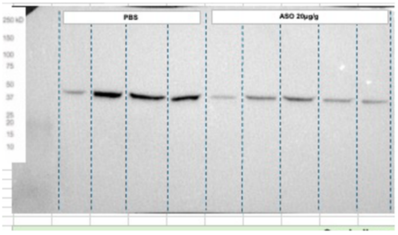

Western blot of the Striatum

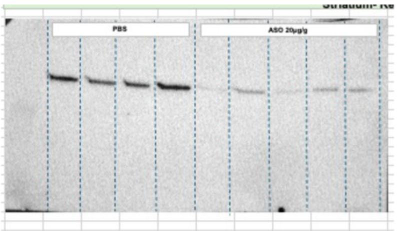

Western blot of the cerebellum

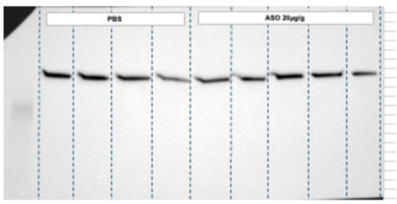

Vinculin (corresponding housekeeping)

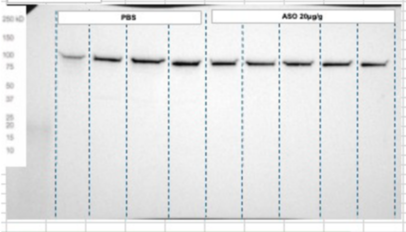

Vinculin (Corresponding housekeeping)

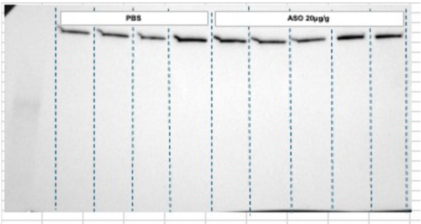

Vinculin (Corresponding housekeeping)

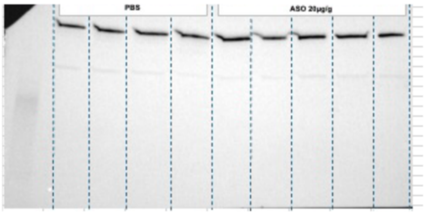

### Western blot images for Figure 5E and 5F

n=3 samples ran in 3 different gels

Loading pattern

| Lane 1 | Lane 2 | Lane 3 | Lane 4 | Lane 5 | Lane 6 | Lane 7 | Lane 8 | Lane 9 |
|---|---|---|---|---|---|---|---|---|
| | P30 | | P60 | | P90 | | P160 | |
| Ladder | WT-PBS | WT-ASO | WT-PBS | WT-ASO | WT-PBS | WT-ASO | WT-PBS | WT-ASO |

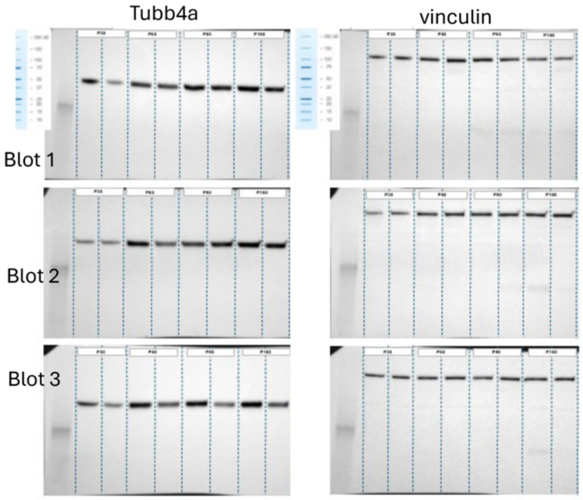

n=3 samples ran in 3 different gels

Loading pattern

| Lane 1 | Lane 2 | Lane 3 | Lane 4 | Lane 5 | Lane 6 | Lane 7 | Lane 8 | Lane 9 |
|---|---|---|---|---|---|---|---|---|
| | P30 | | P60 | | P90 | | P160 | |
| Ladder | WT-PBS | WT-ASO | WT-PBS | WT-ASO | WT-PBS | WT-ASO | WT-PBS | WT-ASO |

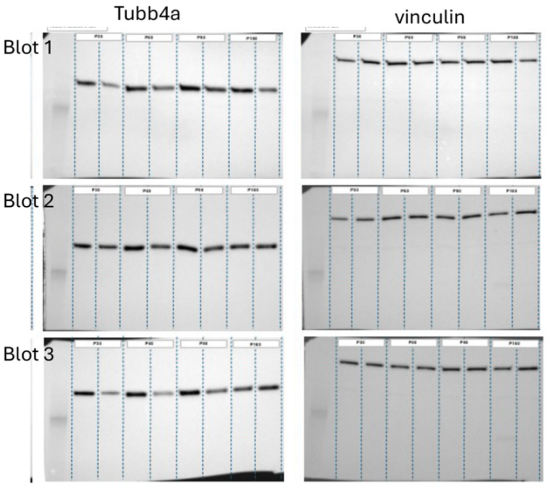

### Western blot images for Supplemental Figure 15B

#### Full western blot for Supplemental Figure 15B

n=3 samples ran in 3 gels

Loading pattern

| Lane 1 | Lane 2 | Lane 3 | Lane 4 | Lane 5 | Lane 6 | Lane 7 | Lane 8 | Lane 9 |
|---|---|---|---|---|---|---|---|---|
| | P30 | | P60 | | P90 | | P160 | |
| Ladder | WT-PBS | WT-ASO | WT-PBS | WT-ASO | WT-PBS | WT-ASO | WT-PBS | WT-ASO |

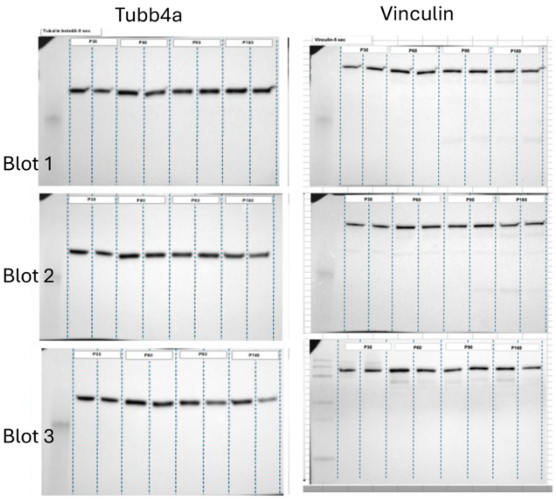

### Western blot images for Supplemental Figure 21E and 21F

Loading pattern

| 1 | 2 | 3 | 4 | 5 | 6 | 7 | 8 | 9 | 10 | 11 | 12 | 13 |
|---|---|---|---|---|---|---|---|---|---|---|---|---|
| Ladder | WT_PBS1 | WT_ASO1 | Tubb4aD249N KO_PBS1 | Tubb4aD249N KO_ASO1 | WT_PBS2 | WT_ASO2 | Tubb4aD249 N KO_PBS 2 | Tubb4aD249N KO-ASO2 | WT_PBS3 | WT_ASO3 | Tubb4aD249 N KO_PBS3 | Tubb4aD249N KO_ASO3 |

#### Full western blot for Supplemental Figure 21E

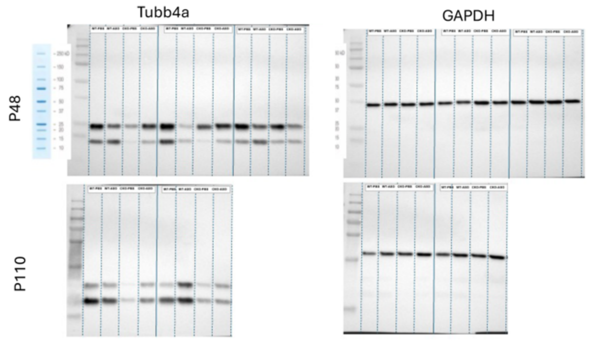

#### Full western blot for Supplemental Figure 21F

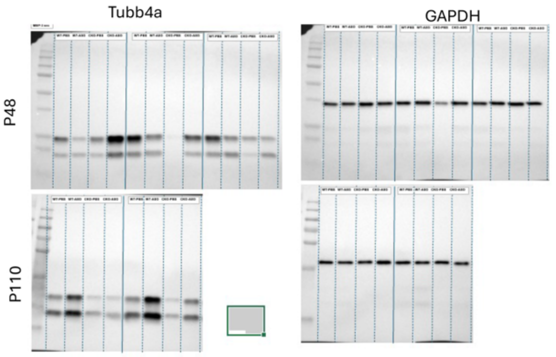
